# Rapid iPSC inclusionopathy models shed light on formation, consequence and molecular subtype of α-synuclein inclusions

**DOI:** 10.1101/2022.11.08.515615

**Authors:** Isabel Lam, Alain Ndayisaba, Amanda J. Lewis, YuHong Fu, Giselle T. Sagredo, Ludovica Zaccagnini, Jackson Sandoe, Ricardo L. Sanz, Aazam Vahdatshoar, Timothy D. Martin, Nader Morshed, Toru Ichihashi, Arati Tripathi, Nagendran Ramalingam, Charlotte Oettgen-Suazo, Theresa Bartels, Max Schäbinger, Erinc Hallacli, Xin Jiang, Amrita Verma, Challana Tea, Zichen Wang, Hiroyuki Hakozaki, Xiao Yu, Kelly Hyles, Chansaem Park, Thorold W. Theunissen, Haoyi Wang, Rudolf Jaenisch, Susan Lindquist, Beth Stevens, Nadia Stefanova, Gregor Wenning, Kelvin C. Luk, Rosario Sanchez Pernaute, Juan Carlos Gómez-Esteban, Daniel Felsky, Yasujiro Kiyota, Nidhi Sahni, S. Stephen Yi, Chee-Yeun Chung, Henning Stahlberg, Isidro Ferrer, Johannes Schöneberg, Stephen J. Elledge, Ulf Dettmer, Glenda M. Halliday, Tim Bartels, Vikram Khurana

## Abstract

Intracellular inclusions accompanying neurodegeneration are histopathologically and ultrastructurally heterogeneous but the significance of this heterogeneity is unclear. iPSC models, while promising for disease modeling, do not form inclusions in a reasonable timeframe and suffer from limited tractability. Here, we developed an iPSC toolbox utilizing piggyBac-based or targeted transgenes to rapidly induce CNS cells with concomitant expression of aggregation-prone proteins. This system is amenable to screening and longitudinal tracking at single-cell and single-inclusion resolution. For proof-of-principle, cortical neuron α-synuclein “inclusionopathy” models were engineered to form inclusions through exogenous seeding or α-synuclein mutation. These models recapitulated known fibril- and lipid-rich inclusion subtypes, uncovering dynamic interactions between them, and refined the classification of inclusions in postmortem brain. Genetic-modifier and protein-interaction screens pinpointed proteins like RhoA whose sequestration into specific inclusion subtypes is likely to be toxic. This iPSC platform should enhance our understanding of proteinaceous pathologies in neurodegeneration and facilitate therapeutics development.

## INTRODUCTION

In neurodegenerative proteinopathies—such as Alzheimer’s disease (AD), Parkinson’s disease (PD) or frontotemporal dementias—aberrant proteinaceous inclusions are widespread within a multitude of neuronal and glial subtypes of the central nervous system (CNS) (Lam et al., 2020). Amyloid fibrils, rich in secondary β-sheet structure, are an important component of these inclusions. Substantial recent evidence points to the importance of amyloid conformers in conferring phenotypic specificity in neurodegenerative disease (Jarosz and Khurana, 2017; Soto and Pritzkow, 2018): different higher-order conformational states, or “strains”, of the same protein can lead to distinct distributions of cellular pathologies and lead ultimately to different diseases (Kaufman et al., 2016; Peng et al., 2018; Perren et al., 2020; Yang et al., 2022a).

Beyond protein conformational heterogeneity, histopathologic analysis reveals a multitude of distinct inclusion types occur in neurodegenerative proteinopathies. For example, Lewy bodies and other α-synuclein (αS)-rich inclusion pathologies have a predilection for neuronal subtypes in brainstem and cortex in synucleinopathies, whereas oligodendroglial cytoplasmic inclusions are pathognomonic in multiple system atrophy (MSA). Tau pathology can manifest as neuronal Pick bodies, tufted astrocytes, globular astrocytic inclusions or oligodendroglial coiled bodies (Chung et al., 2021; Wakabayashi et al., 2013). There is also ultrastructural heterogeneity. Some of this relates to distinct amyloid fiber conformers within inclusions. For example, tau can assemble as straight filaments in progressive supranuclear palsy (PSP), but predominantly assembles into paired helical filaments in AD neurofibrillary tangles (Oakley et al., 2020).

Inclusions can even exhibit different ultrastructural morphology within the same cell type. A striking example is the presence of membrane-rich αS inclusions (so-called “pale bodies”) versus fibril-rich Lewy bodies in PD and other Lewy body diseases. These are best appreciated in substantia nigra dopaminergic neurons, where they are noted even with hematoxylin and eosin staining. A recent study re-affirmed with correlative light and electron microscopy (CLEM) that membrane-predominant inclusion pathologies occur far more frequently than anticipated in PD brain (Shahmoradian et al., 2019). This heterogeneity is frequently overlooked, in part because different inclusions often stain with the same antibodies during neuropathologic examination. For example, both Lewy body and pale body αS inclusions stain avidly for αS phosphorylated at serine 129 (pS129) (Kon et al., 2020)and tau inclusions in different cell types label with the same phospho-, isoform- and conformer-specific antibodies (Ferrer et al., 2014).

Critical and under-appreciated biology may relate to inclusion heterogeneity. It is known that distinct types of inclusions can have very different properties. For example, an aggregating protein can transition from toxic to nontoxic when redirected from juxtanuclear compartments (JUNQ) to insoluble protein deposits (IPODS) (Jarosz and Khurana, 2017; Weisberg et al., 2012). The fate of the cell might thus depend on the functional compartment (JUNQ or IPOD) which the inclusion is sequestered in or associated with.

More generally, the distinct properties of inclusions could explain the disconnect between inclusion formation and neurodegeneration (Halliday, 2015; Libow et al., 2009; Schulz-Schaeffer, 2015) and other conflicting results in the literature. For example, inclusions have variously been described as protective (Arrasate et al., 2004; Chen and Feany, 2005) or detrimental (Mahul-Mellier et al., 2020; Osterberg et al., 2015). Such conflicts speak to the need of a finer-grained classification of inclusion subtypes to better understand their implications for cellular states.

Ideally, inclusion heterogeneity could be tackled in a relevant human cellular model. Induced pluripotent stem-cell (iPSC) models, with the potential to be differentiated into any CNS cell subtype and to model patient-specific pathologies, certainly present an attractive opportunity. However, current human iPSC models have limited tractability, often requiring lengthy differentiation, and mature inclusions do not form in a reasonable timeframe (Iovino et al., 2015; Mazzulli et al., 2016). In models where human iPSC-derived neurons are seeded with aggregates, the heterogeneity of inclusions and how closely they mimic brain pathology has to our knowledge not been sufficiently addressed (Iannielli et al., 2022; Oakley et al., 2020; Tanudjojo et al., 2021). Here, we present a suite of stem cell-based models that enable rapid one-step transdifferentiation from iPSC to CNS cells in conjunction with rapid inclusion formation. The system utilizes rapid, scalable and virus-free overexpression of transdifferentiation factors with Gateway-compatible piggyBac (pB) vectors. This system enables the scalable generation of iPSC-derived CNS cells as readily as with a conventional cell line. Coordinated expression of aggregation-prone proteins is achieved either through all-in-one pB transgenes or transgenes targeted to specific genomic loci. We generated >80 pB-transfected lines that enable rapid directed iPSC-to-CNS trans-differentiation into cortical neurons or astrocytes, and developed an expression system for αS to rapidly induce either lipid-rich inclusions that form spontaneously or fibril-rich inclusions that form with exogenous seeding in induced cortical neurons within ~2 weeks upon induction of transdifferentiation. The system is amenable to longitudinal tracking at single-cell and single-inclusion level, revealing differential effects of specific inclusions on survival. Unexpectedly, longitudinal inclusion tracking, including dynamic lattice-sheet microscopy, reveals biologically impactful interactions between inclusion subtypes within neurons. We show that advanced pathologies found in postmortem brains are recapitulated in this model and our models refine subclassification of inclusions identified in postmortem brain. Moreover, our models enable the identification of novel inclusion subtypes in postmortem brain. Genome-scale CRISPR screens and systematic protein-interaction mapping pinpoint key sequestered proteins, including RhoA, that are likely to be particularly toxic when sequestered.

Our transgenic model sheds light on the dynamic nature of αS inclusions, their potential molecular interactions, and previously unidentified inclusion subtypes of biological relevance. These generalizable inclusionopathy models promise to be useful for biological and drug discovery in neurodegenerative proteinopathies.

## RESULTS

### A robust piggyBac-based expression vector facilitates iPSC transdifferentiation

We aimed to create human CNS proteinopathy models as straightforward to generate and culture as conventional human cell lines, amenable to high-throughput genetic and small-molecule screens and readily transferable between research groups. To do this we (1) leveraged directed transdifferentiation in which overexpression of lineage-specific transcription factors in iPSC/hESC is used to generate rapid, scalable, reproducible and relatively pure populations of distinct CNS cell types (**Figure 1A**, left) (Canals et al., 2018; Zhang et al., 2013); (2) induced inclusion formation rapidly (hence “inclusionopathy” models) through overexpression of the aggregation-prone toxic protein of interest with a variety of methods (**Figure 1A**, right).

**Figure 1:**
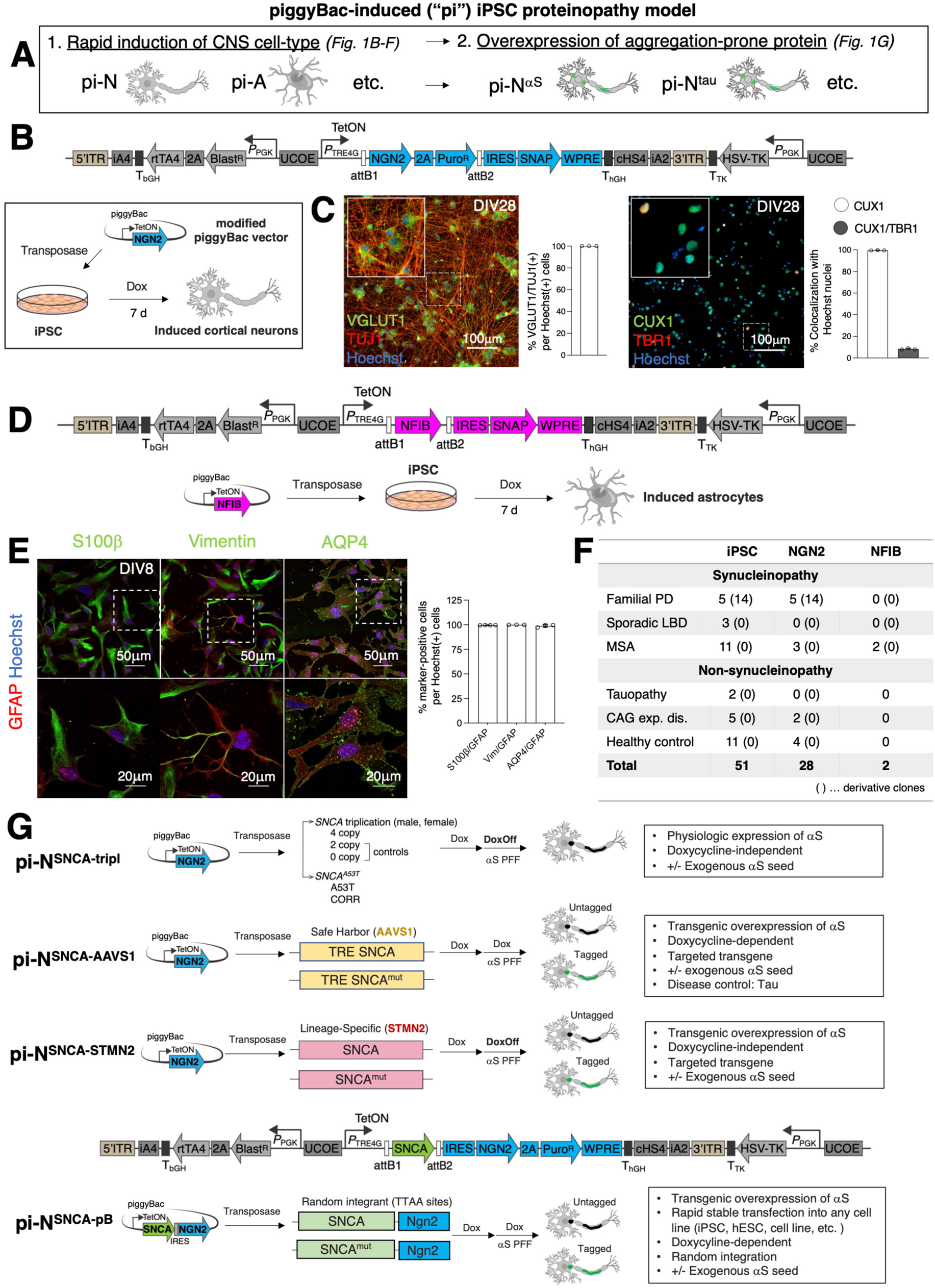
Overview of piggyBac-induced iPSC proteinopathy models. (A) Classification of ***p***iggyBac ***i***nduced (“pi”) iPSC proteinopathy model system. (B) Modified piggyBac (pB) vector containing multiple insulator sequences, antibiotic resistance cassette, and an NGN2-2A-Puro insert regulated by TRE4G inducible promoter, allowing direct transdifferentiation of iPSCs into cortical neurons within 7 days upon treatment with doxycycline. The NGN2 module is followed by an IRES sequence and SNAP tag that can be used as a proxy for transgene expression level. (C) Immunofluorescence (IF) staining and associated quantification of transdifferentiated neurons (pi-N) generated from H9 human embryonic stem cells (hESC) confirm cortical glutamatergic neuron identity (layer II/III) by expression of VGLUT1 (green, left image), TUJ1 (red, left image) and CUX1 (green, right image). TBR1 (red, right image), a general marker for deep layer cortical neurons and a subset of layer II/III, is sporadically expressed in ~10% of neurons in the absence of deep layer marker Ctip2, confirming superficial cortical neuron identity. (D) Modified pB vector with an NFIB insert regulated by TRE4G inducible promoter allowing transdifferentiation of iPSCs into astrocytes within 7 days. (E) Left: IF images of H1 hESC-derived pB-induced astrocytes (pi-A) stained for canonical astrocyte markers GFAP (red), S100β+ (green), Vimentin (green) and AQP4 (green). Nuclei are stained with Hoechst (blue). Right: IF quantification across 3 technical replicates. Each replicate represents 42 image fields (1 image field > 60 cells) acquired from one well. (F) Summary table of iPSC lines introduced with either pB-NGN2 or pB-NFIB. In sum, 51 iPSC lines and derivative clones have been generated from proteinopathy and healthy control cases, of which 28 lines were generated to inducibly overexpress NGN2, and 3 iPSC lines to overexpress NFIB. CAG exp. dis., CAG expansion disease. (G) Overview of proteinopathy platform including either pathologic (but endogenous) expression of the protein of interest, or transgenic overexpression in one of three ways: targeting to a safe harbor locus (*AAVS1*), targeting a lineage-specific locus (*STMN2*), or pB random integration. All lines rely on doxycycline induction of pB-NGN2, but the lines differ in whether doxycycline is needed for proteotoxic transgene overexpression. Physiologic overexpression relies on endogenous *SNCA* promoter, and lineage-specific model relies on the promoter at site of integration, (e.g., *STMN2*), so these models are doxycycline-independent for toxic protein expression. The toxic protein is under doxycycline-inducible promoter for the safe harbor (*AAVS1*) and pB all-in-one random integrant models, so these models are doxycycline-dependent for toxic protein expression.

To target the transcription factor-encoding transgenes, we decided on the pB delivery system. pB is a DNA transposon that specifically integrates at TTAA sites in the genome and has been widely exploited as a tool for transgenesis and genome engineering(Woodard and Wilson, 2015). The pB system offers a virus-free and scalable approach with large cargo size for genomic integration of desired transgenes in iPSCs with a simple transfection protocol. The system avoids the expense and batch-to-batch variability associated with virus transduction. Recently, we incorporated key modifications to the pB integrating vector to ensure stable and high expression levels of the transgene cargo(Hallacli et al., 2022) and we utilized a similar vector backbone here. We increased versatility by utilizing the Gateway cloning system, enabling introduction of different transgenes through straightforward cloning reactions.

To establish proof-of-principle, we selected CNS cell types for which robust methods in lentiviral vectors have been published—NGN2 for superficial layer II/III cortical glutamatergic neurons (**Figure 1B**) (Zhang et al., 2013) and NFIB for astrocytes (**Figure 1D**) (Canals et al., 2018). As with the lentiviral protocols, the pB approach creates robust cell products. hESC-derived neurons with pB Ngn2 (pB-NGN2) exhibited appropriate markers for glutamatergic cortical neuron fate (Vglut1 and Tuj1/β-tubulin III) and, more specifically, for superficial (Cux1; layer II/III) but not deep (Tbr1; layer V) cortical layers (**Figure 1C**). pB-NFIB expression in hESC resulted in appropriate cell type-specific markers–GFAP, S100β, vimentin, and AQP4 (**Figures 1E; S1A-S1B**). We henceforth refer to these differentiated cells as piggyBac-induced neuron (pi-N) and piggyBac-induced astrocyte (pi-A). We have assembled in our laboratory approximately 100 pluripotent stem cell lines, from hESCs to hiPSCs, across a number of different neurodegenerative diseases (a selection is outlined in **Figures 1F**, **S1K**) (**Table S1**). To date, we have successfully introduced pB-NGN2 into 56 iPSC lines and derivative clones and pB-NFIB into 2 iPSC lines (**Figure 1F, Table S1**).

### Rapid pB iPSC proteinopathy (“inclusionopathy”) models are established in conjunction with overexpression of aggregation-prone proteins

To investigate inclusions in rapid timeframes amenable to biological and drug discovery, we developed multiple complementary methods for expression of aggregation-prone protein, each with distinct advantages. We used αS as an initial prototype though have generated tau, β-amyloid and ApoE constructs in our lab also (**Figure 1G; Table S1;** not shown). In a pathologic overexpression system that expresses αS endogenously, pB-NGN2 is integrated in iPSCs reprogrammed from female (Hallacli et al., 2022) and male Iowa kindred patients with an *SNCA* gene triplication. Isogenic allelic series were generated through CRISPR/Cas9 engineering (*SNCA* “4-copy” parental, wild-type “2-copy” knockdown, and null “0-copy” knockout) (**Figures 1G**, top and **2A**, left). We have also introduced pB-NGN2 into iPSC generated from a patient harboring the *SNCA* A53>T mutation (“A53T”) or its isogenic mutation-corrected control line (“CORR”)(Hallacli et al., 2022) (**Figures 1G**, top and **2A**, right). In these models, doxycycline is required for Ngn2 expression, but is withdrawn at 1 week (i.e., once neurons have differentiated).

**Figure 2:**
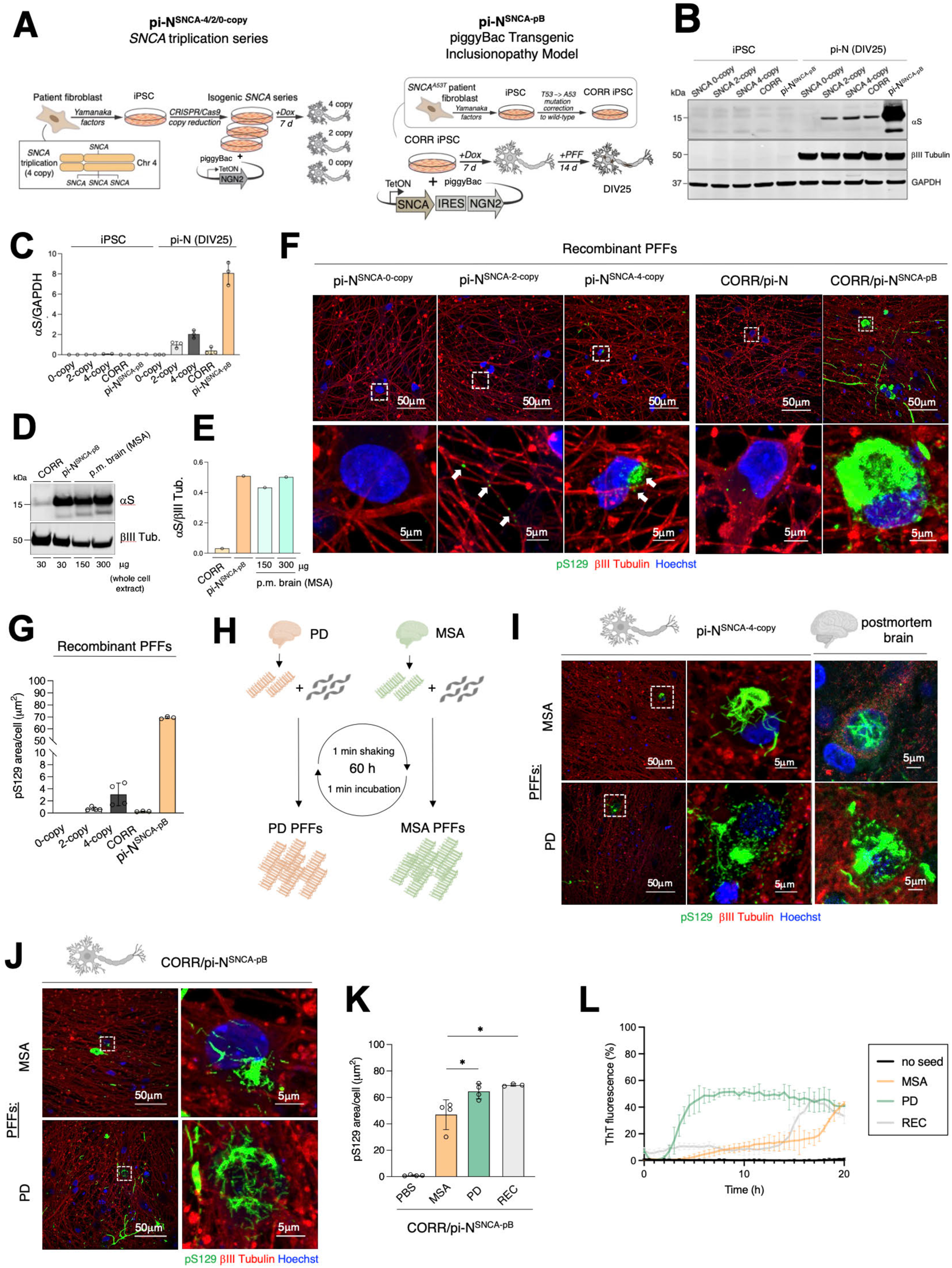
Induction of αS inclusions through amyloid seeds is enhanced by piggyBac-based overexpression of αS. (A) Schematic diagrams of pathologic overexpression (*SNCA* 4-copy) (left) and pB transgenic (right) proteinopathy models. Left: Schematic showing the generation of isogenic human neurons with different *SNCA* copy numbers from PD patient fibroblasts with *SNCA* locus triplication (4-copy). The *SNCA* triplication fibroblasts were reprogrammed to iPSC and subsequently engineered with CRISPR/Cas9 to knock-out *SNCA*, resulting in isogenic 4-copy, 2-copy, and 0-copy lines (pi-N^SNCA-4/2/0-copy^). Integration of pB-NGN2 into these iPSC lines allows direct differentiation to neurons. Right: Schematic showing the generation of a mutation-corrected line (herein referred to as ‘CORR’) derived from an A53T familial PD patient (inset). The CORR iPSC line was subsequently engineered to harbor the all-in-one pB-SNCA-IRES-NGN2 plasmid to allow doxycycline-inducible overexpression of untagged wild-type αS (pi-N^SNCA-pB^). (B) Western blot analysis in *SNCA* 0/2/4-copy, CORR and isogenic CORR/pi-N^SNCA-pB^ iPSCs and neurons reveals substantially higher αS steady-state protein levels in pi-N^SNCA-pB^ neurons. (C) Quantification of αS levels normalized to GAPDH from western blot in panel B across 3 replicates. (D) Western blot analysis in CORR and isogenic CORR/pi-N^SNCA-pB^ neurons versus MSA postmortem brain lysate (frontal cortex) reveals comparable αS steady-state protein levels in pi-N^SNCA-pB^ neurons and MSA postmortem brain upon normalization to neuron-specific marker bIII Tubulin. (E) Quantification of αS levels normalized to bIII Tubulin from western blot in panel D. (F) Representative confocal images of αS pS129 IF in PFF-seeded cortical neurons demonstrate significantly increased seeding efficiency in pi-N^SNCA-pB^ neurons upon PFF exposure compared to pi-N^SNCA-4/2/0-copy^ and CORR/pi-N. (G) Quantification of pS129 area from panel D across 3 replicates. Each replicate represents 42 image fields (1 image field > 100 neurons) acquired from one well. (H) Schematic illustration of seeded amplification assay (SAA) to capture and amplify αS fibrils from MSA and PD postmortem brain. Sarkosyl-insoluble fractions of brain homogenates are subjected to alternating 1 min shaking/1 min incubation periods for 60 h in the presence of recombinant monomeric αS to template and amplify MSA and PD αS fibrils. (I) IF of pS129 in pi-N^SNCA-4-copy^ neurons (left and center) seeded with MSA and PD PFFs reveals inclusion morphologies reminiscent of post-mortem brain inclusions (right) for each respective disease (MSA: perinuclear ‘skein’-like inclusion, PD: diffuse fibrillar inclusion). (J) IF of pS129 in transgenic pi-N^SNCA-pB^ neurons seeded with SAA-generated αS fibrils derived from MSA or PD postmortem brain shows similar inclusion morphologies compared to pi-N^SNCA-4-copy^ neurons, but with higher seeding efficiency. (K) Quantification of panel H revealed decreased seeding efficiency for MSA PFFs (n=1) compared to PD PFFs (n=1) and recombinant WT PFFs. Each replicate represents 42 image fields (1 image field > 100 neurons) acquired from one well. (L) SAA re-amplification of CORR/pi-N^SNCA-pB^ neuronal lysates previously seeded with MSA and PD PFFs for 14 days showed an accelerated aggregation propensity of PD PFFs (n=1) compared to MSA PFFs (n=1). (M) Quantification of lag time in SAA assay from panel G confirms distinct aggregation kinetics of MSA PFF strain compared to PD PFFs and recombinant WT PFFs. One-way ANOVA + Tukey’s multiple comparison test for panels I and K: * p<0.05, *** p<0.001.

Ngn2 can also be coupled with transgenic overexpression. In the past, the *AAVS1* (also known as *PPP1R12C*) safe-harbor locus has been targeted for transgenic expression (Hockemeyer et al., 2011). We utilized a similar system, targeting rtTA transactivator (under pCAGGS promoter) for tetracycline inducible expression to one *AAVS1* allele and different αS or tau transgenic constructs under a tetracycline response element (TRE) to the other allele (**Figures 1G**, second from top and **S1C-S1D**). Thus, the gene encoding the toxic protein (*SNCA* or *MAPT* etc.) is under doxycycline-inducible control of the rtTA transactivator (Tet-On). Targeting at the *AAVS1* locus was verified by Southern blot (**Figure S1D**), and expression of the transgene was confirmed to be doxycycline dose-dependent (**Figure S1C**, inset). Theoretically, even without pB transdifferentiation, this method should enable differentiation into any cell type and doxycycline-inducible expression in those cells. In practice, we have found the system is limited by silencing of the TRE-driven transgene (**Figure S1E** and data not shown) and poor expression in certain cell types, including astrocytes (**Figure S1F**). Introduction of the pB, shown in **Figures S1E** (right), **S1G-S1I** for Ngn2, addressed this issue with αS transgene expression in more than 90% of pi-Ns at DIV25. pi-N^SNCA-mK2-AAVS1^ transdifferentiated with pB-NGN2 exhibited markers for glutamatergic cortical neuron fate (Vglut1, Tuj1) and superficial (Cux1; layer II/III) but not deep (Tbr1; layer V) cortical layers (**Figures S1H-S1I**). The advantage of this system is that the αS transgene is targeted to a defined region in single copy when appropriately verified. The disadvantage is that αS transgene expression is doxycycline dependent.

To show proof-of-principle that a similar system could be adapted for lineage-specific expression in the absence of doxycycline, we targeted *STMN2*, a gene with highly selective expression in neurons (Brain RNA-Seq, brainrnaseq.org)(Zhang et al., 2016) (**Figures 1G**, third from top and **S1J**, left). A construct with internal ribosome entry site (IRES) sequence followed by SNCA-GFP (IRES-SNCA-GFP) flanked by *STMN2* homology arms was targeted to the *STMN2* locus in H9 hESC by CRISPR/Cas9 (**Figure 3H**). Knock-in of *SNCA* transgene at *STMN2* locus does not lead to reduction in *STMN2* expression (**Figure S1J**, right). The advantage here is that *SNCA* expression is under control of the neuron-specific *STMN2* promoter, rendering the system doxycycline-independent. Doxycycline is only required to induce pB-NGN2 expression. As for the *AAVS1*-coupled system, coupling this system to Ngn2 enables transgene expression in more than 90% of pi-Ns at DIV25 (**Figure S1G**).

**Figure 3:**
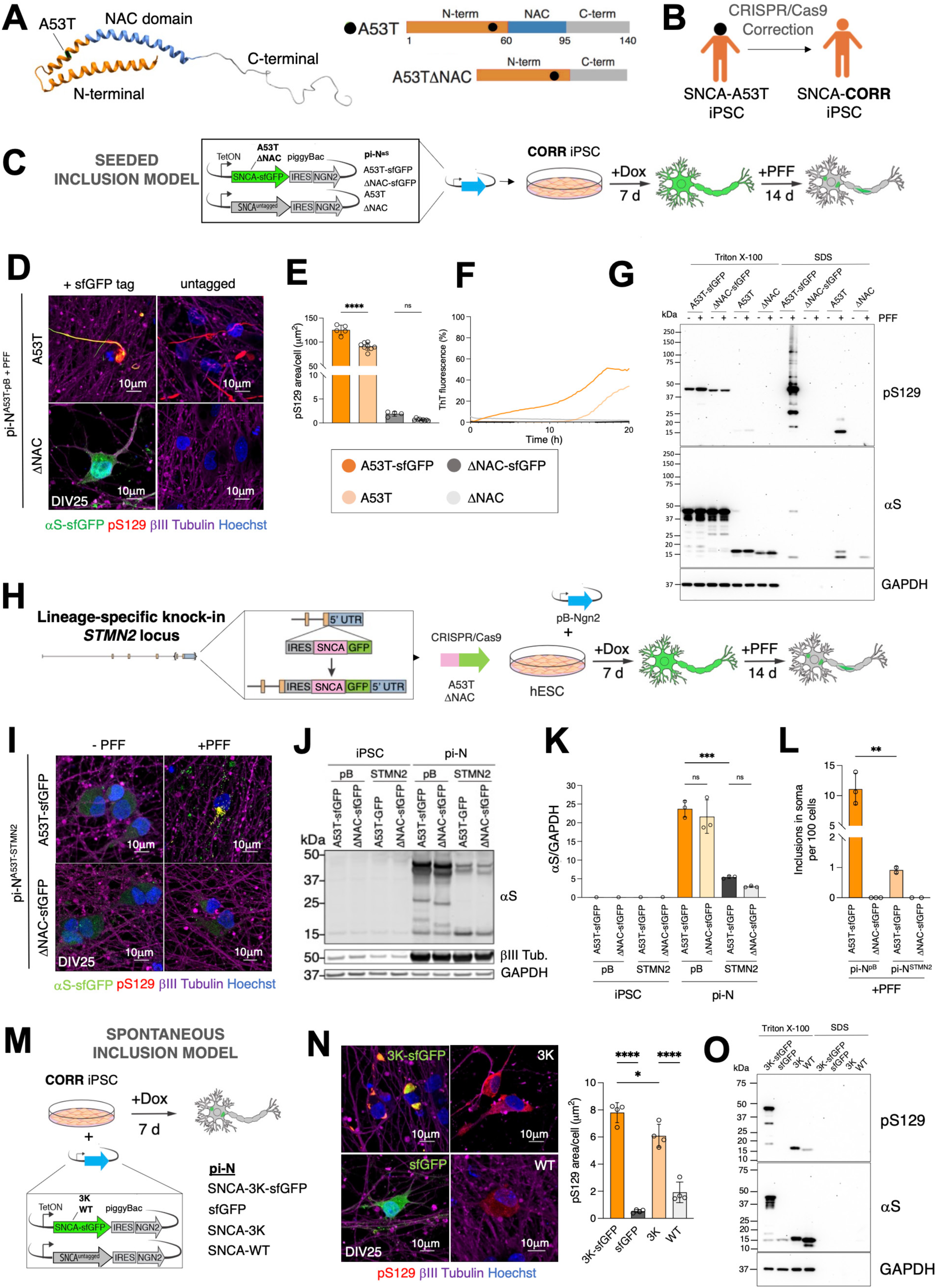
Biochemically distinct pS129-positive αS inclusions in exogenous seed versus spontaneous aggregation model. (A) Protein structure of αS (PDB: 1xq8) juxtaposed with linear maps of full length αS-A53T and seeding-incompetent αS-A53T-ΔNAC indicating relevant amino acid positions. (B) Cartoon depicting the derivation of CORR iPSC line from patient SNCA-A53T iPSC mutation-corrected to SNCA-WT. (C) Schematic outline of seeded inclusionopathy model. CORR iPSCs were transfected with distinct all-in-one pB constructs to simultaneously overexpress NGN2 and sfGFP-tagged or untagged αS-A53T or aggregation-dead αS-A53T-ΔNAC (ΔNAC) upon induction with doxycycline. Cortical neurons were generated within 7 days, and on DIV11, neurons were seeded with PFFs for 14 days to induce rapid and robust inclusion formation. (D) pS129 IF staining in PFF-seeded transgenic cortical neurons overexpressing sfGFP-tagged or untagged αS-A53T (pi-N^A53T-sfGFP-pB+PFF^, pi-N^A53T-pB+PFF^) (A53T) or αS-ΔNAC (ΔNAC) (pi-N^A53T-ΔNAC-sfGFP-pB+PFF^, pi-N^A53T-ΔNAC-pB+PFF^) confirms that αS aggregation is NAC-domain dependent, pS129 signal colocalizes with sfGFP signal, and is morphologically comparable to its untagged counterpart. (E) Quantification of pS129 IF in panel D showed higher seeding efficiency in pi-N^A53T-sfGFP-pB+PFF^ neurons compared to untagged pi-N^A53T-pB+PFF^ neurons. (F) Re-amplification of insoluble αS in PFF-seeded transgenic neurons via SAA showed similar aggregation kinetics for sfGFP-tagged and untagged αS species, whereas no aggregates were amplified in the aggregation-dead (ΔNAC) counterparts. (G) Western blot for total αS and pS129 after sequential Triton X-100/SDS extraction of soluble and insoluble protein fractions confirmed the presence of SDS-insoluble species in both sfGFP-tagged and untagged pi-N^A53T-pB^ neurons. (H) Schematic outline of transgenic αS overexpression driven by *STMN2* promoter. (I) IF for pS129 in PFF-seeded vs unseeded cortical neurons overexpressing sfGFP-tagged αS-A53T (pi-N^A53T-sfGFP-STMN2^) or αS-ΔNAC (pi-N^ΔNAC-sfGFP-STMN2^) leads to formation of punctate neuritic and cell body aggregates in the seeded pi-N^A53T-sfGFP-STMN2+PFF^ model. (J) Western blot analysis showed higher αS steady state protein levels in the pB random integration model compared to *STMN2* lineage-specific targeting in induced cortical neurons. (K) Quantification of panel J across 3 replicates. (L) Quantification of pS129-positive cell soma inclusions in PFF-seeded pB random integration or *STMN2* lineage-specific transgenic neurons showed higher frequency of cell soma inclusions in pi-N^pB^ model compared to pi-N^STMN2^ neurons. (M) Schematic outline of spontaneous inclusionopathy model. PiggyBac vectors with WT or SNCA-3K (sfGFP-tagged or untagged) or sfGFP alone were integrated in CORR iPSCs. (N) Left: IF for pS129 in transgenic cortical neurons overexpressing sfGFP-tagged or untagged αS-3K, untagged αS (WT), or sfGFP control demonstrates spontaneous formation of pS129+ inclusions in neurons overexpressing tagged and untagged αS-3K. Right: Quantification of left panel confirmed higher pS129 burden in pi-N^3K-sfGFP-pB^ and pi-N^3K-pB^ neurons compared to WT overexpression and sfGFP control. (O) Western blot for total αS and pS129 after sequential Triton X-100/SDS extraction of soluble and insoluble protein fractions revealed that spontaneous inclusion models form Triton-X soluble inclusions. One-way ANOVA + Tukey’s multiple comparison test for panels K, L and O: * p<0.05, ** p<0.01, *** p<0.001, **** p<0.0001.

Finally, to avoid the need to target lines each time, we opted for an all-in-one pB plasmid whereby the transgene expressing the aggregation-prone protein lies within the same pB construct as the transcription factor driving differentiation. The protein-encoding transgene is introduced easily through Gateway cloning. The transdifferentiation factor is expressed under the same dox-inducible promoter but separated by an IRES sequence, thus allowing simultaneous transdifferentiation into cortical neurons and overexpression of the aggregation-prone protein. We have thoroughly explored this system for αS inclusion formation (**Figure 1G**, lowest panel). This proved to be the most straightforward and efficient system for studying formation of intraneuronal αS inclusions and their consequences, but drawbacks include doxycycline-dependence and random integration of transgenes.

### Induction of aS inclusions through amyloid seeds is enhanced by pB-based CCC αS

Exogenous exposure of neurons to pre-formed fibrils (PFFs) of αS, in which endogenous αS is templated into insoluble amyloid conformers and deposited into intracellular inclusions, has become a standard way to accelerate αS pathology both in primary neurons(Volpicelli-Daley et al., 2014, 2011) and in rodent models(Kim et al., 2019; Luk et al., 2012). We exposed pi-Ns to PFFs and conducted a head-to-head comparison of the non-transgenic pathologic overexpression (*SNCA* triplication series) versus pB transgenic models. For the latter, we utilized an iPSC line generated from an A53T patient that had been genetically corrected with CRISPR/Cas9-mediated editing (“CORR” iPSC line) to create a line on a “synucleinopathy-permissive” genetic background(Hallacli et al., 2022) (**Figure 2A**, right). In that line we introduced our all-in-one pB expressing wild-type αS, henceforth “pi-N^SNCA-pB^” (**Figure 1G**, lowermost). This line was compared to the triplication line allelic series in which pB-NGN2 alone had been integrated (**Figure 2A**, left; **Figure 1G**, uppermost; henceforth, “pi-N^SNCA-4/2/0-copy^”). Both overexpression models transdifferentiated with pB-NGN2 exhibited the appropriate markers for glutamatergic cortical neuron fate and superficial cortical layers (**Figures S2A-S2B**). pi-N^SNCA-pB^ exhibits substantially higher αS steady-state protein level than pi-N^SNCA-4-copy^ (**Figures 2B-2C**). Importantly, despite this overexpression, actual αS levels generated in our pB model are closely matched to the brain, when normalized for the neuronal marker b-tubulin III (**Figures 2D-2E**). This reflects the low levels of αS expression in iPSC-derived neurons.

We triggered inclusion formation by exposing pi-Ns (11 days after Ngn2 induction) to bath-sonicated recombinant αS pre-formed fibrils (PFFs) (**Figure S2C**). Inclusions staining positive for pS129 were detected in pi-N^SNCA-4-copy^, but inclusion deposition was far more robust in pi-N^SNCA-pB^ neurons, both within neurites and soma (**Figures 2F-2G**). As expected, no pS129 signal was detected in isogenic pi-N^SNCA-0-copy^ neurons, and very few inclusions were detected in PFF-seeded neurons expressing wild-type levels of αS (pi-N^SNCA-2-copy^ or CORR/pi-N).

Recently, there has been interest in amplifying fibrillar material from postmortem proteinopathy brain. We were keen to demonstrate this workflow is also compatible with our pi-N models. We generated PFFs from synucleinopathy brain through a seeded amplification assay (SAA) in which brain lysates are co-incubated with recombinant αS, resulting in templating of the amyloid conformers onto the recombinant protein, hence “seeded amplification” (Shahnawaz et al., 2020; Soto et al., 2002) (**Figure 2H**). For proof-of-principle, we selected one index MSA and one index PD case. We confirmed that the resultant αS fibrils (**Figure S2D**) were comparable to the insoluble fraction of the matching brain lysate by observing similar banding patterns on western blot following proteinase-K digestion (**Figure S2E**). While our intent here was not to compare PD and MSA seeding systematically, seeding pi-N^SNCA-4-copy^ neurons with brain-derived PFFs amplified from these MSA and PD cases resulted in inclusions reminiscent of characteristic inclusion morphologies detected in postmortem brain (MSA: ‘skein-like’ perinuclear neuronal inclusion; PD: diffuse intraneuronal inclusion) (Moors et al., 2021) (**Figure 2I**). Similar inclusions, albeit far more abundant, were induced in the pi-N^SNCA-pB^ neuronal model (**Figure 2J**).

While we can make no claims as to the generalizability of results, we technically could ask whether seeding (as measured by pS129) was comparable to amplification kinetics as measured by the Thioflavin T (ThT) assay which monitors amyloid fibril formation by the binding of fluorescent ThT to amyloids. pS129 seeding in neurons was efficient in PD, MSA and recombinant PFFs. MSA PFFs from this particular case demonstrated slightly lower seeding efficiency in our cortical neuron model (**Figure 2K**). When we re-amplified αS from pi-N^SNCA-pB^ neurons (pi-N^SNCA-pB^ or CORR/pi-N) previously seeded with recombinant, PD or MSA PFFs with SAA, we noted an increased lag time for MSA and recombinant PFFs compared to PD PFFs (**Figures 2L**, **S2F**). Thus, seeding efficiency measurement with pS129 was internally consistent with ThT measurements from matched strains in our neuronal models.

Taken together, our results demonstrate that the pB transgenic overexpression system boosts efficiency of inclusion formation, that inclusion formation can be achieved with both synthetic and brain-derived amyloid seed and that pS129 in these seeded models is proportional to amplification of seed with ThT.

### Biochemically distinct pS129-positive aS inclusions form in exogenous seed versus spontaneous aggregation model

Mutations in αS can alter its propensity to aggregate and bind membranes(Dettmer, 2018; Flagmeier et al., 2016). We explored mutations with our pi-N system, and also compared superfolder GFP (sfGFP)(Pédelacq et al., 2006) tagged and untagged forms of αS, with a view to testing a system for live imaging. sfGFP is a large tag (26.8 kDa) relative to αS (14 kDa) that could theoretically alter propensity to aggregate, although recent studies have suggested αS fibrils form similarly in the presence or absence of a GFP tag(Trinkaus et al., 2021), and known native αS protein interactions were conserved when αS was tagged with a large APEX2 tag (28 kDa)(Chung et al., 2017). We started with the A53T familial αS point mutation that increases the protein’s propensity to aggregate(Flagmeier et al., 2016). As a control, we used *SNCA* with a deletion in the nonamyloid component domain (ΔNAC), an aggregation-dead mutant(Giasson et al., 2001) (**Figure 3A**). We expressed these within our all-in-one pB construct (**Figure 1G**, lowest) in the A53T mutation-corrected (“CORR”) iPSC line (**Figure 3B**). Transgenes were either tagged with sfGFP or untagged.

At first pass, we did not appreciate obvious spontaneous inclusion formation in these transgenic pi-Ns. We thus induced inclusion formation with exogenous recombinant αS PFFs. Seeding with recombinant A53T αS PFFs for 14 days (from DIV11 through DIV25) triggered inclusion formation in A53T-overexpressing pi-Ns but not in the ΔNAC overexpressing counterparts, as indicated through pS129 immunostaining (**Figures 3C-3E**). pS129 signal colocalized with sfGFP signal in the A53T-sfGFP line (**Figure 3D**). Re-amplification of the neuronal lysates via SAA resulted in seeded amplification within 12 h for both sfGFP-tagged and untagged pi-N^A53T-pB+PFF^ models, whereas their respective ΔNAC controls did not induce seeding (**Figure 3F**). These pi-N αS inclusionopathy models were subjected to sequential extraction in Triton X-100 (TX-100) followed by SDS, followed by immunoblotting for pS129 and total αS(Volpicelli-Daley et al., 2014). In the absence of PFF treatment, αS was only detected in the TX-100-soluble fraction (**Figure 3G**). Upon treatment with PFFs, a TX-100-insoluble (SDS-soluble) fraction appeared in pi-N^A53T-pB+PFF^, but not in pi-N^ΔNAC-pB+PFF^ neurons, whether or not the αS was tagged, highlighting the pivotal role of the αS-NAC domain. C-terminal tagging of αS-A53T with sfGFP modestly increased inclusion formation compared to untagged αS – indicated by overall pS129 immunostaining, SAA and TX-100/SDS sequential extraction – but behaviors were overall quite comparable (**Figures 3D-3G**). In our initial characterization of these lines, we asked whether the formation of inclusions in pi-N^A53T-pB+PFF^ cultures was associated with cellular pathologies previously tied to synucleinopathy including mitochondrial respiration, mitochondrial subunit expression and lysosomal flux(Compagnoni et al., 2018; Devi et al., 2008; Hsu et al., 2000; Ludtmann et al., 2018). We did not find any clear perturbation of these pathways upon PFF exposure within the timeframes assayed (**Figures S3A-S3E**).

We also compared the all-in-one pB A53T and ΔNAC neurons to equivalent transgenes knocked in with single copy into the neuronal lineage-specific *STMN2* locus (**Figures 1G**, third from top and **3H**). *SNCA* transgene knock-in at *STMN2* locus results in neuron-specific expression indicated by co-localization with MAP2 (**Figure S3F**), but not with astroglial markers (data not shown). Transdifferentiation of pi-N^A53T-sfGFP-STMN2^ into cortical neurons with pB-NGN2 gave rise to the appropriate markers for glutamatergic cortical neuron fate and superficial cortical layers (**Figures S3G-S3H**). As with the pi-N^SNCA-pB^ neurons, pi-N^STMN2^ neurons expressing A53T develop pS129(+) inclusions when seeded with PFFs, whereas neurons expressing ΔNAC do not (**Figure 3I**). As expected, expression levels were appreciably higher in the pi-N^SNCA-pB^ lines compared to pi-N^SNCA-STMN2^ (**Figures 3J-3K**). Concordant with this, more pS129+ inclusions are detected in pi-Ns overexpressing A53T through pB integration than *STMN2* knock-in (**Figures 3L** and **S3I**). While the *STMN2* and triplication pi-N neurons offer lineage specificity and the advantage of doxycycline-independence, we chose not to proceed with these lines for further analysis in this study because of the relatively low αS seeding propensity. We used the pi-N^A53T-sfGFP-pB^ for further investigation of seeded inclusion formation and consequence in this study.

Finally, we generated an αS construct for an orthogonal pi-N^pB^ model in which inclusionopathy occurs spontaneously through non-amyloid pathway and without need for exogenous αS PFFs. In the so-called “SNCA-3K” mutation, αS membrane-binding is enhanced by amplifying the E46>K familial PD mutation within three imperfect repeats of 11 aa motifs in the N-terminal helix of αS (“3K” refers to E35>K+E46>K+E61>K)(Dettmer et al., 2017; Hallacli et al., 2022). Expression of this mutant with lentiviral vectors in neural cells induces inclusions rich in vesicles and membranous structures(Dettmer et al., 2017; Ericsson et al., 2021). Moreover, a transgenic 3K mouse model recapitulates PD cellular pathologies and clinical manifestations such as levodopa-responsive tremor(Nuber et al., 2018), these 3K models develop cytoplasmic inclusions spontaneously, without needing to seed with exogenous PFFs, offering an entirely orthogonal way to induce aS inclusions. Contrary to the seeded A53T model, inclusion formation in the 3K model is independent of the NAC domain (**Figure S9A**, right). We created a 3K αS pi-N model with our all-in-one pB in the CORR iPSC background (“pi-N^3K-pB^”) (**Figure 3M**). Inclusions in pi-N^3K-pB^ formed spontaneously and stained positive for pS129, in both sfGFP-tagged and untagged models (**Figure 3N**). Sequential TX-100/SDS detergent extraction from pi-N^3K-pB^ and control neurons confirmed that αS-3K is present only in the TX-100 soluble fraction, indicating its absence from large insoluble aggregates (**Figure 3O**). Thus, pi-N^3K-pB^ neurons represent a spontaneous inclusionopathy model, in contrast to the pi-N^A53T-pB^ seeded inclusion model. Like the pi-N^A53T-pB^ model, abundant pS129 inclusions form, but in contrast to the pi-N^A53T-pB^ model, inclusions all appear to be soluble in TX-100. Intrigued by the ability of these two iPSC models to form pS129 inclusions with very different biophysical properties, we investigated these dichotomous models in greater detail.

### Inclusionopathy models are amenable to longitudinal single-cell tracking at the cellular and inclusion level

The impact of αS aggregation pathology on neuronal death or survival is still unclear. This may relate to heterogeneous inclusion biology. In longitudinal cell-tracking assays, the consequence of inclusion formation on cell survival and other cellular pathologies can be assessed (Linsley et al., 2019). Prior longitudinal surveys of cell survival have been highly informative for the field. However, most studies have also tended to focus on binary “with inclusion” or “without inclusion” classifiers. We assessed whether our model systems could discern effects of distinct inclusion subtypes.

We developed an algorithm for single-cell inclusion survival tracking with longitudinal imaging (Biostation CT, Nikon) (**Figure 4A; S4D**). The algorithm inputs time-lapse images of fluorescently-labeled neuronal cultures and automates detection of neurons and inclusions based on fluorescence intensity and size (**Figure S5A**). Next, the algorithm automatically detects neuron live/dead status by evaluating changes in cell area and mean fluorescence intensity between frames (**Figure S4A**). For each neuron that is successfully tracked, the algorithm provides its inclusion status, inclusion size if present, and the timeframe of death (**Figures 4A**, **S4C**). The single-cell inclusion survival tracking is based only on inclusions in the soma, not inclusions in neurites, since it is not always possible to link a neuritic inclusion to its corresponding soma and thus determine the viability status of the neuron. Instead, total neuritic inclusion length is analyzed across cell populations (**Figure S5C**). The accuracy of automated detection and survival status was evaluated for algorithms optimized to the seeded inclusion model and the spontaneous inclusion model (**Figure S4B**).

**Figure 4:**
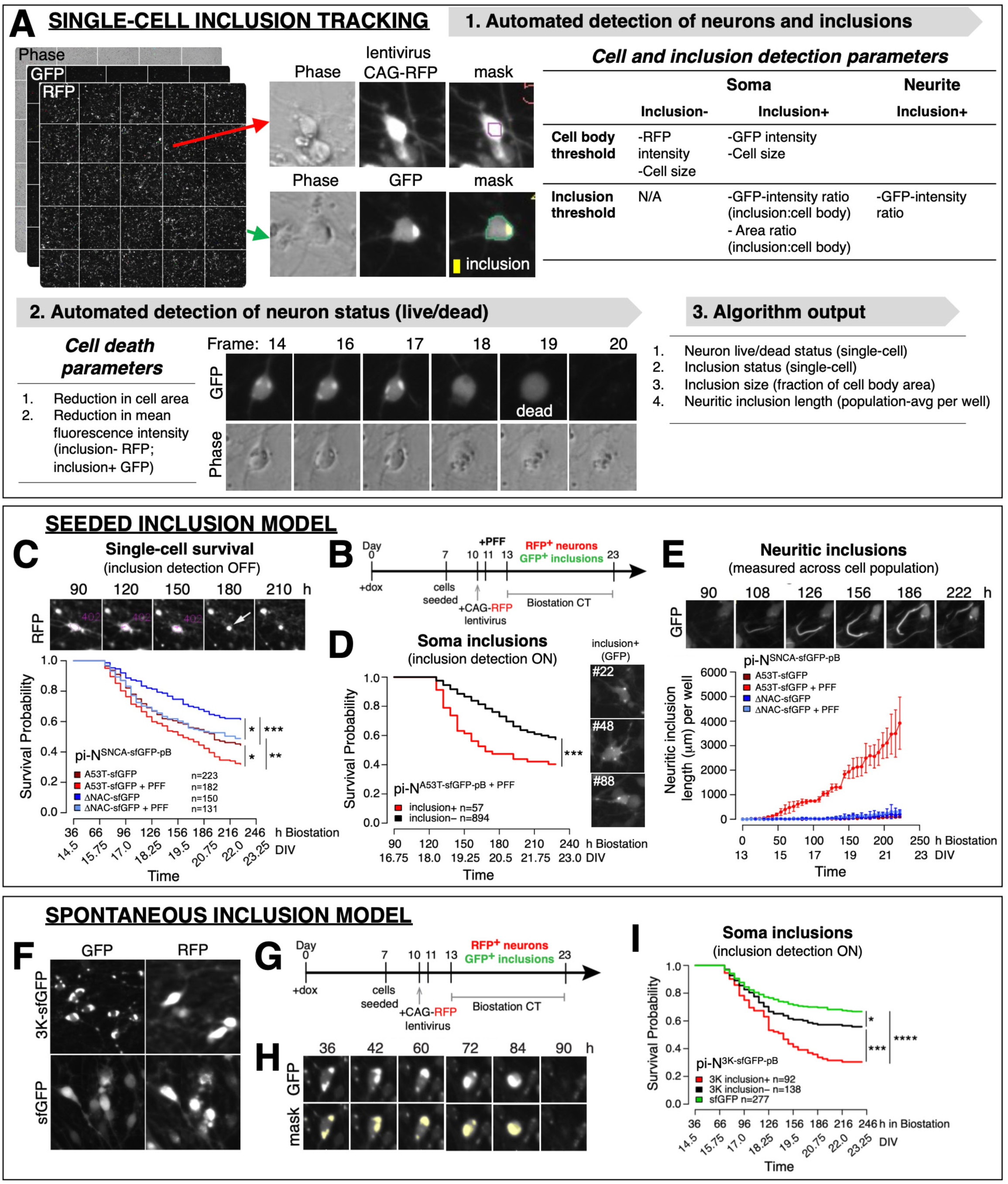
PiggyBac-induced proteinopathy models are amenable to longitudinal quantitative single-cell tracking at the cellular, subcellular and inclusion level. (A) Single-cell inclusion tracking algorithm consisting of automated detection and tracking of neurons and inclusions, and automated detection of neuron live/dead status. Individual wells were captured with 5×5 stitched images with phase, RFP and GFP channels. 1. Automated neuron detection was based on fluorescence intensity and cell size. Soma inclusion detection was based on GFP intensity, ratio of inclusion mean GFP intensity/cell body mean GFP intensity, and ratio of inclusion area/cell body area. 2. Neuron death was detected as reduction in cell area or reduction in mean fluorescence intensity. The micrographs show the frame at which a GFP+ inclusion+ neuron was called dead by the algorithm (frame 19). 3. The output includes tracked neurons classified as inclusion+ or inclusion-, and the live/dead status at each captured frame, therefore providing the frame at which the neuron dies. Other output includes cumulative neurite length per well and soma inclusion size. (B) Timeline of neuron culturing and imaging for inclusion survival tracking in seeded inclusionopathy model. Induced neurons were transduced with CAG-RFP lentivirus on DIV10, seeded with PFFs on DIV11, and imaged in the Biostation CT for 10 days starting from DIV13. Inclusion+ neurons were tracked with GFP, and inclusion-neurons were tracked with RFP. (C) Survival tracking at the cellular level. pi-Ns of the indicated SNCA-sfGFP genotypes (A53T or ΔNAC), seeded or unseeded with PFFs, were longitudinally tracked by RFP fluorescence and revealed lower survival probability of neurons expressing αS-A53T compared to aggregation-dead α-syn-ΔNAC, and upon PFF exposure in both lines. This indicates both seeding-dependent and -independent toxicity of PFFs. Inclusion status was ignored in this analysis. Top: Example of RFP+ tracked pi-N^A53T-sfGFP-pB+PFF^ with white arrow indicating timeframe at which neuron dies. (D) Kaplan-Meier curve of inclusion+ (soma) and inclusion-seeded pi-N^A53T-sfGFP-pB+PFF^. Seeded A53T neurons with inclusions at the start of tracking had a lower probability of survival than those without inclusions throughout the tracking period. Right: Examples of 3 inclusion+ neurons that were tracked (number indicates cell tracked). n, number of neurons tracked. (E) Automated detection of GFP+ neuritic inclusions and cumulative length per well in seeded and unseeded pi-N of the indicated SNCA-sfGFP genotypes (A53T or ΔNAC). Data represent mean ± SD from 3 wells. (F) Inclusion survival tracking in spontaneous inclusionopathy model. Examples of pi-N^3K-sfGFP-pB^ and pi-N^sfGFP-pB^ control neurons detected in the Biostation. (G) Timeline of Biostation experiment with spontaneous inclusionopathy model. (H) Example of automated mask for soma-type inclusions. Neuron was identified as dead at the image captured at 90 h. (I) Kaplan-Meier curve comparing survival probabilities of pi-N^3K-sfGFP-pB^ with and without inclusions in the soma, and pi-N^sfGFP-pB^ control neurons. Log-rank test for panels C, D, I: * p<0.05, ** p<0.01, *** p<0.001, **** p<0.0001. Data in panels C-E and I are representative of two neuronal differentiations.

pi-N^A53T-sfGFP-pB^ neurons were transduced with CAG-RFP lentivirus at DIV10 for sparse labeling of neurons, exposed to recombinant PFFs at DIV11, and imaged every 6 h for 10 d (DIV13–DIV23) in the Biostation for longitudinal single-cell tracking (**Figure 4B**). First, we examined neuron survival irrespective of inclusion status, by detecting neurons with RFP signal, and found that pi-N^A53T-sfGFP-pB^ neurons had lower survival probability than pi-N^ΔNAC-sfGFP-pB^. Seeding with PFFs conferred a similar level of toxicity in both A53T and the aggregation-dead mutant ΔNAC, suggesting that PFFs result in aggregation-independent toxicity (**Figure 4C**). However, we cannot rule out that the inclusion-independent toxicity induced by PFFs was partially due to interactions with endogenous αS in the CORR iPSC line (**Figures 2F-2G**, **S2F**).

Next, we used the inclusion survival tracking algorithm, in which pi-N^A53T-sfGFP-pB+PFF^ neurons with GFP-positive inclusions in the soma are tracked with the GFP channel. Neurons without inclusions at the start of tracking were tracked via RFP signal because the GFP signal was too weak and diffuse. pi-N^A53T-sfGFP-pB+PFF^ neurons that harbored inclusions in the soma from the onset of tracking exhibited a higher risk of death than neurons that never developed inclusions (**Figure 4D**). The cumulative length of neuritic inclusions increased with time in PFF-seeded pi-N^A53T-sfGFP-pB^ neurons, but not in ΔNAC or unseeded neurons (**Figure 4E**). Notably, despite the pronounced presence of neuritic inclusions in seeded A53T neurons, the degree of toxicity that seeding conferred to A53T neurons was not detectably more than the non-specific toxicity it conferred to ΔNAC neurons (**Figures 4C** and **S5B**). These data suggest that in this model system neuritic inclusions may not be intrinsically toxic to neurons, but inclusions in the soma are toxic.

The 3K model (**Figures 3M-3N**) predominantly forms somatic inclusions. We conducted longitudinal single-cell tracking in this model to investigate the effect of inclusions on neuron survival (**Figure 4F-4H**). We identified and tracked inclusion(+) neurons through GFP, and inclusion(−) neurons through RFP, with optimized detection parameters (**Figure S4D**). Tracking of pi-N^3K-sfGFP-pB^ and control pi-N^sfGFP-pB^ neurons revealed that the former had a higher risk of death than the latter (**Figure 4I**). Among neurons expressing αS-3K-sfGFP, those with inclusions at the start of tracking had a higher risk of death than neurons that never developed inclusions (**Figure 4I**). Thus, αS-3K inclusions confer toxicity to neurons, similarly to the tracked inclusions within the soma of the seeded inclusionopathy model.

Thus, our algorithms can accurately track survival at single-cell and single-inclusion level, and pinpoint subsets of inclusions within the soma rather than neurites may be particularly toxic to human neurons within the timeframe examined. These data prompted us to further investigate the molecular subtype of these inclusions, each decorated by the standard neuropathologic marker pS129 but exhibiting distinct behaviors.

### Inclusion subtypes within pB-induced inclusionopathy models recapitulate those within postmortem synucleinopathy brain

Beyond the immunofluorescent labeling of morphologically and ultrastructurally heterogeneous intracytoplasmic inclusions in postmortem brain with pS129, immunohistochemical staining and proteomic analyses indicate that such inclusions comprise a vast number of other molecular components (Leverenz et al., 2007; Moors et al., 2021; Wakabayashi et al., 2013). These include the proteolytic degradation marker ubiquitin and the ubiquitin-binding p62 protein that transports its targets to autophagosomes(Kuusisto et al., 2001; Kuzuhara et al., 1988). Comparisons between ubiquitin and p62 in nigral inclusions revealed differences in immunoreactivity among morphologically heterogeneous inclusions(Kuusisto et al., 2003). o determine whether inclusions in our inclusionopathy models also stain for molecular markers typical of brain αS inclusions. Morphologically, most of the inclusions that form in the PFF-seeded tagged (pi-N^A53T-sfGFP-pB^) or untagged (pi-N^A53T-pB^) inclusion models appeared thread-like along neurites. Within somata, distinct subtypes of inclusions formed (**Figure 5A**), and were distinguishable by their resistance to detergents of differing strength during immunostaining (0.2% Triton X-100 or 0.1% saponin, data not shown). While all inclusions stained positive for pS129 (**Figures 5B** and **S6A-S6B**), they differed by subcellular localization (soma versus neurite) and staining with p62 or ubiquitin markers (**Figure 5C**, left, center; **S6A-S6B**). One week after seeding with PFFs, the majority (~80%) of inclusions in pi-N^A53T-sfGFP-pB^ soma were ubiquitin(+) (DIV18), whereas only a minority (~30%) were p62(+) (**Figure 5D**). However, three weeks after seeding (DIV32) the frequency of p62(+) inclusions increased to 80%. These data suggest that in our models ubiquitination precedes p62 labeling of seeded in inclusions. In contrast, the inclusions forming in the “spontaneous” inclusionopathy model (pi-N^3K-sfGFP-pB^ neurons) were all consistently negative for ubiquitin and p62, despite being pS129(+) (**Figure 5C**, right).

**Figure 5:**
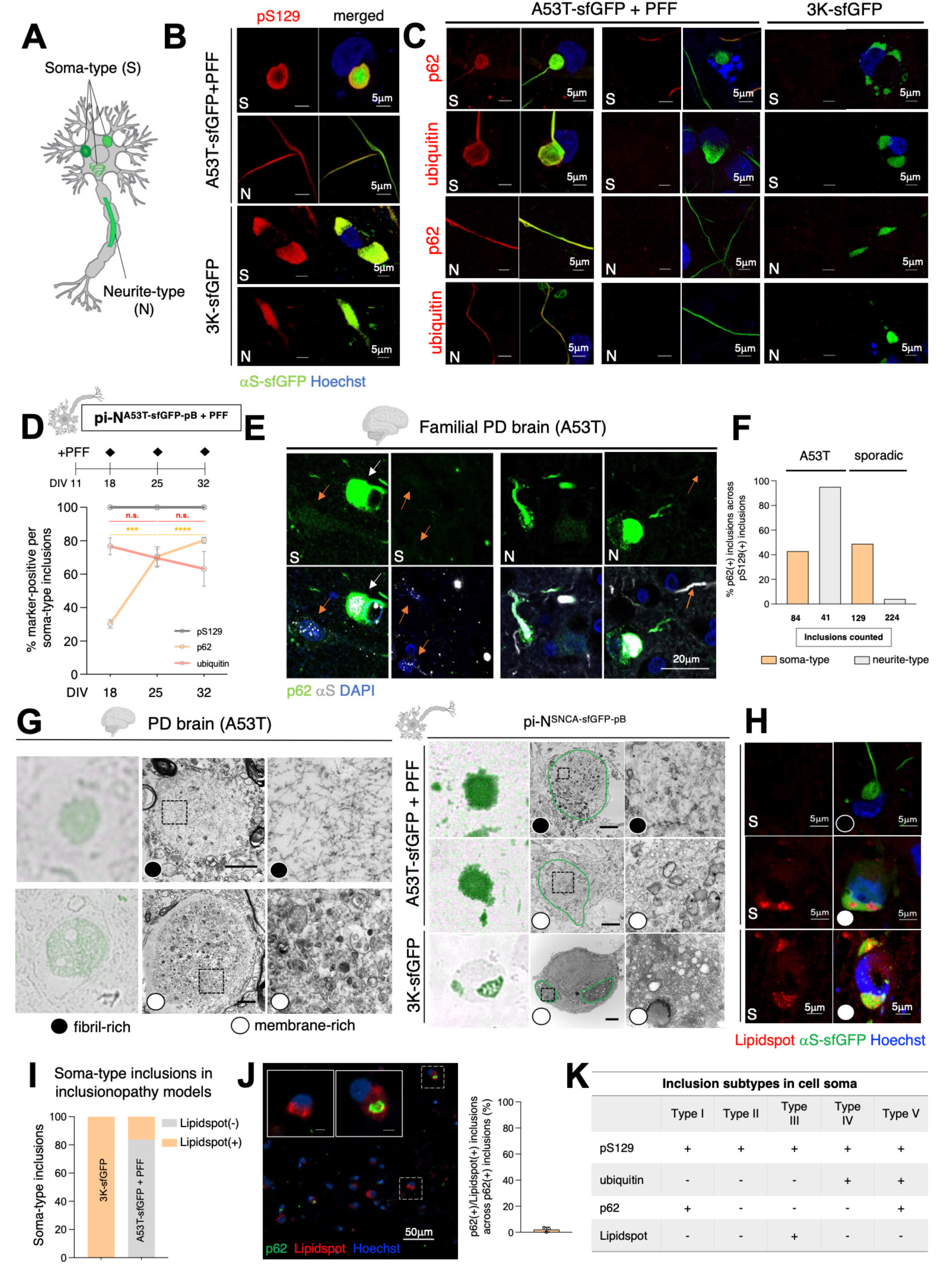
Molecular and subcellular classification of inclusions is conserved from pi-N model to postmortem brain. (A) Schematic diagram of inclusions classified by subcellular location. (B) Confocal microscopy examples of pS129+ soma-type (S) and neurite-type (N) inclusions in seeded and spontaneous inclusionopathy models. (C) IF of soma-type and neurite-type inclusions in the seeded inclusionopathy model showed that an inclusion subset stained positive for ubiquitin and/or p62. In the spontaneous inclusionopathy model, all inclusions stain negative for ubiquitin and p62. (D) Quantification of inclusions staining positive for pS129, p62 and ubiquitin at different DIV in seeded inclusionopathy pi-Ns showed that 100% of inclusions stained positive for pS129. Over 70% of inclusions stained positive for ubiquitin at 7 days post-seeding, whereas only ~30% of inclusions co-localize with p62. At DIV32, the fraction of p62(+) inclusions increases to ~80%. One-way ANOVA + Tukey’s multiple comparison test: *** p<0.001, **** p<0.0001. (E) IF for p62 and αS in cingulate cortex of familial PD (A53T) postmortem brain revealed subclasses of p62(+) and p62(−) soma-type or neuritic pS129(+) inclusions. (F) Quantification of frequency of p62(+) inclusions in soma and neurites in A53T familial and sporadic postmortem brain. 40% of soma-type inclusions in familial (A53T) and sporadic PD brain (n=1 each) were immunopositive for p62, and while >95% of neurite-type inclusions were p62+ in A53T brain, less than 5% of neurite-type inclusions in sporadic PD brain labeled for p62. (G) Correlative light and electron microscopy (CLEM) for αS in substantia nigra of familial PD (A53T) brain (left) and seeded versus spontaneous inclusionopathy models (right) uncovered ultrastructurally distinct inclusion subtypes rich in amyloid (left top; right top) or lipids (left bottom; right middle and bottom). (H) Neutral lipid dye Lipidspot labeled inclusions in the spontaneous inclusionopathy model as well as a subset of soma-type inclusions in the seeded inclusionopathy model. (I) Frequency of Lipidspot(+) and Lipidspot(−) soma-type inclusions in seeded and spontaneous inclusionopathy model. (ca. 5000 cells counted per condition) (J) IF for p62 and Lipidspot shows p62(+) inclusions were rarely Lipidspot(+). (K) Summary table of molecular markers in inclusion subtype classification.

Transgenic overexpression and tagging in our neuronal models provide shortcuts to a tractable model but may also trigger non-physiologic accumulation of αS. It was thus essential for us to confirm key findings in postmortem brain. We first asked whether inclusion heterogeneity noted in our models is also a feature in postmortem analysis. Ideally, since our model was established with the A53>T mutation, we would consider postmortem brain from carriers of that mutation also. Postmortem brains for αS mutation carriers are exceptionally rare and often suboptimally preserved (for example, delipidated and poorly amenable for EM). We thus cross-compared brains of two patients, one with A53T αS mutation and one with sporadic PD(Huang et al., 2011). We found subsets of p62(+) and p62(−) inclusions in the soma (perinuclear; **Figure 5E**, left) and neurites (**Figure 5E**, right) of the cingulate cortex, quantitated in **Figure 5F**. Thus, a key finding of inclusion heterogeneity identified in our culture models was borne out in postmortem brain.

We next turned to correlative light and electron microscopy (CLEM) to compare our A53T pi-N models with A53T postmortem brain. This technique enables ultrastructural features of αS-immunopositive inclusions to be revealed. Due to poor tissue preservation of the A53T brain, we were unable to assess the ultrastructural morphologies of inclusions in the cortex, but were able to do this in the substantia nigra. In the A53T brain, we identified αS(+) inclusions consisting of a dense fibril-rich core (**Figure 5G**, left top), and others rich in clustered vesicles and dysmorphic organelles (**Figure 5G**, left bottom), reminiscent of Lewy bodies and pale bodies, respectively, and similar to recent reports(Shahmoradian et al., 2019). An analogous study of the seeded pi-N A53T inclusion model, both sfGFP-tagged and untagged, revealed something very similar, namely a class of inclusions in the soma that was fibrillar (**Figure 5G**, right top) and another class that was composed of clustered vesicles, often interspersed with lipid droplets and containing dysmorphic mitochondria (**Figure 5G**, right middle, **S6D**). In contrast, inclusions detected by CLEM in pi-N^3K-sfGFP-pB^ were only of the vesicle- and lipid-rich class (**Figure 5G**, right bottom). These data suggested that, while the inclusions in the seeded pi-N A53T model were heterogeneous comprising fibril- and membrane-rich subtypes, the pi-N 3K model exhibited only the membrane-rich subtype that is also p62- and ubiquitin-negative.

The presence of lipid-rich inclusions was further corroborated by co-localization with the neutral lipid stain Lipidspot in both seeded pi-N A53T and spontaneous pi-N 3K inclusion models (**Figure 5H**). Close to 20% of soma-type inclusions in pi-N^A53T-sfGFP-pB^ were Lipidspot(+), whereas all inclusions in pi-N^3K-sfGFP-pB^ were Lipidspot(+) (**Figure 5I**). Few examples of inclusions that co-localized with both Lipidspot and p62 were detected in pi-N^A53T-sfGFP-pB^, reinforcing the idea that these two markers label two distinct classes of inclusions (**Figure 5J**). Thus, our pB inclusion models capture diverse inclusions—both fibril- and lipid-rich inclusion subtypes—that exist in PD patient postmortem brains. The molecular subclassification of pS129(+) inclusions (Types I-V based on markers) is depicted in **Figure 5K**.

Certain pharmacological modulators of αS cytotoxicity (trifluoperazine (TFP)(Höllerhage et al., 2014), nortriptyline (NOR) (Collier et al., 2017)) are known to rapidly clear cells of αS-3K inclusions (Imberdis et al., 2019). We confirmed this in our pi-N^3K-sfGFP-pB^ model (not shown) but also tested their effect on inclusions in the seeded inclusionopathy model. TFP and NOR selectively abrogated lipid-rich (Lipidspot(+)) cytoplasmic inclusions within minutes in a dose-dependent manner (**Figure S6E, Movies S1-S2**). However, within the same timeframes these tool compounds had no effects on fibrillar neuritic or ribbon cytoplasmic inclusions in our seeded inclusionopathy model.

### Intraneuronal fusion interaction events between inclusion subtypes alter neuronal survival

Membrane-rich and fibril-rich αS pathologies are not always distinct in postmortem brain. Fibril-rich Lewy bodies can appear at the periphery of membrane-rich pale bodies(Kuusisto et al., 2003). Indeed, classical Lewy bodies have membrane-rich components at their periphery. It is unclear whether pale bodies and Lewy bodies represent different stages of inclusion formation, whether pale bodies progress to compact Lewy bodies over time, or whether these are distinct entities that form in parallel(Dale et al., 1992). While postmortem human studies only offer a snapshot, prior work in rodent neurons suggested that maturing fibrillar inclusions seeded by PFFs ultimately incorporate lipids to form *bona fide* Lewy bodies over a period of 21 days(Mahul-Mellier et al., 2020). To better understand this in a human cellular model, we analyzed the pi-N^A53T-pB^ models in more detail because fibril- and lipid-rich inclusions coexist in these models (see **Figures 5G-5K**). Surprisingly, upon closer scrutiny, we noted that Lipidspot(+) (i.e. lipid-rich) inclusions occurred in this model prior to seeding with PFFs, regardless of tag. Thus, lipid-rich inclusions form spontaneously upon αS-A53T overexpression (**Figures 6A** and **S7A**). Even more unexpectedly, treatment with PFFs led to reduction in Lipidspot(+) soma-type inclusions despite increasing the overall frequency of pS129(+) inclusions (**Figure 6B** and **S7B**). These data suggested that in our model seeding with PFFs altered the balance between different inclusion subtypes.

**Figure 6:**
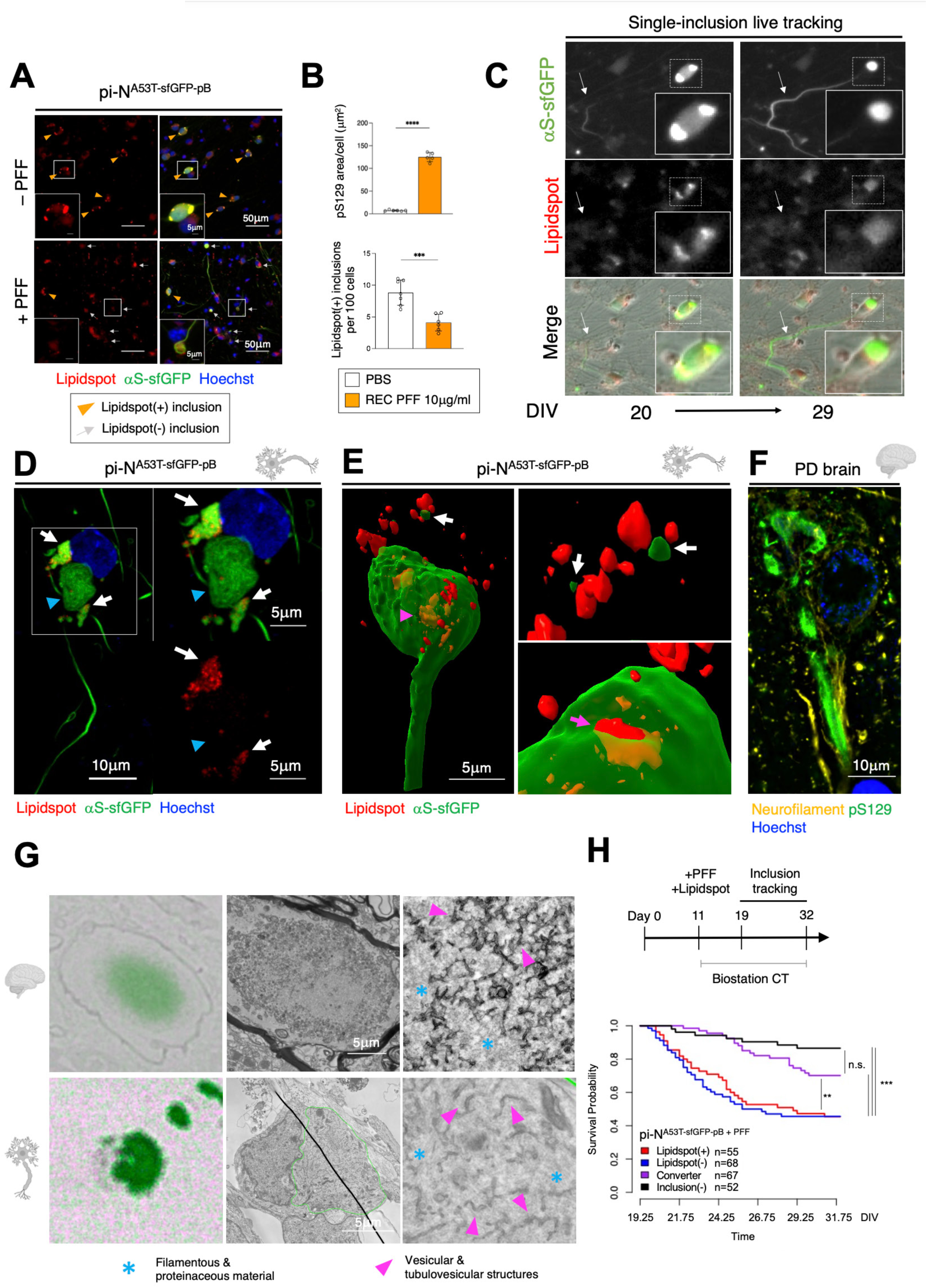
Fusion events between lipid-rich and presumed fibril-rich inclusions are detected. (A) IF for Lipidspot in pi-N^A53T-sfGFP-pB^ neurons showed lipid-rich Lipidspot(+) inclusions were present in the absence of PFFs (top) and decreased upon PFF seeding (bottom). Orange arrowheads, Lipidspot(+) inclusions; gray arrows, Lipidspot(−) inclusions. (B) Top: Quantification of pS129 area/cell in pi-N^A53T-sfGFP-pB^ +/− PFFs. Bottom: Concomitantly, the frequency of Lipidspot(+) inclusions decreases upon seeding with recombinant PFFs. One-way ANOVA + Tukey’s multiple comparison test: ***, p<0.001,**** p<0.0001. (C) Time-lapse-imaging in seeded PFF model showed interaction between Lipidspot(+) inclusions and elongating neuritic inclusions in the cell soma, possibly indicating a fusion event. White arrow indicates neuritic inclusion (GFP(+) Lipidspot(−)) that was absent in image from DIV20, but prominent in the image at DIV29. Inset shows cell soma with 2 GFP(+) Lipidspot(+) inclusions at DIV20, which becomes 1 GFP(+) Lipidspot(−) inclusion in the image at DIV29. (D) Confocal image example of adjacent lipid-rich Lipidspot(+) inclusion (top left GFP(+) inclusion with scattered red puncta), and a fibril-rich inclusion (bottom center GFP(+) inclusion) possibly representing a fusion event between the two inclusion subtypes. (E) Dynamic lattice light sheet microscopy (3D rendering) of a soma-type inclusion in the seeded inclusionopathy model live-stained with Lipidspot dye revealed the presence of accumulated lipids both within and surrounding the inclusion. White arrows indicate small αS-sfGFP positive aggregates; pink arrowheads point to sequestered lipid accumulations; pink arrow points to a lipid aggregate partially internalized into the inclusion. (F) IF for pS129 and neurofilament in frontal cortex of sporadic PD postmortem brain shows soma-type inclusion connected with neuritic inclusion and reminiscent of inclusion fusion examples in panels (D-E). (G) CLEM example of of αS(+) inclusions with mixed amyloid and lipid pathology, consisting of disorganized filaments, and vesicular and tubulovesicular structures in substantia nigra of sporadic PD postmortem brain (top) and seeded inclusionopathy model (bottom). (H) Manual longitudinal single-inclusion tracking of Lipidspot(+), Lipidspot(−), and Lipidspot(+) to (−) (“converter”) inclusions revealed that while Lipidspot(+) and Lipidspot(−) inclusions were toxic, conversion from Lipidspot(+) to Lipidspot(−) conferred a survival benefit. Log-rank test: ** p<0.01, *** p<0.001.

When we examined inclusion temporal dynamics by single-cell longitudinal tracking, we detected intraneuronal fusion interaction events between lipid-rich (Lipidspot(+)) inclusions and fibril-rich (Lipidspot(−)) inclusions. In examples where the cell soma contained Lipidspot(+) lipid-rich inclusions, movement of a Lipidspot(−) neuritic inclusion towards the cell soma resulted in dispersal of the Lipidspot(+) signal (**Figure 6C** and **Movie S3**). Consistent with a fusion process, neurons containing both Lipidspot(+) (white arrows) and Lipidspot(−) (blue arrowheads) inclusions within the same cell soma were also detected by confocal microscopy (**Figure 6D**). Dynamic lattice-sheet microscopy in 4D (x, y, z, time) enabled us to compellingly visualize areas in which solid neuritic inclusions appeared to protrude into lipid-rich structures in the cell. αS-sfGFP and Lipidspot(+) fragments directly opposed to each other in the vicinity of the apparent fusion event (**Figure 6E** and **Movie S4**). A similar observation of a pS129(+) neuritic inclusion seemingly continuous with a pS129(+) inclusion in the soma was made in PD brain (**Figure 6F**). We also identified inclusions by CLEM containing dense filamentous material immediately adjacent to vesicular structures in both postmortem brain and seeded inclusion model (**Figure 6G**).

To examine the potential biological effect of fusion events, we conducted manual single-inclusion survival tracking. We were keen to identify differences in viability among neurons in which fusion events occurred versus those in which they did not. pi-N^A53T-sfGFP-pB^ were seeded with PFFs followed by staining with Lipidspot on DIV11, and inclusions were tracked for 13 days (**Figure 6H**, top). Both classes of soma-type inclusions (“Lipidspot(+)”, “Lipidspot(−)”) conferred lower neuron survival probability compared to neurons that did not develop inclusions (“inclusion(−)”) (**Figure 6H**, bottom), confirming that distinct subtypes of soma-type inclusions are neurotoxic. Surprisingly, neurons with Lipidspot(+) inclusions at the start of tracking, which then converted to Lipidspot(−) over time (“converter”), our surrogate marker for a fusion event, showed improved survival probability. Altogether, data from our pi-N models suggested that intraneuronal fusion events between lipid-rich and fibril-rich inclusions occur in our iPSC models with analogous processes appearing to occur in brain. These events may be biologically meaningful, including through mitigating the toxicity of lipid-rich inclusions.

### Proximity labeling and membrane yeast two-hybrid (MYTH) analyses pinpoint proteins sequestered in membrane-rich inclusions

The distinct biological impact of different intraneuronal proteinaceous inclusions may relate to sequestration of different proteins and lipid species into these inclusions (**Figure 7A**)(Haenig et al., 2020). Because our data pinpointed lipid/membrane-rich αS inclusions (designated Type III in **Figure 5K**) as important drivers of cell autonomous toxicity in neurons (**Figures 4I** and **Figure 6H**), we decided to focus on which key proteins could be sequestered into this class of inclusions. To investigate this, we exploited the fact that the pi-N 3K model only exhibited lipid-rich inclusions. We took advantage of both existing and new protein interaction mapping datasets to narrow in on protein targets to test by immunostaining, and compare staining of different inclusion subtypes.

**Figure 7:**
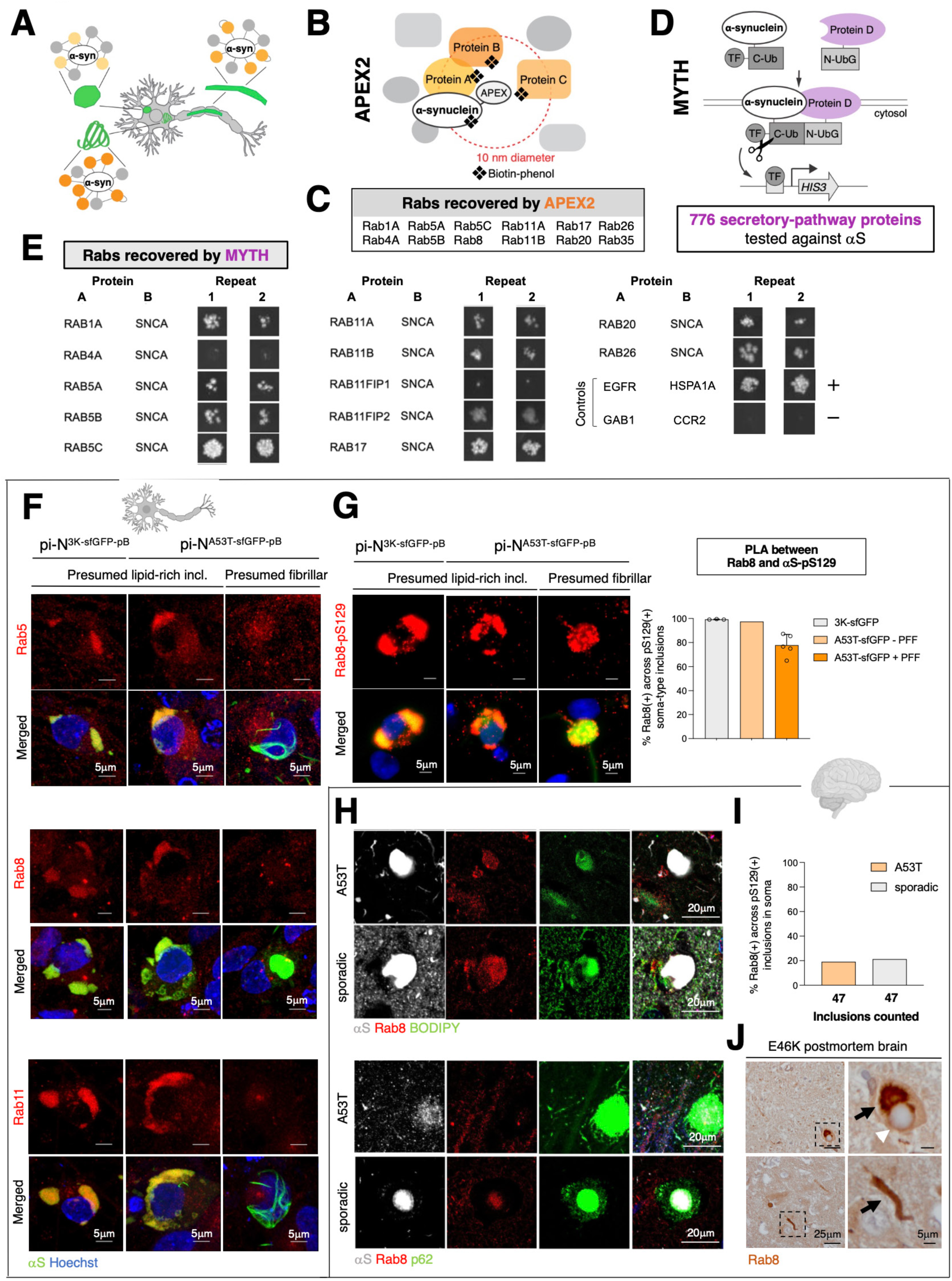
Proximity labeling and membrane two-hybrid as tools for identifying proteins sequestered in membrane-rich inclusions. (A) Schematic diagram of αS protein-protein interactions and how they might differ within different inclusion subtypes. (B) Schematic diagram of αS-APEX proximity labeling, in which a short pulse of peroxide activates APEX on the C-terminus of αS, creating a free radical from biotin tyramide and biotinylating proteins in the immediate vicinity (within 10 nm diameter) of αS. (C) Rab proteins found in the vicinity of αS by APEX (Chung et al., 2017). (D) Membrane yeast two-hybrid (MYTH) detected αS protein-protein interactions with human membrane-associated proteins expressed in *S. cerevisiae*. MYTH relies on a split ubiquitin system: a “bait” protein is fused to the N-terminal half of ubiquitin and directed to the ER membrane, while the “prey” protein (αS) is fused to the C-terminal half of ubiquitin. If the bait and prey proteins interact, the split ubiquitin is reconstituted, and cleavage by a deubiquitinase releases a transcription factor (TF) that controls expression of *HIS3*, thus allowing growth in media lacking histidine. (E) Rab proteins found to interact with αS by MYTH, as detected by growth in selective media. Repeat 1, 2 indicates replicate experiments. The EGFR-HSPA1A bait-prey pair was used as positive control, and GAB1-CCR2 pair as negative control in the assay. (F) IF in seeded (pi-N^A53T-sfGFP-pB+PFF^) and spontaneous (pi-N^3K-sfGFP-pB^) inclusionopathy models demonstrate Rab5, Rab8 and Rab11 co-localization with lipid-rich Type III inclusions, but less with fibrillar cytoplasmic inclusions. (G) *In situ* detection of interaction between Rab8 and pS129 by Proximity Ligation Assay (PLA). Rab8-pS129 interaction was detected more prominently in lipid-rich inclusions, but also detected in fibrillar inclusions. Bar graph shows quantification of pS129(+) soma-type inclusions that were also Rab8(+). Nearly all (>90%) pS129(+) cytoplasmic inclusions in pi-N^3K-sfGFP-pB^ and unseeded pi-N^A53T-sfGFP-pB^ were also Rab8(+), indicating that lipid-rich inclusions are Rab8(+). In contrast, fewer pS129(+) cytoplasmic inclusions in PFF-seeded pi-N^A53T-sfGFP-pB^ were Rab8(+) (~70%). Data points represent the average percentage of quantified inclusions per well across 42 image fields. (H) IF in anterior cingulate cortex of familial A53T PD and sporadic postmortem brain revealed Rab8 co-localization with Lipidspot(+)/αS(+) inclusions in cingulate cortex (top), but only in a subset of p62(+) inclusions (bottom). (I) Quantification of panel H (bottom). (J) Rab8(+) immunoreactivity (black arrows) in dystrophic neurites (bottom) and at the periphery of a Lewy body (top, white arrowhead) in postmortem brain (substantia nigra pars compacta) of patient harboring the SNCA-E46K mutation.

Previously, a network of 255 proteins in the immediate vicinity of αS in primary rat cortical neurons was defined by ascorbate peroxidase (APEX)-based labeling coupled with mass spectrometry (**Figure 7B**)(Chung et al., 2017). APEX2 hits included proteins related to endocytic vesicle trafficking, retromer complex, synaptic processes, phosphatases and mRNA binding proteins. Of the vesicle trafficking proteins, several members of the Rab family were prominently recovered (**Figure 7C**). To determine whether αS interacts (directly or indirectly) with those proteins, we utilized a binary interaction assay known as membrane yeast two-hybrid (MYTH) (**Figure 7D**) (Chung et al., 2017). Interactions of EGFR and ATP13A2 with 10 prey proteins each were included in the assay as positive controls, and 188 bait-prey pairs were included as random controls to evaluate background protein-protein interactions. In total, 776 proteins were tested for interactions with αS, with a strong focus on proteins in the secretory pathway. Of the 13 Rab proteins that interacted with αS in the MYTH assay, ten overlapped with APEX2 hits (Rab1A, Rab4A, Rab5A/B/C, Rab11A/B, Rab17, Rab20, Rab26) (**Figure 7E**). We prioritized a subset of these with good available antibodies for further analysis.

Certain Rab proteins co-localize with 3K inclusions in neuroblastoma models(Dettmer et al., 2017; Ericsson et al., 2021). We performed immunofluorescent labeling of Rab5, Rab8 and Rab11 in both our spontaneous (pi-N^3K-pB^) and seeded (pi-N^A53T-pB+PFF^) human neuronal models. In the seeded inclusion model, in keeping with their heterogeneous nature of lipid-rich versus fibril-rich inclusions (**Figures 5G-5K** and **6A**), Rab antibodies only decorated a subset of inclusions. Inclusions that were fibrillar based on ribbon-like morphology (“presumed fibrillar” in **Figure 7F**) were Rab immunonegative. Labeling was diffuse in control neurons without inclusions (**Figure S8A**). Thus, these Rab proteins (and potentially the vesicular compartments they are associated with) are enriched specifically within lipid-rich inclusions, consistent with our ultrastructural finding that these typically contain membrane-rich structures (**Figure 5G**).

The proximity ligation assay (PLA) utilizes oligonucleotide-hybridized antibodies to detect close protein-protein interactions *in situ*(Maio et al., 2016). It is more sensitive than immunofluorescence. We used it to specifically analyze close interactions between αS-pS129 and Rab8. PLA demonstrated αS-pS129-Rab8 interaction in pi-N^3K^ lipid-rich inclusions and presumed lipid-rich inclusions (by morphology) in seeded inclusion models (**Figure 7G**). Surprisingly, this sensitive assay could also detect Rab8 and αS-pS129 interaction, albeit less uniformly, in fibrillar inclusions. The PLA signal was specific to pS129 and Rab8, and abrogated in the absence of one or both primary antibodies (**Figure S8B**). Nearly all (>90%) pS129(+) soma-type inclusions were also positive for Rab8 signal in both pi-N^3K-sfGFP-pB^ and unseeded pi-N^A53T-sfGFP-pB^ neurons, in which inclusions are uniformly of the lipid-rich class (**Figure 7G**, right). In contrast, Rab8 and pS129 interaction was not detected in control neurons (**Figure S8C**).

Type I through V inclusion subtypes (**Figure 5K**) in our pi-N models correlate with inclusions identified in human postmortem brain (**Figure 5E**). Since lipid-rich Lipidspot(+) (Type III) and presumed fibrillary inclusions (Types I, IV, V) differentially co-labeled with Rab8 in our pi-N models (**Figures 7F-7G**), we asked if this held true in human brain also. While classic pale and Lewy bodies are identifiable with pS129 staining in the substantia nigra, they are not readily identifiable in the cortex. It is possible, however, that these dichotomous inclusion types, albeit far more subtle, are also present in the cortex. We hypothesized that αS interactors discovered in our inclusionopathy models may be useful in identifying them. Rab8 immunostaining did indeed co-localize with inclusions staining positive for the neutral lipid marker BODIPY in both A53T familial and sporadic PD postmortem brains (**Figure 7H**, top). As expected, Rab8 co-localization was not as prominent with p62+ inclusions (**Figure 7H**, bottom). The frequency of pS129(+) cytoplasmic inclusions that stained positive for Rab8 was lower (~20%) than in our neuronal models (**Figure 7I**), presumed in part due to the more sensitive PLA assay used in the pi-N models. Notably, Rab8-positive inclusions were also readily identifiable in an exceptionally rare postmortem brain analyzed from the αS-E46K kindred (**Figures 7J**, **S8D**).

Altogether, these data indicate that protein-interaction mapping can uncover markers that label specific inclusion subtypes in our pi-N inclusion models and human postmortem brain, providing more granularity than a generic marker of αS inclusions like pS129.

### Convergence of genetic and protein-protein interaction analyses enable identification of novel RhoA-positive inclusions in postmortem brain

Some proteins sequestered into inclusions might be causal to neurodegeneration, and others may just be bystanders. We asked whether our pB models could be used to identify the former category specifically for lipid-rich inclusions. One way to do this is to determine which proteins, when downregulated, lead to toxicity in the presence of 3K inclusions, but not with equivalent levels of WT αS overexpression. Those among these proteins that also colocalize with αS would be high-priority candidates for identifying in human brain because they might mark particularly toxic subsets of αS inclusions.

To identify at genome-scale genes that, when deleted, lead to toxicity only in the presence of 3K inclusions, we turned to CRISPR/Cas9, a system that enables genome-scale deletion screens in human cells(Naxerova et al., 2021; Shalem et al., 2013). Advantageously, the pB system can be adapted easily to cell lines as discovery model amenable to high-throughput genome-wide pooled genetic screening. A pB U2OS cell model expressing SNCA-3K-sfGFP along with equivalent levels of SNCA-WT-sfGFP and sfGFP in control lines, was generated (**Figures 8A** and **S9A**). The 3K model formed inclusions and showed cellular toxicity upon doxycycline induction, whereas few inclusions or toxicity were detected with SNCA-WT-sfGFP expression (**Figures 8A** and **S9A**). The 3K versus WT comparison allowed us to recover genetic modifiers in the context of mistrafficking compared to a model with equivalent levels of αS transgene overexpression but without inclusion formation. We also generated the analogous pB U2OS cell models expressing SNCA-A53T-sfGFP (A53T) and SNCA-A53T-DNAC-sfGFP (DNAC) (**Figure S9B**). As expected, inclusions were detected in A53T when seeded with recombinant PFFs, but not in DNAC U2OS cells (**Figure S9B**). Notably, unlike PFF-seeded inclusions in pB-A53T model (e.g. **Figures 3E, 4E**, **S9B**), 3K inclusions formed in a mostly NAC-independent manner, underscoring the biological distinctness of these inclusions (**Figure S9A**, right).

**Figure 8:**
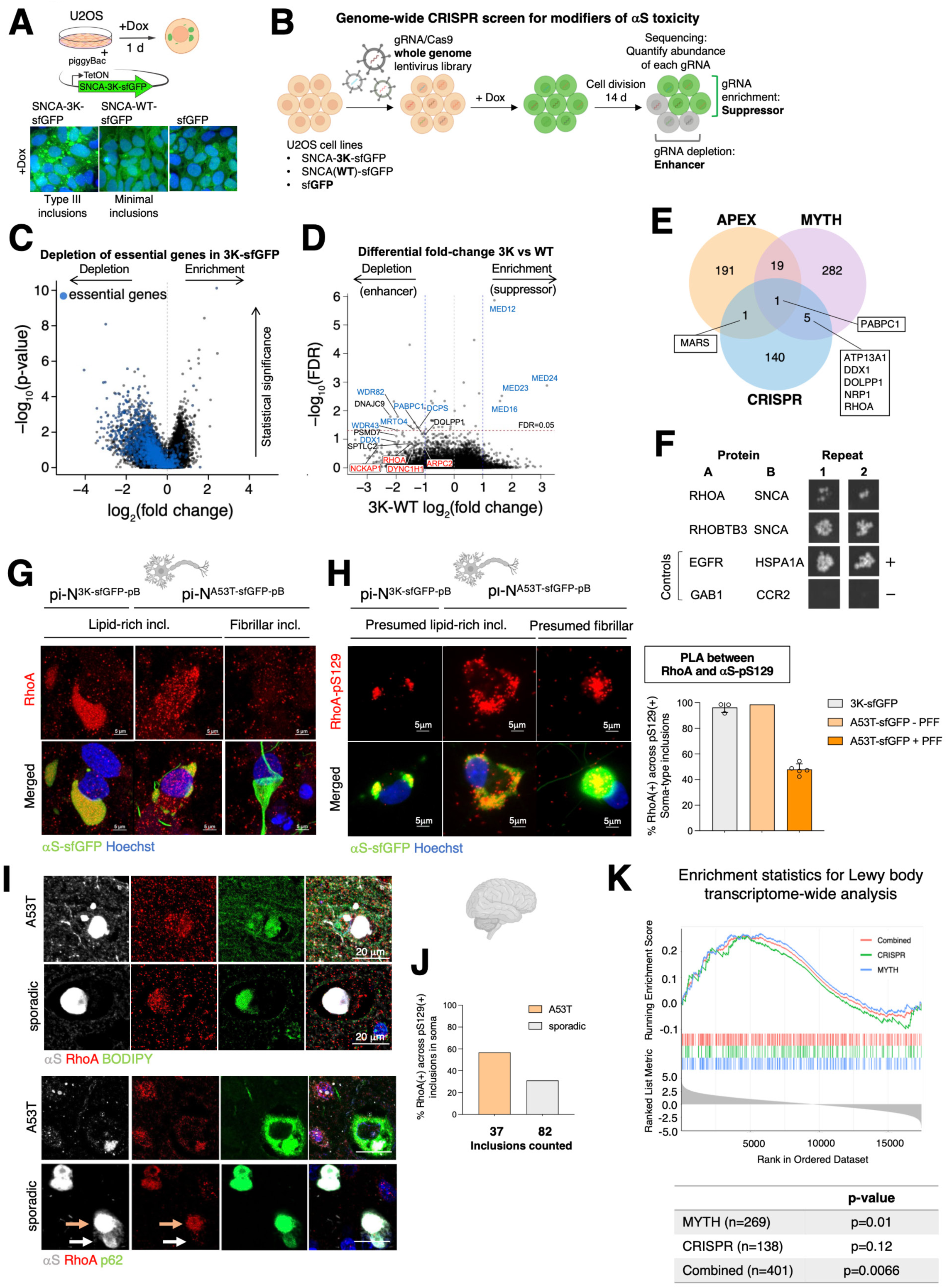
Convergence of CRISPR screen and MYTH on RhoA and other cytoskeleton regulators leads to identification of novel RhoA-positive inclusions in postmortem brain. (A) Cartoon of U2OS transgenic model harboring pB-SNCA-3K-sfGFP transgene. Micrographs show GFP signal from transgene in doxycycline-treated cells. Lipid-rich Type III inclusions were detected in cells expressing SNCA-3K-sfGFP, whereas minimal inclusions were detected in cells expressing SNCA-WT-sfGFP transgene. (B) Schematic of genome-wide CRISPR/Cas9 knock-out screen for modifiers of αS toxicity. (C) Volcano plot showing depletion of essential genes (blue) (Hart et al., 2015) in 3K-sfGFP screen. (D) Volcano plot comparing fold-change differential between SNCA-3K-sfGFP and SNCA-WT-sfGFP genotypes, and statistical significance for each target gene. (E) Venn diagram showing overlap between spatial (APEX, MYTH) and genetic (CRISPR) screen hits. (F) Actin cytoskeleton-related proteins RhoA and RhoBTB3 interacted with αS by MYTH. (G) IF for RhoA in seeded and spontaneous inclusionopathy models showed co-localization with lipid-rich Type III inclusions, but less with fibrillar inclusions. (H) RhoA-pS129 PLA showed *in situ* interaction within lipid-rich αS-GFP+ inclusions and the central core of a fibrillar inclusion. Right: Quantification of pS129(+) soma-type inclusions positive for RhoA in spontaneous and seeded inclusion models. (I) IF in anterior cingulate cortex of familial A53T PD and sporadic postmortem brain revealed RhoA(+)/ Lipidspot(+) inclusions (top) and a subset of p62(+)/RhoA(+) inclusions (bottom). Orange arrow indicates p62(+) inclusion with strong RhoA signal in sporadic PD postmortem brain; white arrow indicates region of inclusion that stains weakly for p62 and shows little/no RhoA signal. (J) Quantification of immunostainings in A53T and sporadic PD brain showing frequency of pS129(+) cytoplasmic inclusions that are also RhoA(+). (K) Gene set enrichment analysis of MYTH, CRISPR, or combined hits ranked according to association with Lewy body stage based on transcriptome-wide analysis.

U2OS cells harboring the pB doxycycline-inducible SNCA-sfGFP (WT or 3K) or sfGFP control transgenes were transduced with a ~90,000 sgRNA/Cas9 lentivirus library for genome-wide knock-out screening, followed by doxycycline treatment to induce transgene expression (**Figure 8B**). Cells were expanded and harvested at days 0, 7 and 14 post-induction. Genomic DNA was extracted and processed for next-generation sequencing to quantify the abundance of each gRNA. Depletion of gRNAs relative to t=0 is indicative of target genes that enhance toxicity when knocked out. Conversely, enrichment of gRNAs relative to t=0 indicate suppressors of toxicity when knocked out. Depletion of essential genes provided confirmation that the screening pipeline was effective (**Figure 8C**).

Genes selectively toxic to 3K versus WT cells when knocked out (**Figures 8D** and **S9C**) were enriched in GO Panther pathways such as positive regulation of cytoskeleton organization (FDR = 2.72×10^−2^), RNA metabolic process (FDR = 8.11×10^−8^), ribosome biogenesis (FDR = 1.34×10^−7^) and protein metabolic process (FDR = 7.81×10^−3^) (**Figures S9D-S9E, Tables S2-S3**). For example, genes encoding actin cytoskeleton regulators (including *ARPC2, RHOA*, *NCKAP1*, *DYNC1H1*) were recovered in the screen as enhancers of *SNCA* toxicity when deleted, as were RNA processing genes (*DCPS*, *WDR82*, *MRTO4*, *DDX1*, *DDX49,* etc), two classes of genes that have been previously tied to αS toxicity (Hallacli et al., 2022; Sarkar et al., 2021). Enhancers of3K toxicity also included genes relating to protein misfolding, aggregation, and lipid posttranslational modifications: heat shock protein family members (*DNAJC2*, *DNAJC9*), proteasome-related genes (*PSMD7*, *PSMG2*), prefoldin subunit (*PDRG1*)(Takano et al., 2014), and palmitoyltransferase (*SPTLC2*)(Ho et al., 2021). We next compared top hits in this CRISPR screen to top-hits in our MYTH αS-protein interaction assay. There were 6 overlapping proteins (**Figure 8E**). One, PABPC1, we have previously discovered as a genetic modifier of αS toxicity and to be translationally dysregulated in αS mutant neurons. It was also a top-hit in the APEX screen(Chung et al., 2017; Hallacli et al., 2022).

The Rho family members *RHOA* and *RHOBTB3* also emerged as protein interactors of αS in MYTH (**Figure 8F**). *RHOA* was a top CRISPR screen hit, suggesting that sequestration of RhoA and other cytoskeletal factors could be a key neurotoxic event associated with inclusion formation. We examined whether top hits from the CRISPR screen show altered subcellular localization in the presence of 3K inclusions. While ArpC2 exhibited a subtle change from punctate staining pattern in control neurons (pi-N^sfGFP-pB^), to a diffuse staining in the cytosol of neurons carrying lipid-rich inclusions (**Figure S10A**), it did not co-localize with inclusions. PABPC1 did not colocalize either (**Figure S10B**). In contrast, the Rho GTPase RhoA co-localized with lipid-rich (Type III) inclusions in spontaneous 3K inclusion model and presumed lipid-rich inclusions in seeded A53T model, but not with inclusions that were clearly fibril-rich based on morphology (**Figure 8G**). We confirmed the co-localization by PLA; RhoA-pS129 interaction *in situ* was detected in lipid-rich inclusions, and to a lesser extent in fibrillar inclusions (**Figure 8H** and **S10E**). In neurons that had solely lipid-rich inclusions (pi-N^3K-pB^, unseeded pi-N^A53T-pB^), nearly all pS129(+) cytoplasmic inclusions showed RhoA signal (**Figure 8H**, right). The frequency of pS129(+) inclusions co-localizing with RhoA signal decreased in PFF-seeded pi-N^A53T-pB^, presumably due to the increased diversity in inclusions. Few fibril-rich inclusions stained positive for RhoA, so their appearance upon PFF seeding shifted the relative frequency of RhoA(+) inclusions. Indeed, few (<5%) inclusions that were p62(+) (Type I/V in **Figure 5K**) exhibited RhoA signal (**Figure S10C-S10D**).

RhoA also marked lipid-rich inclusions in patient brain: immunostaining of familial A53T or sporadic PD patient brains showed BODIPY(+) inclusions staining positive for RhoA (**Figure 8I**, top; **Figure 8J**). In A53T brain, just as in our model, only occasional co-localization between RhoA and p62(+) inclusions was detected, but some inclusions were positive for both markers in sporadic PD brain (**Figure 8I**, bottom). Our pB transgenic models thus enabled the discovery of novel inclusion subtypes in the brain that are rich in RhoA, a protein that, when sequestered, is likely to be neurotoxic.

To establish whether our screens identified genes and proteins of relevance to synucleinopathy more broadly, we turned to the Religious Orders Study and Memory and Aging Project (ROS/MAP). ROS/MAP is a population-based study in which detailed measures of postmortem neuropathology can be directly related to a multitude of clinical and molecular phenotypes. We analyzed mRNA abundance in dorsolateral prefrontal cortex (DLPFC) of approximately 1000 brains(Felsky et al., 2022). DLPFC is matched to our iPSC cortical neuron model, but also a region with relatively early PD pathology, thus avoiding end-stage neuronal and glial responses. We specifically asked whether transcriptional changes in our top-hits were altered in response to accumulation of αS, as measured by Lewy body staging. We detected significant enrichment of MYTH (n=269 genes) and combined MYTH/CRISPR screen (n=401) gene sets with Lewy body stage (**Figure 8K**). The gene hits in MYTH and CRISPR/Cas9 showed enrichments in the positive direction with the Lewy body stage, indicating that they are increasingly dysregulated as Lewy body pathology advances.

## DISCUSSION

Proteinaceous aggregation in neurodegenerative diseases is often conceptualized as a single linear process – for example “monomer to oligomer to amyloid.” This may be an oversimplification that belies conformational, ultrastructural and spatial heterogeneity of proteinaceous inclusions that form in these diseases. Importantly, the heterogeneous nature of these inclusions is not readily distinguishable with common neuropathologic markers. Moreover, the heterogeneity may indicate different consequences for the cell and may help explain the mismatch that neuropathologists describe between extent of neurodegeneration and proteinaceous inclusion formation, and the many controversies that abound regarding whether inclusions are “protective” or “detrimental.” Neuronal models to date rely on either human stem cell-based models or rodent models that do not reliably recapitulate the advanced pathologies of CNS degenerative diseases in reasonable timeframes. Here, we exploited various technological advances – piggyBac technology, genome-editing, Gateway cloning, one-step iPSC transdifferentiation and single-cell longitudinal imaging. The result is a tractable, reproducible and readily transferable suite of human stem cell-based models for understanding the formation and consequences of diverse proteinaceous aggregates in human CNS cells.

To establish disease relevance, we required our models not only to recapitulate key findings in the brain, but also to discover new aspects of brain pathology. Proof-of-principle was established for synucleinopathies in which diverse cytoplasmic inclusions occur – lipid and membrane-rich pale bodies, amyloid fibril-rich Lewy bodies, and poorly understood hybrids of the two(Moors et al., 2021; Wakabayashi et al., 2013). Initially, we noted that seeding our models, whether pathologic overexpression of αS (triplication) or pB transgenic “pi-N” overexpression models, faithfully reproduced complex morphologies of brain inclusions (**Figure 2I**). Moreover, distinct biochemical properties could be induced in the system: exogenous seeding with PFFs induced pS129(+) inclusions with SDS-extractable aggregates (**Figures 3D** and **3G**). Aggregation of these inclusions was αS-NAC domain-dependent (**Figures 3D-3F**). In contrast, expression of αS membrane-avid E>K mutations(Dettmer, 2018), without any exogenous seeding, induced inclusions that were abundant and pS129(+) but fully extracted in Triton X-100 (**Figures 3N-3O**). Moreover, these inclusions formed in the absence of the αS-NAC domain. In both models, longitudinal single-cell and -inclusion resolution tracking pointed to somatic (rather than neuritic) inclusions being a primary site of cell-autonomous toxicity (**Figures 4D** and **4I**).

Immunofluorescence and ultrastructural studies defined somatic inclusions more clearly. Inclusions in the exogenously seeded PFF model were a mixture of fibril- and lipid-rich inclusions, analogous to Lewy and pale bodies in the brain, respectively. The ultrastructural similarities with EM and CLEM were clear (**Figure 5G**). Ubiquitinated nontoxic inclusions in neurites (**Figure 5C**), that ultrastructurally were also fibril-rich (**Figure S6C**) had counterparts in the soma that were initially ubiquitin(+) pS129(+) but evolved into ubiquitin(+) ps129(+) p62(+) inclusions over time (**Figure 5D**). An entirely separate second type of pS129(+) Lipidspot(+) ubiquitin(−) p62(−) lipid-rich inclusion formed spontaneously in the cell with αS overexpression and could be generated in high abundance and purity in the 3K pi-N model. Importantly, all subtypes of these somatic inclusions were identifiable in the human brain, leading us to generate a classification from Types I through V (**Figure 5K**) with more markers than phosphorylated S129 αS marker alone.

Lipid-rich inclusions were not only biochemically distinct from fibril-rich inclusions, but also exhibited a distinct dynamic behavior. First, these inclusions formed spontaneously with over-expression of αS. Second, longitudinal tracking revealed that these were rapidly “dissolved” by two pharmacological modulators of αS toxicity, TFP and NOR (**Figure S6E, Movies S1-S2**). In contrast, TFP and NOR had no effect on Lipidspot(−) inclusions. Third, we observed a physical interaction between Lipidspot(+) and Lipidspot(−) inclusions. Single-cell tracking revealed that neuritic Lipidspot(−) inclusions grow out contiguously from multiple seeded foci and never retract (**Movie S3)**. Strikingly, when they grew into the vicinity of Lipidspot(+) cytoplasmic inclusions, these lipid-rich inclusions were seemingly absorbed, a process captured after the fact by dynamic lattiche sheet microscopy (**Figures 6C-6E, Movie S4**). Surprisingly, such fusion events conferred a survival benefit to the neuron in which they occurred (**Figure 6H**). These findings may also shed light on mysterious hybrid inclusions that are noted in synucleinopathy. Classic Lewy bodies are often encircled with membrane-rich vesicles and organelles. We also present ultrastructural evidence of membrane and fibril-rich structures in close juxtaposition (**Figure 6G**). But in other cases, especially noted in the substantia nigra, fibril-rich Lewy bodies are juxtaposed against membrane-rich pale bodies (Wakabayashi et al., 2013). The assumption has been that one inclusion is “evolving” into the other, but our dynamic imaging suggests that these mixed inclusions may result from fusion events and impact cell survival. The sequestration of soluble forms of αS into stable proteinaceous inclusions may also compromise the endogenous function of αS.

Rather than simply label an aggregating protein within an inclusion (for example, with αS pS129) a more biologically meaningful way to subtype may be to co-label proteins sequestered into the inclusion. Our models provide a discovery tool through which to define such components. We focused on lipid/membrane-rich αS somatic neuronal inclusions because these were clearly toxic. Moreover, in the cortex, where morphologic pale bodies and Lewy bodies are hard to distinguish, we hypothesized that certain sequestered proteins would be informative in delineating inclusion subtypes more meaningfully than pS129. Previously, we utilized proximity labeling with APEX2 (**Figures 7B-7C**) to identify numerous secretory pathway proteins in the immediate vicinity of αS(Chung et al., 2017). To identify those secretory pathway proteins that form multiprotein complexes with αS, we tested ~800 secretory proteins with the MYTH method(Chung et al., 2017; Stagljar et al., 1998) (**Figures 7D-7E**). These datasets converged on numerous Rab proteins that colocalized with lipid/membrane-rich-inclusions (**Figure 7F**) and we confirmed that Rab8 was an effective marker for identifying this subset of inclusions in postmortem brain from A53T and sporadic PD patients (**Figures 7H-7I**).

Finally, we developed an approach to distinguish between proteins that are merely sequestered into inclusions as bystanders versus sequestered proteins that actually render cells vulnerable. We focused specifically on factors that render cells vulnerable in the inclusion-positive state. pB constructs can be just as easily introduced into cell lines as iPSC cells to create analogous inclusion models (**Figure 8A** and **S9A**). In a genome-wide CRISPR/Cas9 screen, we identified genes that, when knocked out, led to lethality in cells that developed αS-3K inclusions but not in cells expressing equivalent levels of non-inclusion bearing cells that express wild-type αS (**Figures S8D**, **S9C**). RNA processing and actin cytoskeleton modulators emerged as major classes of proteins that, when knocked out, led to specific dropout of 3K-expressing cells (**Figure S9D**). These pathways have both been heavily implicated recently in synucleinopathy(Hallacli et al., 2022; Ordonez et al., 2018; Sarkar et al., 2021). Among these, 7 proteins overlapped with our spatial αS mapping screens (**Figure 8E**). One of these, the cytoskeletal regulator RhoA, labels a subset of inclusions in sporadic and familial synucleinopathy brain (**Figure 8I-8J**) and we suggest it may be a marker of particularly toxic αS intraneuronal inclusions. In future investigations, it will be important to analyze our models for non-proteinaceous components of the cell that are also now known to be sequestered into fibrils(Yang et al., 2022b).

We have previously mapped systematic genetic and physical interactors of αS in non-human cellular models. These maps have connected αS to numerous genetic risk factors for PD(Jarosz and Khurana, 2017; Khurana et al., 2017; Lam et al., 2020). We extend that work in the current paper with genetic mapping in U2OS pB models and spatial mapping with MYTH. We can now envisage systematic primary mapping of αS interactors in distinct cell types of the CNS in isolation and combination across different αS mutations or conformers (for example DLB versus PD versus MSA “strains”). Direct mapping strategies in human CNS cell types will avoid the drawbacks of simpler systems, for example the polyploidy that may have limited our U2OS screen. Our pB system can also easily be introduced into standardized isogenic iPSC lines, an unprecedented resource now being generated by the stem-cell community for disease modeling(Pantazis et al., 2022). Moreover, the pi-N system, especially with Gateway cloning compatibility built in, can readily be extended beyond synucleinopathy to other proteinopathies.

### Limitations of the study

There are numerous technical workarounds we introduced into our models for tractability. Each of these has potential drawbacks. For example, our models use doxycycline for induction of the transcription factor (NGN2, NFIB, etc) in iPSC/hESC transdifferentiation protocols, and in some αS overexpression models (*AAVS1*, pB). Doxycycline is known to be biologically active in neurons, especially implicated in altering mitochondrial function(Moullan et al., 2014), and more recently in attenuating αS aggregation(Dominguez-Meijide et al., 2021). This may limit the use of downstream pathways, especially when doxycycline is chronically needed for transgene expression. It is possible that readouts related to mitochondrial and lysosomal assays in our pB neurons (**Figures S3A-S3E**) were confounded by the presence of doxycycline. We are actively developing and testing doxycycline-independent alternatives. We also present a safe-harbor alternative (*STMN2*) in which neuronal overexpression occurs in the absence of doxycycline. Notably, the safe-harbor alternatives we present (*AAVS1* safe-harbor locus knock-in, *STMN2* lineage-specific knock-in) offer the advantage of single-site genome integration rather than the random integration of pB. Random integration could introduce unpredictable genetic effects and lead to genome instability. In our group we have used the splinkerette assay to assess integration sites(Uren et al., 2009) but more typically we create polyclonal pB lines to avoid oversized effects of single-site mutations. We then expand, karyotype and use these polyclonal lines at very early passage.

Aside from the *SNCA* triplication lines, the cell models we present here also all utilize transgenic overexpression to trigger robust inclusion formation in a reasonable timeframe. That being said, we have found and confirmed that endogenous expression of αS in iPSC-derived neurons is far lower than in postmortem brain, especially in neurons. Indeed, our pB models quite faithfully recapitulate the levels of αS in brain (**Figure 2D**). Beyond this, as noted above, we extensively cross-compared our models to postmortem brain to demonstrate disease-relevance.

In future studies, efforts could be made to accelerate both maturation (Hergenreder et al., 2022) and aging (Miller et al., 2013; Vera et al., 2016) “in the dish” and to combine neurons and glia in co-culture (we purposely showed proof-of-principle that our tractable pB system could be used for glia just as easily; **Figures 1D-1E; S1A-S1B**). An increasing number of CNS cell types are being generated through transdifferentiation (Chen et al., 2021; Dräger et al., 2022; Holzer et al., 2022; Limone et al., 2022; Sonn et al., 2022). Recent studies(Guttikonda et al., 2021) show that the combination of glia in the right proportions are important for triggering appropriate neuroinflammatory responses. Such non-cell-autonomous effects will be critical to examine in our models. For example, the toxicity of PFFs may be deeply related to neuroinflammation(Garcia et al., 2022; Iba et al., 2022; Stoll and Sortwell, 2022). The relative lack of toxicity we see with PFFs in our neurons departs from what is seen in vivo in mouse models(Luk et al., 2012; Mao et al., 2016; Park et al., 2021) and may relate to a lack of inflammatory responses. Beyond 2D cultures, certain non-cell-autonomous effects may only be recapitulated in 3D sphere and organoid systems. We envisage our models will also be very helpful in creating more reproducible 3D co-cultures. Beyond culture models, iPSC-derived models can be engrafted to create mouse/human chimeras. Such models would allow modeling of inclusion formation and dynamics in more disease-relevant timescales, including with multi-photon imaging (Osterberg et al., 2015).

## Conclusion

Ultimately, the inclusionopathy models we describe balance tractability with disease-relevance to take up several key challenges in neurodegeneration. First, they bring a tractable suite of tools to tackle the daunting heterogeneity of inclusions in CNS proteinopathies, to better molecularly characterize inclusion subtypes. We can envisage in the future that each major subclass will be definable and assigned to a distinct biological consequence, for example on cell survival. These insights may lead to a more thorough classification and understanding of inclusion subtypes in the CNS. Second, these models will enable systematic cross-proteinopathy analysis in different CNS cell types. And, finally, these models offer a simple, scalable path to creating a patient-specific model that incorporates both “host” cells and proteinaceous “strains.” In conjunction with technologies that now enable us to amplify aggregating proteins from patient tissue and body fluids(Kuzkina et al., 2021; Russo et al., 2021), our models may offer a tractable path to patient-specific models in which diagnostics like radiotracers, or therapeutics like antibodies, can be readily tested and ultimately matched to individual patients.

## ACKNOWLEDGEMENTS

We thank all the members of the ARCND community for their critical input on the paper. We thank Lai Ding at BWH NeuroTechnology studio for Fiji macros; Jian Peng for assistance with R scripts for survival plots; Maria Ericsson and Margaret (Peg) Coughlin at the Harvard Medical School Electron Microscopy Facility for help with processing samples. We thank Tim Moors for helpful discussion. Select schematics within figure panels (Graphical abstract, Figures 1A-1D; 2H-2I, 3B, 8A-8B) were generated with illustrations from BioRender.com. I.L. was supported by American Parkinson Disease Association (APDA), PhRMA Foundation, and American Academy of Neurology (AAN). A.N. was supported by a Max Kade Fellowship. N. Morshed is supported by a T32 training grant (5 T32 AG 222-30). B. Stevens stem cell lab is supported by the Stanley Center for Psychiatric Research and Howard Hughes Medical Institute. N. Sahni was supported by the NIH grant R35GM137836. S. Yi was supported by NIH grant R35GM133658. Theresa Bartels was supported by a scholarship of the Felgenhauer Foundation for the Support of Young Neuroscientists and a DAAD PROMOS scholarship from the German Academic Exchange Service. V.K. is a New York Stem Cell Foundation Robertson Investigator (NYSCF-R-I49) and a George C. Cotzias Fellow of the APDA. V.K. is supported by NIH R01NS109209. Additional grants to V.K. that supported this work include Aligning Science Across Parkinson’s Initiative ASAP-000472, Brigham Research Institute Director’s Transformative Award, Michael J. Fox Foundation 18768 (Ken Griffin Alpha-Synuclein Imaging Competition). The project was begun through seed funding from the Multiple System Atrophy Coalition.

## AUTHOR CONTRIBUTIONS

I.L., A.N., and V.K. designed the project. I.L., A.N., and V.K. wrote the manuscript with input from co-authors. I.L. and A.N. conducted and analyzed all experiments with the exception of those specified below. They received assistance from A. Verma, C.O.-S., M.S., X.Y., K.H. and C.P. A.J.L. performed CLEM on postmortem brain and iPSC-derived neurons with input from H.S. Y.F. and G.T.S performed quantitative immunofluorescence on postmortem brain samples with experimental input and supervision from G.M.H. L.Z. conducted SAA, ThT and Proteinase K digestion assays with supervision from Tim Bartels. J.S. constructed the pB-NGN2 plasmid and STMN2 targeting. A. Vahdatshoar cloned derivatives of pB-NGN2 plasmid and generated pB cell lines. R.L.S. and N.R. established and characterized pB U2OS models. R.L.S., T.D.M., S.J.E., V.K. conceptualized U2OS CRISPR/Cas9 screen. I.L. and A. Verma conducted this CRISPR/Cas9 screen with supervision from T.D.M; T.D.M. analyzed data from this screen. N.M. contributed pB-NFIB plasmid and provided experimental input on astrocyte differentiation, supervised by B.S. T.I. and Y.K. developed Biostation inclusion survival algorithms. A.T. and U.D. contributed to experiments related to pharmacological modulation of inclusions. T.W.T. and H.W. performed gene targeting for the AAVS1 model with input and supervision by R.J. Theresa Bartels analyzed these AAVS1 lines with supervision from V.K., C.Y.C., T.W.T and S.L. E.H. performed STMN2 qPCR experiment. C.T., Z.W. and H.H. performed dynamic lattice-sheet microscopy with supervision from J.S. N.S. and G.W. provided supervision and intellectual input to A.N. K.C.L. provided A53T PFFs. I.F. conducted immunohistochemistry on E46K postmortem brain provided by R.S.P. and J.C.G.-E. D.F. conducted the ROSMAP analysis. N.S. and S.S.Y. performed and analyzed MYTH experiments. V.K., C.Y.C., and X.J. oversaw iPSC reprogramming and genetic corrections.

## METHODS LIST

- Molecular cloning
- Generation of targeted inducible transgene at *AAVS1* locus in hESC via TALENs
- Generation of targeted transgene at *STMN2* locus in hESC via CRISPR/Cas9
- Stable integration of piggyBac plasmids into iPSCs
- Stable integration of piggyBac plasmids into U2OS cells
- U2OS cell culture
- iPSC generation and lines
- Induced neuron differentiation
- Induced astrocyte differentiation
- Conventional differentiation
- Recombinant aS expression and purification
- Generation of preformed aS fibrils
- PFF seeding in iPSC-derived cortical neurons
- PFF seeding in U2OS cells
- Electron microscopy
- Whole cell protein extraction
- Western blotting
- Immunofluorescence and microscopy
- Lattice light-sheet microscopy and 3D rendering
- Sequential extraction of insoluble fraction from human brain
- Real time-quaking induced conversion (RT-QuiC) (Seeded amplification assay)
- ELISA
- Proteinase K digest
- αS Triton X-100/SDS sequential extraction
- Seahorse XF Cell Mito Stress Test
- Autophagic flux assay
- Automated longitudinal single-cell inclusion survival tracking
- Lipidspot live-cell staining and manual quantification
- Live-cell compound treatments and imaging
- Immunohistochemistry in post-mortem brain
- Correlative light- and electron microscopy (CLEM) in iPSC-derived neurons and postmortem brain
- Membrane Yeast Two-Hybrid
- Proximity ligation assay (PLA)
- Genome-wide CRISPR/Cas9 screen in U2OS cells
- ROS/MAP analysis

## METHODS

### Key Resources Table

#### Antibodies for immunofluorescence and western blotting

| Reagent or Resource | Source | Identifier |
| --- | --- | --- |
| Anti-Alpha-Synuclein | BD Biosciences | Cat. # 610786 |
| Anti-Alpha-Synuclein | R&D | Cat. # AF1338 |
| Anti-Alpha-Synuclein (4B12) | Thermo Fisher | Cat. # MA1-90346 |
| Anti-Alpha-Synuclein (Syn1) | BD Transduction Laboratories | Cat. # 610787 |
| Anti-AQP4 | Sigma | Cat. # CL0178 |
| Anti-ArpC2 | Abcam | Cat. # 133315 |
| Anti-Beta-Tubulin III | Stem Cell Technologies | Cat. # 60052 |
| Anti-Brn2 | Cell Signaling | Cat. # D2C1L |
| Anti-Ctip2 | Abcam | Cat. # ab18465 |
| Anti-Cux1 | Abcam | Cat. # ab54583 |
| Anti-GAPDH | EMD Millipore | Cat. # MAB374 |
| Anti-GAPDH | Thermo Fisher | Cat. # MA5-15738 |
| Anti-GFAP | Dako, Agilent Technologies | Cat. # Z0334 |
| Anti-GFP | Abcam | Cat. # 6556 |
| Anti-GFP | Roche Diagnostics | Cat. # 11814460001 |
| Anti-GFP | Rockland | Cat. # 600-101-215 |
| Anti-LC3A/B | Cell Signaling | Cat. # 4108 |
| Anti-p62 | EMD Millipore | Cat. # MABN130 |
| Anti-Phospho-Ser-129 | Abcam | Cat. # ab51253 |
| Anti-Rab5 | Cell Signaling | Cat. # C8B1 |
| Anti-Rab8 | BD Biosciences | Cat. # 610844 |
| Anti-Rab11 | Cell Signaling | Cat. # D4F5 |
| Anti-Rab35 | ProteinTech | Cat. # 11329-2-AP |
| Anti-RhoA | Cell Signaling | Cat. # 2117 |
| Anti-RhoA | Santa Cruz | Cat. # sc-166399 |
| Anti-S100b | Sigma | Cat. # S2532 |
| Anti-Tbr1 | Abcam | Cat. # ab18303 |
| Anti-TUJ1 | BioLegend | Cat. # 801201 |
| Anti-Ubiquitin (FK2) | EMD Millipore | Cat. # ST1200 |
| Anti-VGLUT1 | Synaptic Systems | Cat. # 135303 |
| Anti-Vimentin | EMD Millipore | Cat. # CBL202 |
| BODIPY 493/503 | Thermo Fisher | Cat. # D3922 |
| LipidSpot 610 | Biotium | Cat. # 70069 |
| Total OXPHOS Human WB Antibody Cocktail | Abcam | Cat. # ab110411 |

#### Resource availability

##### Lead Contact

Further information and requests for resources and reagents should be directed to and will be fulfilled by the lead contact, Dr. Vikram Khurana

##### Material availability

All unique/stable reagents generated in this study are available from the lead contact with a completed materials transfer agreement.

## METHOD DETAILS

### Molecular cloning

For Gateway cloning, gene blocks (double stranded DNA fragments) and primers were purchased from IDT (Integrated DNA Technologies). LR and BP clonase mix were purchased from Invitrogen and used per recommended protocol from supplier for Gateway cloning. Donor or destination plasmids containing ccdB sequence were propagated in ccdB-resistant *E. coli* strain One Shot ccdB Survival 2 T1R Competent Cells (Life Technologies). Expression clones were transformed into 10-beta competent *E. coli* (NEB).

### Generation of targeted inducible transgene at *AAVS1* locus in hESC via TALENs

To establish Tet-On system transgene at the *AAVS1* locus within the *PPP1R12C* gene, two rounds of TALEN-mediated gene editing were conducted in hESC lines (male WIBR-1, clone 22, or female WIBR-3, clone 38). First, one construct containing the M2rtTA reverse tetracycline transactivator under the control of the constitutive CAGGS promoter (P_CAGGS_-M2rtTA) was targeted to one *AAVS1* allele. The second *AAVS1* allele was subsequently targeted with a construct containing the transgene of interest driven by the M2rtTA-responsive TRE-Tight promoter (e.g., P_TRE-Tight_-SNCA-mK2). Both constructs have flanking 5’ *AAVS1* and 3’ *AAVS1* homology arms.

#### Southern blotting

Correct integration of the Tet-On constructs at the AAVS1 locus within the *PPP1R12C* gene was confirmed by Southern blot analysis. An AAVS1 internal 5’-probe, corresponding to the 5’ homology arm of the AAVS1 donor targeting vector, was used to detect extra integration sites beyond the AAVS1 locus. An AAVS1 external 3’-probe, which hybridizes with a sequence downstream of exon 3 of the *PPP1R12C* gene, was used to confirm integrity of the AAVS1 locus.

Genomic DNA was extracted according to the manufacturer’s instructions (DNeasy Blood and Tissue Kit, Qiagen) from hESCs harvested from a well of a 12-well plate, at 70-90% confluency. Genomic DNA was digested with EcoRV-HF restriction enzyme according to the manufacturer’s instructions (New England Biolabs). DNA restriction fragments were size-fractionated by electrophoresis in a 0.8% agarose gel (SeaKem GTG agarose, Lonza) in Tris-acetate-EDTA (TAE) electrophoresis buffer containing 0.5 μg/mL ethidium bromide (Thermo Fisher Scientific). The gel was washed for 15 min in 0.25 M HCl solution (nicking buffer) at 80 rpm, followed by 15 min at 80 rpm in 0.4 M NaOH solution (denaturing and transfer buffer), and assembled in a transfer stack for alkaline Southern transfer of the single-stranded DNA fragments onto a nylon membrane (Amersham Hybond-XL, GE Healthcare). Southern transfer was conducted overnight via upwards capillary action mediated by the transfer buffer. The next day, the transfer membrane was rinsed in 0.2 M Tris-Cl, pH 7.0 and 2X saline-sodium citrate (SSC; 0.3 M NaCl with 7.5 mM trisodium citrate), for 2 min each at 80 rpm. The transfer membrane was dried for 15 min in a 55°C oven, followed by a pre-hybridization (blocking) step with hybridization buffer (1% [w/v] bovine serum albumin/BSA, 1 mM ethylenediaminetetraacetic acid/EDTA, 0.5 M NaPO_4_, 7% [w/v] sodium dodecyl sulfate/SDS in deionized water; all Sigma-Aldrich) for 1 h in a 60°C hybridization oven with rotation.

In preparation for radioactive labeling of the AAVS1 internal 5’-probe, a restriction fragment within the 5’ homology arm was derived by restriction endonuclease digestion of the AAVS1 donor targeting vector with SacI and EcoRI according to the manufacturer’s instructions (NEB). DNA restriction fragments were size-fractionated by electrophoresis in a 1% agarose gel as described above and the 643 bp restriction fragment was recovered after gel excision using silica membrane spin columns according to the manufacturer’s instructions (MinElute gel extraction kit, Qiagen). DNA concentration was determined with a NanoDrop ND-1000

Spectrophotometer. Radiolabeling of the 5’-probe was carried out by random-sequence oligonucleotide-primed DNA synthesis. A 28.5 µL reaction volume containing 100 ng of the 5’ homology arm fragment and 5 µL of 50 µM random nonamers (Sigma-Aldrich) in Ambion nuclease-free water (Thermo Fisher Scientific) were incubated for 5 min at 100°C for denaturation into single-stranded DNA. After 5 min on ice, 5 μL 10X NEBuffer 2 (NEB), 5 μL 100 mM 3dNTPs (minus dCTP; Thermo Fisher Scientific), 5 μL of the radioactively labeled nucleotide [α-^32^P]dCTP (PerkinElmer; 10 μCi/μL) and 1.5 μL Klenow fragment of the *E. coli* DNA Polymerase I (NEB) were added for a final volume of 50 µL, and incubated for 30 min at 37°C. The reaction was stopped with 50 µL of buffer TE (Qiagen), and the radiolabeled probe DNA was separated from unincorporated dNTPs by gel filtration chromatography using pre-equilibrated CHROMA SPIN columns (Clontech) with centrifugation at 3,500 rpm for 5 min. The double-stranded probe DNA was denatured for 5 min at 100°C.

The transfer membrane was hybridized with the single-stranded 5’-probe DNA, diluted in fresh hybridization buffer, overnight in the 60°C hybridization oven with rotation. After the hybridization step, the DNA blot was washed at low-stringency in 2X SSC with 0.2% (w/v) SDS for 30 min in a gently shaking 60°C water bath. Any remaining nonspecifically bound probe DNA was washed off during a high-stringency wash with 0.2X SSC (0.03 M NaCl with 0.75 mM trisodium citrate) with 0.2% (w/v) SDS for a minimum of 20 min in a 60°C water bath with gentle shaking. The membrane was sealed in Saran wrap, placed between an autoradiography film (Carestream Kodak BioMax MS film, Eastman Kodak) and an intensifying screen (Eastman Kodak), exposed for 24-72 h at −80°C, brought to room temperature, and developed using the Kodak X-OMAT 1000A film processor.

To re-hybridize the DNA blot with an AAVS1 external 3’-probe, the transfer membrane was rinsed in 0.08 M NaOH solution (stripping buffer) for a minimum of 15 min at room temperature with gentle shaking. The transfer membrane was washed three times for 5 min with 2X SSC. If any radioactive signal was still detectable, the nylon membrane was stripped in 0.4 M NaOH for 30 min at room temperature, with gentle shaking. The transfer membrane was dried in a 55°C oven before the pre-hybridization, hybridization and autoradiography steps were repeated for the external 3’-probe (a gift from the Rudolf Jaenisch laboratory, Whitehead Institute for Biomedical Research) as described above.

The size of the DNA restriction fragments as detected by the AAVS1 internal 5’- and external 3’-probes was calculated using the SeqBuilder program in the DNASTAR Lasergene Core Suite v12.0.0 based on the EcoRV restriction sites (one within the integrated targeting vector; one each upstream of exon 1 and downstream of exon 3 of the *PPP1R12C* gene).

### Generation of targeted transgene at *STMN2* locus in hESC via CRISPR/Cas9

*STMN2* is a neuron-specific gene, which allows for relatively neuron-specific expression of the targeted transgene from the *STMN2* locus. Site-specific genome editing via CRISPR/Cas9 was used to insert sequences coding for *SNCA* into endogenous *STMN2* gene locus.

To target the *SNCA*-*GFP* cassette into the *STMN2* locus, a plasmid was generated bearing ~1800 bp of homology surrounding the *STMN2* stop codon. An IRES-SNCA-GFP coding sequence was then cloned into the *STMN2* homologous sequence such that ~900 bp of homology flanked the IRES-SNCA-GFP cassette. A FRT flanked PGK-Neomycin cassette was then cloned between the IRES-SNCA-GFP cassette and the *STMN2* 3’ homology arm. To incorporate the cassette into the *STMN2* locus, 800,000 H9 hES cells were nucleofected using the Amaxa P3 Primary Cell 4D-Nucleofector X Kit with program CA137. The nucleofection reaction contained 15 µg of sgRNA (5’-tgtctggctgaagcaaggga-3’), 20 µg of ThermoFisher Truecut Cas9 v2 protein and 5.5 µg of the *STMN2* targeting plasmid. After the nucleofection, cells were plated in a 1:1 mixture of StemFlex (Invitrogen) and MEF conditioned StemFlex with Rock inhibitor (Peprotech). The cells were allowed to recover for 48 h before G418 selection was initiated. After visible colonies survived the selection, they were picked and plated into a 96-well plate. The expanded cells were replica-plated into two 96-well plates, one of which was used for genotyping. PCR was used to confirm the proper integration of the 5’ (primers STMN2.FOR2 and IRES-REV) and 3’ (primers NEO-F and STMN2-REV1) arms of the targeting cassette into the *STMN2* locus. After targeting confirmation, a clone was expanded and a CAG-FLPo-Puro cassette was nucleofected into the cells following the above protocol. Puromycin selection allowed for the identification of cells which expressed FLP recombinase and colonies derived from these cells were picked, expanded, and genotyped by PCR (primers STMN2.FOR2 and STMN2-REV1) to confirm removal of the PGK-Neo cassette.

### Quantitative PCR for *STMN2* expression in pi-N^SNCA-STMN2^ neurons

For RNA isolation, DIV21 neurons (see Induced Neuron Differentiation methods) were harvested from 6-well plate cultures by directly applying 1 mL Trizol (ThermoFisher, 15596018) on the cells and slowly shaking them for 10 min at room temperature. 200 µL of chloroform-isoamyl alcohol (Sigma, 25668) was added to 1 mL of Trizol extract and shaken at full speed on a thermoblock for 30 sec at room temperature, followed by 15 min 21000*g* centrifugation (table top, 4°C). The resultant aqueous phase (~400-500 μL) was recovered with PureLink RNA mini kit (ThermoFisher, 12183018A) as per manufacturer’s guidelines and final RNA was eluted with 50 µL RNAse Free water. 100 ng of RNA from each sample was reverse transcribed for cDNA production by SuperScript™ IV VILO™ Master Mix with ezDNase™ Enzyme (ThermoFisher; 11766050). Real-time qPCR measurement was performed with TaqMan Fast Advanced Master Mix (ThermoFisher; 4444557) with the following inventoried Taqman probe assays (ThermoFisher; 4331182); GAPDH: Hs02786624_g1, PGK1: Hs00943178_g1, SNCA: Hs01103383_m1, STMN2: Hs00199796_m1. The amplification was carried out on an Applied Biosciences Vii7 thermal cycler.

### Stable integration of piggyBac plasmids into iPSCs

Transfection of hiPSCs with the piggyBac constructs was carried out as follows: iPSCs were dissociated into single cells using Accutase (Invitrogen) and replated at a density of 1.5×10^6^ cells in one well of a 6-well plate coated with Matrigel (Corning). The following day, 2 μg of piggyBac construct pEXP-piB-BsD-Tet-NGN2-Puro-SNAP-PGKtk, 1.5 μg transposase pEf1α-hyPBase, and 10.5 μL TransIT-LT1 transfection reagent (Mirus) were added to 200 μL serum-free OPTI-MEM (Invitrogen). The transfection mix was incubated at room temperature for 20 min and added to cell culture containing 2 mL StemFlex medium (Invitrogen) that supports the robust expansion of feeder-free pluripotent stem cells, supplemented with 10 μM ROCK inhibitor (Peprotech). After 6 h incubation at 37°C CO_2_ incubator, the medium was changed to StemFlex plus 10 μM ROCK inhibitor. On the second day of transfection, 5 μg/mL blasticidin was added to 2 mL StemFlex plus 10 μM ROCK inhibitor. Media change was performed daily. After five days of blasticidin selection in the presence of ROCK inhibitor, cells were cultured in StemFlex without blasticidin or ROCK inhibitor until the culture became confluent. The stably transfected cell line was then ready for passaging and expansion.

### Stable integration of piggyBac plasmids into U2OS cells

U2OS cells were dissociated using 0.25% Trypsin-EDTA into single cells and replated at 1.5×10^6^ cells in a 6-well plate. On the following day, 2 µg piggyBac construct, 1.5 µg transpose pEf1a-hyPBase and 10.5 µL TransIT-LT1 transfection reagent (Mirus) were added in 200 µL serum-free OPTI-MEM (Invitrogen). The transfection mix was incubated at room temperature for 20 min and was added to the cell culture containing McCoy’s 5A medium (ATCC) supplemented with 10% FBS. After 6 h incubation in a 37°C CO_2_ incubator, the medium was changed to McCoy’s 5A media supplemented with 10% FBS. On the second day of transfection, 10 µg/mL blasticidin was added to 2 mL growth media (McCoy’s 5A media supplemented with 10% FBS) to select for transfected cells. Media was changed every 3 d. After 5 d, the stably transfected cells were passaged and expanded.

### U2OS cell culture

The U2OS cell line was purchased from ATCC. U2OS cells were maintained in McCoy’s 5A medium (ATCC) supplemented with 10% heat-inactivated Fetal Bovine Serum (FBS) and incubated in 5% CO_2_ at 37°C. U2OS cell lines with piggyBac plasmid integration were cultured in the presence of 10 µg/mL blasticidin. piggyBac transgene was induced by adding 100 ng/mL doxycycline to media. Media was changed every other day. Confluent cells (80-100% confluency) were passaged by washing cells with 1X DPBS and incubating with Trypsin (Gibco) at 37°C for 3 min. Once cells had lifted, DMEM was added and the cell suspension transferred to a Falcon tube for centrifugation at 500*g* for 5 min. The resulting cell pellet was resuspended in McCoy’s 5A, 10% FBS media, counted, and plated at the desired cell density. Mycoplasma testing was performed every other week on medium from overnight cultures. U2OS cell confluence and inclusion area (GFP) (**Figures S9A-S9B**) were tracked and quantified with Incucyte live-cell analysis instrument (Sartorius).

### iPSC generation and lines

Fibroblasts obtained from a female Contursi kindred (*SNCA* A53T) and a female Iowa kindred patient (*SNCA* triplication) were previously described in(Chung et al., 2013a; Hallacli et al., 2022). Fibroblasts from a male Iowa kindred patient (age 48) with severe early-onset parkinsonism were collected under Stanford protocol (IRB-15028) and previously described in(Byers et al., 2011; Chung et al., 2013b). Fibroblast cultures were subjected to mRNA-based reprogramming at Cellular Reprogramming Inc. using engineered and chimeric transcription factors to facilitate lineage conversion(Warren et al., 2012, 2010; Yang et al., 2015). Polyclonal pools were generated. Resulting induced pluripotent stem cell (iPSC) colonies were further expanded on rLaminin-521 (BioLamina, Sundbyberg, Sweden) in Nutristem XF media (Corning) for at least 3 passages prior to freezing; iPSCs were maintained on Matrigel in StemFlex Medium (Thermo Fisher Scientific). iPSC identity was confirmed via staining for pluripotency markers Oct4 and Tra-1-60.

Isogenic SNCA knock-down/-out controls were obtained via CRISPR/Cas9 mediated gene editing. Generation of SNCA allelic series in the female Iowa kindred patient line was described in (Hallacli et al., 2022). For generation of SNCA allelic series in the male Iowa kindred patient line, Guide4 (sequence: 5’ GCCATGGATGTATTCATGAA) targeting exon 2 was used to knock-out *SNCA*. The sequence is 16 bp downstream of Guide3, which was used for knock-down/-out of the female Iowa kindred line. The SNCA knock-down/-out lines from the male Iowa kindred patient were made using Neon transfection system to transfect ribonucleoprotein (RNP) of sgRNA and Cas9 protein with the standard protocol for iPSC transfection. The sgRNAs were purchased from Synthego and the TrueCut Cas9 protein was purchased from ThermoFisher. Genotypes were subsequently confirmed by Sanger sequencing as described in (Hallacli et al., 2022). Quality control steps for selected clones include normal karyotype, confirmation of trilineage pluripotency, and copy number variation at the Bcl2-2 via CGH array. αS protein levels were assessed across the isogenic series (4-copy, 2-copy, 0-copy) via Western blot. All *SNCA* triplication experiments in this study were performed with the male Iowa kindred patient iPSC line.

### Conventional feeder-based hESC culture

WIBR3 (clone 38)-derived hESCs were routinely cultured as colonies on a monolayer of mouse embryonic fibroblasts (MEFs) and served as starting material for conventional human cortical neuron (c-N) differentiation (**Figures S1E** left and **S1F**), as well as for immunoblotting (**Figure S1C**) and Southern blotting (**Figure S1D**).

hESCs were kept in 6-well plates at 37°C with 5% O_2_ and 3% CO_2_; cell culture and wash media were pre-warmed to 37°C before use. Primary MEFs were prepared and mitotically inactivated using mitomycin C (Sigma-Aldrich) as described previously(Conner, 2000). MEFs were plated at a density of 4 x 10^5^ cells/cm^2^ onto gelatinized cell culture plates, which were prepared by incubating with a 0.2% (w/v) gelatin solution (Sigma-Aldrich) for 1 h at 37°C. hESC cultures were supplied daily with hESC medium, and tested for mycoplasma infection every 2-4 weeks according to the manufacturer’s instructions (MycoAlert, Lonza). hESC medium was DMEM/F-12, HEPES (Thermo Fisher Scientific), supplemented with 15% (v/v) Hyclone defined fetal bovine serum (Hyclone Laboratories), 5% (v/v) Knockout serum replacement (Thermo Fisher Scientific), 1% (v/v) GlutaMAX supplement (Thermo Fisher Scientific), 1% (v/v) 100X MEM non-essential amino acids (Thermo Fisher Scientific), 1% (v/v) penicillin-streptomycin (Thermo Fisher Scientific), 0.1 mM β-mercaptoethanol (Sigma-Aldrich) and 4 ng/mL fibroblast growth factor 2 (FGF2; R&D Systems). hESCs cultures were passaged, either manually or enzymatically, when they reached 70-80% confluency. For manual passaging, colonies were cut in a grid excluding differentiated parts, using a stereomicroscope and a 26-G needle that was attached to a 1 mL syringe and bent to a 45° angle. The colony fragments were dislodged and collected in a 15 mL centrifuge tube primed with hESC medium. Colony fragments were re-plated at a 1:3-1:6 ratio onto fresh 6-well feeder plates, which were primed with hESC medium. For enzymatic passaging, differentiated parts were removed by aspiration with a glass Pasteur pipette, followed by incubation with 1.5 mg/mL collagenase type IV (Thermo Fisher Scientific) in DMEM/F-12 (Thermo Fisher Scientific) for 15-30 min at 37°C. Colonies were washed off the plate with DMEM/F-12, collected in a 15 mL centrifuge tube and broken into smaller fragments by trituration. The supernatant was aspirated, the colony fragments were washed with DMEM/F-12 twice and were reconstituted in hESC medium. hESCs were re-plated as described above.

To achieve clonal purity, hESC colonies shown in **Figure S1D** were grown from a single cell suspension plated at very low seeding density. The resultant colonies were exposed to 2 μg/mL doxycycline for 10 d (*SNCA-mK2-AAVS1*) or 21 d (*GFP-AAVS1*) from DIV10 for the induction of transgene expression. The micrographs showing transgene-driven GFP/mKate2 fluorescence in **Figure S1D** were obtained using an inverted epifluorescent microscope (Eclipse Ti, Nikon Instruments), and were visualized and processed with the NIS-Elements AR software package (Nikon). In preparation for creating a single cell suspension, routinely grown feeder-based hESC cultures were pre-incubated for 30 min at 37°C with hESC medium that was supplemented with 10 μM of the small molecule Y-27632 (Stemgent); Y-27632 was used as rho-associated protein kinase (ROCK) inhibitor (RI). After a wash step with Dulbecco’s phosphate-buffered saline (DPBS; Thermo Fisher Scientific), cells were incubated with StemPro Accutase cell dissociation reagent (“Accutase”; Thermo Fisher Scientific) for 10 min at 37°C. Accutase was diluted with hESC medium/RI, the cell suspension was collected in a 15 mL centrifuge tube and each well was washed with hESC medium/RI. Cells were centrifuged at 350*g* for 10 min, resuspended in hESC medium/RI, followed by trituration to create a single cell suspension and filtered through a 40 μm cell strainer. The single cell suspension was further diluted with hESC medium/RI and re-plated at a density of 50-2,000 cells per well of a 6-well feeder plate. ROCK inhibitor was withdrawn after 48 h.

### Conventional human cortical neuron (c-N) differentiation

For conventional neuronal differentiation (c-N^SNCA-mK2-AAVS1^ in **Figure S1E** left, and c-N^GFP-AAVS1^ in **Figure S1F**), WIBR3 (38)-derived hESC lines harboring integration of *SNCA-mK2* or *GFP* transgene, respectively, were cultured feeder-free prior to differentiation and neuralized by embryoid body (EB) formation. Neural progenitor cells were differentiated into cortical neurons of anterior forebrain identity(Chung et al., 2013b). The full protocol has been published previously(Chung et al., 2013b; Khurana et al., 2017) and results in cell cultures that are enriched for VGLUT1+ glutamatergic neurons and also contain a fraction of GFAP+ astrocytes(Chung et al., 2013b).

### Induced neuron differentiation

On day 0, iPSCs at ~95% confluency were lifted by incubating with Accutase (Life Technologies), a natural enzyme mixture with proteolytic and collagenolytic enzyme activity, for 4 mins at room temperature, combined with equal volume of StemFlex media, centrifuged at 800 rpm for 4 min, resuspended in StemFlex media, and counted. Cells were seeded at a density of 1.25 × 10^6^ cells per well (for 6-well plates) with 0.5 μg/mL doxycycline to induce expression of Ngn2 on the piggyBac transgene. For 10-cm plates, 10 million cells were seeded. This was considered day 0. Plates were previously coated with Matrigel. For the first 2 days of neuron differentiation, media change was conducted daily with Neurobasal N2/B27 media (1X B27 supplement (Life Technologies), 1X N2 supplement (Life Technologies), 1X Non-Essential Amino Acids (Gibco), 1X GlutaMAX (Gibco), 1X Pen-Strep (Gibco), Neurobasal Media (Life Technologies)), 5 μg/mL blasticidin and 0.5 μg/mL doxycycline; for days 3-6, media changes were done the same as for days 1-2, with the addition of 1 μg/mL puromycin to select cells expressing the piggyBac transgene.

On day 7, Accutase was used to dissociate the neurons before re-plating them onto the appropriate polyethyleneimine (PEI)/laminin-coated plates for downstream assays (e.g., 3 million cells per well of 6-well, 1 million cells per well of 24-well, 50,000 cells/well of 96-well plates). The following day (day 8), an equal volume of Neurobasal N2/B27 media supplemented with 20 ng/mL Brain-derived Neurotrophic Factor (BDNF; Peprotech 450-02), 20 ng/mL Glia-derived Neurotrophic Factors (GDNF; Peprotech 450-10), 2 mM Dibutyryl cyclic AMP (cAMP; Sigma D0260), 2 μg/mL laminin, 0.5 μM cytosine b-D-arabinofuranoside hydrochloride (AraC; Sigma) was added to the existing cell media. Doxycycline was withdrawn from medium on day 8. For neurons transfected with the all-in-one piggyBac transgene containing *NGN2* and *SNCA*, doxycycline supplementation was continued throughout neuronal culture to maintain αS overexpression. At day 11, media change occurred with equal volumes of Neurobasal N2/B27 and Neurobasal Plus (Life Technology A35829-01) N2/B27 Plus media, and 10 ng/mL BDNF, 10 ng/mL GDNF, 1 mM cAMP, 1 μg/mL laminin. At day 14, half media change occurred with Neurobasal Plus media, 10 ng/mL BDNF, 10 ng/mL GDNF, 1 mM cAMP, 1 μg/mL laminin. Thereafter, half media change occurred every three days with Neurobasal Plus media, 10 ng/mL BDNF, 10 ng/mL GDNF, 1 mM cAMP, 1 μg/mL laminin.

### Induced astrocyte differentiation

On day 0, H1 hESCs at ~95% confluency were dissociated with Accutase, and 4 × 10^6^ cells were replated in Matrigel-coated 10-cm dishes using StemFlex medium with 10 μM ROCK inhibitor (StemCell Technologies, Y-27632) and 500 ng/mL doxycycline to induce human NFIB epression (420 aa protein; NCBI Reference Sequence NP 005587.2). On days 1 and 2, cells were cultured in Expansion medium (DMEM/F-12, 10% FBS, 1% N2 supplement, 1% Glutamax (Thermo Fisher Scientific)). From days 3 to day 5, Expansion medium was gradually switched to FGF medium (Neurobasal, 2% B27 supplement, 1% NEAA, 1% Glutamax, and 1% FBS (Thermo Fisher Scientific); 8 ng/mL FGF, 5 ng/mL CNTF, and 10 ng/mL BMP4 (Peprotech)). On day 6, the mixed medium was replaced by FGF medium. Selection was carried out on days 1–6 with 5 μg/mL blasticidin for cell lines harboring vectors conferring blasticidin resistance. On day 7, cells were dissociated with Accutase and replated in Matrigel-coated wells. The day after, FGF medium was replaced, and afterwards 50% of the medium was replaced by Maturation medium (1:1 DMEM/F-12 and Neurobasal, 1% N2, 1% sodium pyruvate, and 1% Glutamax (Thermo Fisher Scientific); 5 µg/mL *N*-acetyl-cysteine, 500 µg/mL dbcAMP (Sigma-Aldrich); 5 ng/mL heparin-binding EGF-like growth factor, 10 ng/mL CNTF, 10 ng/mL BMP4 (Peprotech)) every 2–3 d, and cells were kept for either 8 days or 21 days.

### Recombinant aS expression and purification

Lyophilized monomeric αS was provided by Dr. Tim Bartels. Briefly, plasmid pET21a-SNCA was expressed in BL21(DE3) *E. coli*. After cell lysis, αS was purified via ion exchange chromatography (5 mL HiTrap Q HP columns, GE Life Sciences, 17516301) and size exclusion chromatography (13 mL HiPrep 26/60 Sephacryl S-200 HR, GE Life Sciences, 17119501) using the ÄKTAprime plus FPLC system (Rovere et al., 2019) and subsequently lyophilized in protein low binding tubes (Eppendorf).

### Generation of preformed αS fibrils

For the generation of wild-type recombinant preformed fibrils (PFFs), 1 mg of lyophilized monomeric wild-type αS was reconstituted with 100 µL of sterile DPBS (pre-cooled to 4°C) on ice without further resuspending. Tubes were then rotated on a tube rotator for 10 min at 4°C and subsequently centrifuged for 10 min at 15,000*g* at 4°C to pellet preformed aggregates. The supernatant was then transferred to a new protein low binding tube and concentration was measured spectrophotometrically using Nanodrop (A280 – MW=14,5 kDa; Extinction coefficient ε for human αS = 5,960 M^−1^cm^−1^). Samples were then diluted down to a final concentration of 5 mg/mL and aliquoted into 100 µL aliquots. A 1-2 mL aliquot was diluted down to 500 µg/mL for electron microscopy and flash-frozen in a dry ice/ethanol slurry. Samples were placed into an orbital thermomixer with a heated lid for 7 d at 37°C, shaking at 1,000 rpm. At the end of the 7-day period, the contents of the tube appeared turbid. The tube was gently flicked to resuspend preformed fibrils, aliquoted in 1-2 µL volumes for TEM, and flash-frozen in a dry ice/ethanol slurry prior to storage at −80°C.

A53T PFFs generated from αS-A53T monomer was kindly provided by Dr. Kelvin Luk.

### PFF seeding in iPSC-derived cortical neurons

For PFF seeding of cortical neurons, WT PFFs were used for neurons overexpressing WT αS, and A53T PFFs for αS-A53T overexpressing lines to match the amino acid sequence of intracellular overexpressed αS and PFF strain. On DIV11, a PFF aliquot and PBS aliquot (negative control) were thawed at RT for 2-3 min and subsequently transferred to an ice bucket for water bath-based sonication using the Bioruptor Plus (Settings: High power, 10 cycles, 30 sec on – 30 sec off per cycle, temperature 10°C). Sonicated samples were subsequently transferred into the tissue culture hood and diluted in culture medium to a concentration of 10 µg/ml. At DIV14, no media was removed, but instead the same amount of fresh medium was added. From DIV18, half media changes were performed according to NGN2 protocol.

### PFF seeding in U2OS cells

For seeding of U2OS cells, A53T PFFs were used for cells overexpressing αS-A53T-sfGFP or αS-A53T-ΔNAC-sfGFP to match the amino acid sequence. On Day −2, Cells were plated at 2000 cells/well in 96-well plate (Greiner) in 100 µL McCoy’s 5A medium supplemented with 10% FBS, 10 µg/mL blasticidin and 100 ng/mL doxycycline. Two days later (48 h post-induction of transgene), cells were seeded with PFFs by quick-thawing an aliquot of PFFs in water-bath and re-sonicating with Bioruptor Pico (10°C, 10 cycles, 30sec ON, 30sec OFF, HIGH power). PFFs were resuspended in media at desired concentration (0, 1, 3, 5 and 10 µg/mL) and added to cells. This was considered Day 0. On Day 3 (72 h post-seeding), an equal volume of fresh media (without PFFs) was added to the wells. On Day 6, cells were fixed in 4% PFA, 20% sucrose in PBS for immunostaining.

### Electron microscopy

For electron microscopy of PFFs, 5 µL of 500 µg/mL PFFs was adsorbed for 1 minute to a carbon coated grid (Electron Microscopy Sciences, CF400-CU) that had been made hydrophilic by a 20 sec exposure to a glow discharge (25 mA). Excess liquid was removed with a filter paper (Whatman #1), the grid was then floated briefly on a drop of water (to wash away phosphate or salt), blotted again on a filter paper and then stained with 0.75% uranyl formate (Electron Microscopy Sciences, 22451) or 1% uranyl acetate (Electron Microscopy Sciences, 22400) for 20-30 sec. After removing the excess stain with a filter paper the grids were examined in a JEOL 1200EX Transmission electron microscope or a TecnaiG² Spirit BioTWIN and images were recorded with an AMT 2k CCD camera.

For electron microscopy of induced neurons, iPSC-derived neurons were seeded at day 7 at 0.6 million cells/well on ACLAR plastic discs pre-coated with polyethyleneimine (PEI)/laminin in 12-well plate (Corning). At 25 days of differentiation, iPSC-derived neurons were fixed in 2.5% glutaraldehyde, 1.25% paraformaldehyde, 0.03% picric acid in 0.1 M sodium cacodylate buffer (pH 7.4) for 60 min. After 3 washes in 0.1M cacodylate buffer, postfixation was done in 1% osmium tetroxide (OsO4)/1.5% potassiumferrocyanide (KFeCN6) for 30 min followed by 3 washes in dH_2_O. Coverslips were then incubated in 15 aqueous uranyl acetate for 30 min, washed twice in dH_2_O and subsequently dehydrated in grades of alcohol (5 min each; 50%, 70%, 95%, 2x 100%). Cells were then embedded in TAAB Epon (Marivac Canada Inc. St. Laurent, Canada) and polymerized at 60°C for 48 h. Ultra-thin sections (about 80 nm) were cut on a Reichert Ultracut-S microtome and picked up onto copper grids. For immunogold labeling, the sections were etched using a saturated solution of sodium metaperiodate in water for 5 min at RT. Grids were then washed 3x in dH_2_O and floated on 0.1% Triton X-100 (TX-100) for 5 min at RT. Blocking was carried out using 1% BSA + 0.1% TX-100/PBS for 1 h at room temperature. Grids were incubated with an anti-GFP antibody (1:50, Abcam 6556) in 1% BSA + 0.1% TX-100/PBS overnight at 4°C. Grids were washed three times in PBS to remove unbound antibody followed by incubation with nanogold particles for 1 h at RT. Grids were washed with PBS and water, stained with lead citrate and examined in a JEOL 1200EX Transmission electron microscope (JEOL USA Inc. Peabody, MA USA) and images were recorded with an AMT 2k CCD camera.

### Whole-cell protein extraction

For cell lysis, frozen cell pellets were thawed on wet ice and resuspended in 100 µL of 1X NuPage™ LDS sample buffer (Thermo Fisher) (diluted using dH_2_O) containing protease (cOmplete™ EDTA-free Protease Inhibitor Cocktail, Sigma Aldrich) and phosphatase inhibitors (PhosSTOP phosphatase inhibitor cocktail, Sigma Aldrich). Samples were sonicated twice for 15 sec with a tip-sonicator at 40% power, keeping sample on ice and centrifuged at maximal speed for 10 min. The supernatant was then transferred to a new Eppendorf tube and the cell pellet discarded. To quantify protein concentration, 5 µL of each sample was used in Pierce™ BCA Protein Assay Kit (Thermo Fisher) according to manufacturer’s guidelines.

### Western blotting

For western blots, 30 µg of protein per sample was subjected to SDS-PAGE using NuPAGE 4-12% Bis-Tris protein gels in NuPAGE MES SDS Running Buffer, electrophoresed at 150 V for 55 min or until the protein ladder and the loading dye indicated a sufficient electrophoretic separation. Dry transfer from polyacrylamide gel to nitrocellulose membrane was conducted with the iBlot 2 Gel Transfer Device (Thermo Fisher) using preset P0 program (20V 1 min; 23V 4 min; 25V 2 min). The membrane was fixed in 4% paraformaldehyde in dH2O (to improve aS detection (Lee and Kamitani 2011 PLoS One) for 30 min at room temperature with orbital shaking and washed three times for 5 min with PBS. Membranes were blocked in Li-COR Odyssey blocking buffer (PBS) for 1 h with orbital shaking and subsequently incubated overnight in primary antibody solution, i.e. Odyssey blocking buffer (PBS), 0.1% Tween20 and the respective primary antibodies at the desired dilution, at 4°C with orbital shaking. After four washes for 5 min in 0.05% Tween-20/PBS, membranes were incubated with the secondary antibody solution, i.e. Odyssey blocking buffer (TBS), 0.1% Tween-20 and secondary antibody (IRDye® 800CW, IRDye ® 680RD, LI-COR Biosciences) at 1:10,000 dilution, at room temperature with orbital shaking and protected from light. After four washes for 5 min in 0.05% Tween-20/PBS, the blot was scanned using the Odyssey CLx Infrared Imager. For quantification, ImageStudio software was used.

For the doxycycline dose-response shown in **Figure S1C**, routinely grown WIBR3 (38)-derived hESC cultures harboring integration of wild-type *SNCA* transgene were exposed to different concentrations of doxycycline for 30 h. Control hESC samples were cultured in the absence of doxycycline. After the 30 h treatment, hESCs were harvested and lysed for immunoblotting. Media residues were washed with 4°C chilled DPBS. Fresh chilled DPBS was added and the cell monolayer was detached with a cell scraper or a micropipette. Samples were kept on ice from this point on until cell lysis. Samples were centrifuged at 400*g* for 5 min in a 4°C refrigerated bench-top microcentrifuge. The supernatant was aspirated and fresh chilled DPBS was added, followed by centrifugation as above. The supernatant was aspirated and the cell pellets were resuspended in cell lysis buffer (2 mM EDTA, 1% [v/v] 100X protease inhibitor cocktail, 2% [w/v] SDS, 50 mM Tris-HCl in deionized water [diH_2_O]; all Sigma-Aldrich). The cell lysates were boiled for 10 min at 100°C, followed by centrifugation at 10,000*g* for 10 min. The supernatant was transferred to a new Eppendorf tube and the total protein concentration of cell lysates was determined using the Pierce bicinchoninic acid (BCA) protein assay kit (Thermo Fisher Scientific) following the manufacturer’s instructions. Briefly, 0.5-1 μL cell lysate were diluted with diH_2_O to a total volume of 25 μL (1:25-1:50 dilution). 50 parts BCA reagent A were mixed with one part BCA reagent B (50:1 ratio) and 200 μL of this solution was mixed with the diluted cell lysate, 25 μL diH_2_O (as a blank control) or 25 μL of a bovine serum albumin (BSA) standard (20-500 μg/mL BSA in diH_2_O), each in triplicate. After incubation for 26 min at 37°C the absorbance of the purple-colored reaction products was measured at 562 nm using a spectrophotometer (Epoch 2 microplate reader, BioTek). The results were averaged and corrected by the blank control. The protein concentration was determined using a standard curve derived from the BSA standards. Next, the extracted proteins were separated using a one-dimensional SDS-PAGE. According to the manufacturer’s instructions (NuPAGE SDS-PAGE gel electrophoresis system, Thermo Fisher Scientific), 20 μg bulk protein were mixed with NuPAGE LDS sample buffer and NuPAGE sample reducing agent and heated to 75°C for 10 min. The protein samples were loaded onto a NuPAGE 10% Bis-Tris gel, together with a protein molecular weight standard (Precision Plus Protein Dual Color Standards, Bio-Rad). NuPAGE MES SDS was used as electrophoresis buffer and electrophoresis was conducted at 150 V until the loading dye and the molecular weight control indicated sufficient separation (40-120 min). Electrophoretic transfer to a polyvinylidene difluoride (PVDF) membrane was carried out by wet electrophoretic transfer using an electrophoresis cell (Criterion Cell, Bio-Rad). The NuPAGE gel was briefly washed in transfer buffer (Abbiotec) that was prepared according to the manufacturer’s instructions, and assembled with an activated PVDF membrane (10-second activation in methanol) and Whatman filter papers. The transfer was carried out with chilled transfer buffer at 4°C at either 60 V for 120 min for one transfer or 65 V for 150 min for two simultaneous transfers. In preparation for chemiluminescent detection, PVDF membranes were briefly washed in PBS (Boston BioProducts) and from this point on all incubation and wash steps were performed on an orbital shaker. Immunoblots were immersed in a blocking solution, consisting of 5% (w/v) nonfat milk powder (Bio-Rad) in PBS with 0.1% (v/v) Tween-20 (Sigma-Aldrich), for 40 min, followed by incubation with mouse anti-α-synuclein (BD Biosciences; 1:500) and mouse anti-GAPDH (EMD Millipore; 1:3,000) primary antibodies, diluted in blocking solution, overnight at 4°C. The next day, immunoblots were briefly washed three times with diH_2_O and three times for 10 min with PBST (PBS with 0.1% [v/v] Tween-20), and then incubated for 1 h in blocking solution, supplemented with horseradish peroxidase (HRP)-conjugated rabbit anti-mouse IgG secondary antibody (Sigma-Aldrich; 1:10,000). Immunoblots were briefly washed three times with diH_2_O and three times for 10 min with PBST. Immunoblots were exposed to SuperSignal West Pico Chemiluminescent Substrate solution (TFS) according to the manufacturer’s instructions and exposed to a film that was developed using a Kodak X-OMAT 1000A film processor.

### Immunofluorescence and microscopy

Immunofluorescence analysis was performed as follows. iPSC-derived neuron cultures grown in 96-well glass bottom plates (Brooks Life Science Systems, MGB096-1-2-LG-L) were fixed with 100 μL of 4% paraformaldehyde in PBS for 15 min. Cells were blocked and permeabilized using 10% goat serum, 0.1% Saponin (to preserve integrity of lipid-rich inclusions) in PBS for 1 h at room temperature. Primary antibody was incubated in 2% goat serum, 0.02% Saponin overnight at 4°C. Cells were washed three times with PBS, 5 min per wash, and incubated with secondary antibody in 2% goat serum, 0.02% Saponin and 0.05% Hoechst for 1 h at 37°C. Finally, cells were washed three times with PBS, 5 min per wash. Images of the immunostained cells were captured with a Nikon TiE/C2 confocal microscope.

### Immunostaining of c-N^GFP-AAVS1^

Conventionally differentiated c-N^GFP-AAVS1^ neuronal cell cultures shown in **Figure S1F** were plated at high density (500,000 to 1y10^6^ cells/cm^2^) onto poly-D-lysine (2 mg/mL, Sigma) and mouse laminin (1 mg/mL, BD Biosciences)-coated 8-well chambered cover glasses (Lab-Tek, Thermo Fisher Scientific) as described previously(Chung et al., 2013b). From DIV8, c-N^GFP-AAVS1^ were exposed to 2 μg/mL doxycycline for 3 weeks. On DIV29, cell cultures were washed in DPBS for 5 min, followed by fixation in 4% (w/v) paraformaldehyde (Electron Microscopy Sciences) in DPBS for 15 min, and washed in DPBS another three times for 5 min. Samples were permeabilized and blocked in a blocking solution, consisting of 10% (v/v) normal donkey serum (Jackson ImmunoResearch) in DPBS with 0.1% (v/v) Triton X-100 (Sigma-Aldrich), for 1 h. Conventionally differentiated c-N cell cultures typically contain a fraction of astrocytes(Chung et al., 2013b), which were visualized with immunostaining for the astroglial marker GFAP together with immunostaining for GFP, in order to visualize even low levels of transgene expression. Samples were incubated with rabbit anti-GFAP (Dako, Agilent Technologies) and mouse anti-GFP (Roche Diagnostics) primary antibodies, each diluted 1:1,000 in blocking solution, overnight at 4°C. The next day, samples were washed three times for 5 min with PBST (DPBS with 0.1% [v/v] Triton X-100), and then incubated for 1 h in blocking solution, supplemented with fluorochrome-conjugated donkey anti-rabbit and donkey anti-mouse secondary antibodies (Thermo Fisher Scientific), diluted 1:500, as well as 10 μg/mL Hoechst 33342 nuclear counterstain (Thermo Fisher Scientific). Samples were washed twice for 5 min with PBST and 8-well chambers were supplied with DPBS. Samples were imaged with a multispectral spinning disk confocal microscope (Ultraview Perkin Elmer; Zeiss Axiovert 200 inverted microscope; 100X Zeiss 1.4 NA oil immersion lens); images were visualized and processed with Volocity software package (Perkin Elmer).

### Lattice light-sheet microscopy and 3D rendering

Lattice light-sheet microscopy was performed using the lattice light-sheet mode of a custom built Multimodal Optical Scope with Adaptive Imaging Correction (MOSAIC). Neurons were plated on a 25 mm coverslip and imaged at diffraction limited resolution using a Special Optics 0.65 NA, 3.74 mm working distance water dipping objective for excitation and a Zeiss 1.0 NA, with 2.2 mm working distance water-dipping objective for detection on two Hamamatsu Orca Flash 4.0 v3 sCMOS cameras. Upon image deconvolution of the raw data files, 3D surface rendering was performed using Imaris.

### Sequential extraction of insoluble fraction from human brain

Sequential extraction from human brain was performed according to (Peng et al., 2018). Briefly, 0.5 mg of brain tissue was homogenized in high salt (HI) buffer (50 mM Tris-HCl, 750 mM NaCl, 5 mM EDTA, 10 mM NaF, pH 7.40) containing protease inhibitors. After ultracentrifugation at 100,000*g* for 30 min at 4°C, supernatant was removed, and fresh HI buffer added. The same steps were subsequently repeated with HI buffer containing 1% Triton, HI buffer with 1% Triton and 30% sucrose, HI buffer with 1% sarkosyl, and finally in PBS to resuspend the sarkosyl-insoluble fraction of the brain homogenate enriched in aggregated αS.

### Real time-quaking induced conversion (RT-QuiC) (Seeded amplification assay)

Flash-frozen MSA and PD brain tissue (500 µg, frontal cortex) was homogenized and subjected to serial extraction using detergents in increasing strength and subsequent ultracentrifugation to obtain an insoluble protein fraction containing aggregated αS as previously described (Peng et al., 2018). For the RT-QuiC reaction(Soto et al., 2002) to amplify and monitor αS aggregates, 10 µL of brain-derived seed was incubated with recombinant monomeric αS at 42°C in a BMG FLUOstar Omega plate reader to amplify amyloid αS by incorporating monomeric αS into the growing aggregate. Before each RT-QuiC experiment, lyophilized monomeric protein was dissolved in 40 mM phosphate buffer (pH=8), filtered using a 0.22 mm filter, and the concentration of recombinant protein was measured via absorbance at 280 nm using a Nanodrop One spectrophotometer. Brain-derived insoluble protein was tip-sonicated for 30 sec (1 sec off, 1 sec on) at 30% of amplitude and added to a 96 well plate with 230 mM NaCl, 0.4 mg/mL αS and a 3 mm glass bead (Millipore Sigma 1040150500). Repeated shaking (1 min incubation, 1 min double-orbital shaking at 400 rpm) disrupts the aggregates to produce an expanded population of converting units. The amyloid dye Thioflavin T was used in adjacent wells to monitor the increase in fibrillar content via fluorescence readings at 480 nm every 30 min until the signal plateaued towards the end of the amplification interval of 6 d.

### ELISA

To determine the concentration of αS after amplification using SAA, an αS ELISA protocol from MSD was used. For sulfo-tag labeling of detection antibodies, 200 μL of SOY1 antibody (1.37 mg/mL in PBS) was incubated at room temperature for 2 h with 16 μL of 3 nmol/μL MSD NHS-Sulfotag reagent (150 nmol freshly suspended in 50 μL PBS). Next, 250 μL PBS was added to antibody solutions, concentrated using Amicon ultra filter tubes (10,000 MWCO), and brought up to 500 μL PBS again. This was repeated 5 times to dilute out the tag reagent. Protein concentration was subsequently measured using BCA assay. For plate preparation, MSD Standard plates were coated with 30 µL of 200 ng filtered 2F12 (1 mg/mL) from recently filtered batches diluted in PBS and stored overnight at 4°C. Plates were then tapped out, blocked with 150 µL per well in 5% MSD blocker A in 0.05% PBS-T, sealed and placed on an orbital shaker for 1 h at room temperature. Plates were subsequently washed 5 times with 150 µL PBS-T per well, samples were added in OG-RIPA, PBS-T with 1% MSD blocker A, as well as recombinant αS at different concentration gradients in PBS-T with 1% MSD blocker A (0.5% NP-40) and incubated for 2 h at room temperature with orbital shaking. Plates were washed 5 times with 150 µL TBS-T per well prior to addition of detection antibody solution, i.e. 30 µL per well of 200 ng sulfo-tagged SOY1 antibody in PBS-T with 1% MSD blocker A. Plates were incubated for 1 h at room temperature with orbital shaking and protected from light. After 5 washes with PBS-T, 150 µL of 2X MSD reader buffer diluted in MilliQ water was added and the plate was read with the Meso Sector S 600.

### Proteinase K digest

Sarkosyl-insoluble and SAA-amplified samples were treated with 1 µg/ml of proteinase K at 37°C for 1 h in gentle shaking. The digestion was stopped by adding NuPAGE LDS sample buffer and boiling the sample at 95°C for 7 min. Samples were then loaded onto a Novex 16% Tricine gels (Invitrogen) for protein separation. After electrophoresis, gels were incubated in 20% ethanol for 5 min at RT and blotted onto iBlot 2 NC Regular Stacks (Invitrogen) using the iBlot Dry Blotting. The membrane was rinsed in ultrapure water and incubated in 4% paraformaldehyde/PBS for 30 min at room temperature. The membranes were blocked in Odyssey blocking buffer (PBS)/PBS buffer 1:1 (LI-COR) or casein buffer 0.5% (BioRad) for 1 h at room temperature. After blocking, membranes were incubated overnight at 4°C with anti-αS clone 42 (BD Biosciences). After three washes in PBS-Tween-20 0.1%, the membrane was incubated for 1 h at room temperature with the secondary antibody (goat anti-mouse IgG F(Ab)2 conjugated with horseradish peroxidase (HRP), Abcam) in blocking solution. Membranes were washed in PBS-Tween-20 0.1% and then the signal was detected using Invitrogen iBright imaging system and the Luminata Crescendo Western HRP substrate (Millipore).

### αS Triton X-100/SDS sequential extraction

Sequential extraction of αS with Triton X-100 and SDS was performed as described in (Volpicelli-Daley et al., 2014). Briefly, neurons that were seeded at 3×10^6^ cells/well in 6-well plate were rinsed twice with PBS, kept on ice, and scraped in the presence of 250 µL of 1% (vol/vol) Triton X-100/TBS with protease and phosphatase inhibitors. The lysate was transferred to polyallomar ultracentrifuge tubes and sonicated ten times at 0.5 s pulse and 10% power (Misonix Sonicator S-4000). Samples were incubated on ice for 30 min, then centrifuged at 100,000*g* at 4°C for 30 min in an ultracentrifuge. The supernatant (Triton X-100 extract) was transferred to a microcentrifuge tube and combined with 4x Laemmli buffer for SDS-PAGE (small aliquot of ~20 μL is saved prior to mixing with Laemmli buffer for protein assay). In the meantime, 250 μL of 1% Triton X-100/TBS was added to the pellet and sonicated ten times at 0.5 s pulse and 10% power, followed by ultracentrifugation at 100,000*g* at 4°C for 30 min. Next, 125 μL of 2% (wt/vol) SDS/TBS with protease and phosphatase inhibitors was added to the pellet. The sample was sonicated fifteen times at 0.5 s pulse and 10% power, ensuring that the pellet is completely dispersed. The supernatant (SDS extract) was transferred to a new microcentrifuge tube and diluted to 2x volume for the corresponding Triton X-100 fraction to make the insoluble αS species more abundant and easier to visualize by western blot. For example, 60 uL of 4x Laemmli buffer was added to 180 μL of Triton X-100 extract, and 30 uL of 4x Laemmli buffer to 90 μL SDS extract.

BCA protein assay was performed on the Triton X-100 supernatant and SDS extract. For SDS-PAGE, 5 mg of protein samples were boiled for 5 min, centrifuged for 2 min at maximum speed, and loaded onto 4-12% Bis-Tris gel. The samples were electrophoresed at 150V for approximately 90 min. Protein was transferred to PVDF membrane using iBlot 2 Dry Blotting System (Invitrogen). The membrane was fixed for 30 min in 0.4% PFA/PBS if detecting untagged αS. The membrane was subsequently blocked for 1 h with 5% (wt/vol) milk/TBS-T before incubating with primary antibody overnight at 4°C with shaking. The primary antibody was diluted in 5% (wt/vol) milk/TBS-T. The following primary antibodies were used: rabbit anti-pS129 (Abcam 51253) 1:5000, mouse anti-αS 4B12 (Thermo Fisher, Cat# MA1-90346) 1:1000, goat anti-GFP (Rockland 600-101-215) 1:5000, mouse anti-GAPDH (Thermo Fisher MA5-15738) 1:5000. After incubation with primary antibody, the membrane was rinsed three times with TBS/T, 10 min with rocking for each rinse. The membrane was then incubated with HRP-conjugated secondary antibodies for 1 h at room temperature, with rocking. The following secondary antibodies were used: anti-rat-HRP (Sigma Aldrich NA935) 1:10,000, anti-rabbit-HRP 1:10,000 (Bio-Rad 170- 6515), anti-goat-HRP 1:10,000 (R&D Systems HAF109). The membrane was rinsed three times with TBST/T, 10 min per rinse, with rocking, before developing with chemiluminescence.

### Seahorse XF Cell Mito Stress Test

iPSC-derived neurons were seeded at day 8 on polyethyleneimine (PEI)/laminin coated Seahorse assay plates (Agilent Technologies Inc.) at a density of 1×10^5^ cells/well and cultured for 12 days. For transgenic lines, half of the wells were seeded with 10 µg/mL bath-sonicated recombinant A53T PFFs on day 11. On the day of the experiment, cells were pre-incubated for 1 h in Assay Media (Seahorse XF Base Medium without Phenol Red; Agilent Technologies Inc., 103335-100) supplemented with 10 mM Glucose, 1 mM Pyruvate solution, 2 mM glutamine solution (Agilent Technologies Inc., 103577-100, 103578-100, 103579-100). Measurement of intact cellular respiration was performed using the Seahorse XF96 analyzer (Agilent) and the XF Cell Mito Stress Test Kit (Agilent Technologies Inc., 1103010-100) according to the manufacturer’s instructions. Respiration was measured under basal conditions, and in response to ATP synthase inhibitor oligomycin (2 mM) followed by the addition of the ionophore 4-(trifluoromethoxy) phenylhydrazone (FCCP; 0.75 mM) to induce maximal mitochondrial respiration. Finally, respiration was stopped by adding the mitochondrial complex I inhibitors Rotenone and Antimycin A (0.5 mM).

### Autophagic flux assay

iPSC-derived neurons were seeded at day 7 on polyethyleneimine (PEI)/laminin coated 24-well plates at a density of 1×10^6^ cells/well and cultured up to Day 56. For the autophagic flux assay, 100 nM bafilomycin (Sigma) and DMSO for control conditions was added and incubated at 37°C for 12 h prior to cell collection. Whole cell extraction, BCA analysis and western blotting were performed as described earlier (see ‘Whole-cell protein extraction’, ‘Western Blotting’), however excluding membrane fixation in 4% PFA and subsequent wash steps. Primary antibodies used included rabbit-anti-actin 1:1000 (Sigma-Aldrich Inc, Cat.# A2066), mouse-anti-p62 1:1000 (Millipore, Cat.# MPN130), rabbit-anti-LC3 1:1000 (Cell Signaling Technology, Cat.# 2775S). To determine basal LC3-II and p62 levels, bafilomycin-untreated lanes were used and normalized to actin. To assess autophagic flux, the ratio of normalized LC3-II and p62 levels between treated versus untreated samples was calculated.

### Automated longitudinal single-cell inclusion survival tracking

#### Culturing of induced inclusion neurons for live imaging with Biostation CT

Neurons were differentiated and cultured as described in “Induced neuron differentiation”. Neurons were seeded on 96-well plate at DIV7 at 50,000 cells/well. On DIV10, neurons were transduced with CAG-RFP lentivirus for sparse labeling of neurons, and on DIV11 treated with 10 µg/ml recombinant PFFs. On DIV13 live imaging in the Biostation CT (Nikon) was initiated with the following settings: 10X objective, 5 x 5 tiled images per well; 2.5 x 100 ms exposure time and 10 nm luminance for excitation at 475 for Ch2; 3 x 100 ms exposure time and 50 nm luminance for excitation at 542 for Ch 3; imaging every 6 h for 10 d.

#### Single-cell inclusion survival tracking and image analysis

All image analysis was performed at Nikon Corporation. Survival curves were plotted with the survival package library in R. To detect the morphologic properties of the inclusion neuron models in culture, morphologic masks were designed to identify and quantify the different features, including cell body size, inclusions, neurites, and intensity. After the masks for detection of each cell were optimized, the tracking was performed using the mask for cell body. The setting parameters used for this tracking are described below and summarized in (**Figure S4D**).

##### Cell and inclusion detection in seeded inclusion model

Detection and tracking of inclusion-positive neurons was based on GFP fluorescence. Whereas bright, discrete GFP signal was detected in inclusion-positive neurons, the GFP signal in inclusion-negative neurons was too diffuse for accurate longitudinal single-cell tracking, so inclusion-negative neurons were tracked by RFP fluorescence. For inclusion-positive cells, objects were identified as cell bodies if GFP fluorescence intensity was greater than 60 and length and width were 6–30 pixels. Areas with fluorescent intensities that were the mean fluorescent intensity of the cell body plus 70 or higher were identified as inclusions. Inclusion GFP intensity threshold was a flexible threshold in that it changes for each cell and each frame since it relied on the mean fluorescence intensity of the cell body for the specific cell at the specific frame. If the ratio of inclusion area to cell body area was above 0.015, these cells were tracked as inclusion-positive cells. For survival analysis, a secondary filtering step was implemented to the tracked inclusion-positive cells to eliminate rounded dying cells with bright GFP signal that were incorrectly identified as inclusion-positive cells. The secondary filter consisted of an upper threshold of inclusion area in the soma and mean GFP fluorescence intensity. Tracked neurons with ratio of inclusion area to cell body area of less than 0.4 and cell body mean GFP fluorescent intensity below 150 were used for the survival analysis. For inclusion-negative cells, objects were detected as cell bodies if RFP fluorescence intensity was greater than 250 and length and width were 10–30 pixels.

##### Cell and inclusion detection in spontaneous inclusion model

Detection and tracking of inclusion-positive neurons was based on GFP fluorescence, whereas that of inclusion-negative neurons was based on RFP fluorescence. For inclusion-positive cells, objects were identified as cell bodies if GFP fluorescence intensity was greater than 40 and length and width were 15–30 pixels. Areas within cell bodies with fluorescent intensities of 160 or higher were identified as inclusions. If the ratio of inclusion area to cell body area was above 0.02, these cells were tracked as inclusion-positive cells. For inclusion-negative cells, objects were detected as cell bodies if RFP fluorescence intensity was greater than 180 and length and width were 10–40 pixels.

##### Single-cell tracking and live/dead identification

To track individual cells, a tentative cell identification (cell ID) number was assigned to each identified cell body at the beginning of the analysis. Neurons were tracking targets if they were detected at the analysis starting frame and were tracked for 5 frames or more. When cell area decreased by 50% compared to the previous frame or the fluorescence intensity decreased below the threshold, the cell was considered dead. The matching of individual cells between two time-frames was performed based on distance between each cell and changes in cell size and mean intensity. If the tracked cell merged with other cells, the tracking result was regarded as inaccurate, and the tentative cell ID was deleted. The final tracking ID number was renumbered automatically after tracking analysis.

##### Detection of neuritic inclusions in seeded inclusion model

Neuritic inclusions in the seeded inclusion model were detected and measured using GFP fluorescence. Objects above length and fluorescence thresholds except for cell bodies were recognized as neuritic inclusions. Objects that were 20–30 pixels long with GFP fluorescence intensity above 80 were identified as short and thick neuritic inclusions. Objects more than 30 pixels long and with GFP fluorescent intensity above 60 were recognized as long neuritic inclusions. Measurement of neuritic inclusions was performed per well (population-based), instead of single-cell, since it was not always feasible to assign a neuritic inclusion to a cell soma (especially over long distances).

### Lipidspot live-cell staining and manual quantification

For live cell staining, LipidSpot™ 610 dye (Biotium) was added at 1:1000 dilution on DIV11 of NGN2 transdifferentiation together with preformed fibrils and twice every week to maintain lipid stain. For manual quantification, image frames were assessed using the CL-Quant software and findings were validated by an additional independent observer.

### Live-cell compound treatments and imaging

For live-cell imaging, neurons were pretreated with LipidSpot™ 610 dye (Biotium) at 1:1000 dilution to visualize lipid-rich inclusions and imaged using an encoded stage on the Nikon TiE fluorescence microscope with a 20X Plan Apo dry objective and Andor Zyla 4.2P sCMOS camera (No binning, 200 MHz readout rate, 12 bit & Gain4 dynamic range). After 2 h, trifluoperazine and nortriptyline were added at the respective concentrations and images were acquired every 60 min for 4 h and every 24 h thereafter.

### Immunohistochemistry in post-mortem brain

This study has used immunofluorescence in six sporadic PD cases (mean age 80 (range 71-88) years old, five males and one female, mean post-mortem delay 18 (range 12-33) h), two A53T cases (48 and 54 years old, both males, post-mortem delay of 27 and 48 h), and three non-neuropathological control cases (mean age 91 (range 89-93) years old, two males and one female, mean post-mortem delay 33 (range 24-46) h). Formalin-fixed paraffin-embedded (FFPE) sections of the cingulate cortex were cut at 8 µm with a rotary microtome (Thermo/Microm, HM325) and mounted on Series 2 adhesive microscope slides (Trajan Scientific Medical, AU). Sections were de-waxed and rehydrated with xylene and a series of graded ethanol before staining. Each antibody was tested with peroxidase immunohistochemistry to determine the optimal heat-induced antigen retrieval (HIAR, performed in a programmable antigen retrieval cooker (Aptum Bio Retriever 2100, Aptum Biologics Ltd, UK)). Tests showed that TE buffer (pH 9.0) was best for RhoA, citrate buffer (pH 6.0) was best for Rab8, citraconic anhydride buffer (0.05%, pH7.4) was best for Beta-III Tubulin, and additional formic acid treatment (70% concentration for 30 min) was required for αS. To eliminate autofluorescence, sections were immersed in PBS with 0.1% sodium borohydride for 30 min on ice, washed in PBS, incubated with 100 mM glycine in PBS for 30 min; then immersed in 0.1% Sudan Black in 70% ethanol for another 30 min. After treatment with blocking buffer (containing 2% serum and 1% bovine serum albumin in PBS) for 1 h at room temperature, sections were incubated with the cocktail of primary antibodies (rabbit anti-pS129 (Abcam, Cat.# ab51253), mouse anti-Rab8 (BD Biosciences, Cat.# 610844), mouse anti-p62 Lck ligand (BD Biosciences, Cat.# 610833), mouse anti-αS (BD Bioscience, Cat.# 610787), mouse anti-RhoA (Santa Cruz, Cat.# sc-166399), goat anti-αS (R&D, Cat.# AF1338), BODIPY 493/503 (Thermo Fisher, Cat.# D3922), mouse anti-b-tubulin III (Stemcell Technologies, Cat.# 60052)) in blocking buffer at the appropriate dilutions for 48 h at 4°C, followed by their corresponding Alexa Fluor secondary antibodies and 4′,6-diamidino-2-phenylindole (DAPI, Sigma cat# D9542, 1 mg/mL) for 2 h at room temperature. To further quench autofluorescence, the fluorophore-labelled slides were finally treated with 10 mM CuSO_4_ in 50 mM ammonium acetate buffer (pH 5.0) for 1 h before coverslipping with mounting medium (DAKO, cat# S3023) and sealing with nail polish. Negative controls were performed for each batch of staining by omitting either the primary or secondary antibodies and using control case sections. Images were captured using a confocal microscope (Nikon C2) with parameters (laser power, gain, and offset) set based on the negative control and single channel labelling.

### Correlative light- and electron microscopy (CLEM) in iPSC-derived neurons and postmortem brain

Correlative light and electron microscopy (CLEM) was performed as follows. iPSC-derived cortical neurons were seeded at day 7 at 0.6 million cells/well on ACLAR plastic discs coated with polyethyleneimine (PEI)/laminin in a 12-well plate (Corning). At 25 days of differentiation, iPSC-derived neurons were fixed in 0.1% glutaraldehyde, 4% EM grade paraformaldehyde, in 0.1 M cacodylate buffer in water for 1 h at room temperature. The coverslips were further processed by washing three times with 0.1M cacodylate buffer, incubating in 1% osmium tetroxide (OsO4)/1.5% potassium ferrocyanide (KFeCN6) for 30 min, followed by a series of three washes and 30 min incubation in 1% aqueous uranyl acetate. Water was used to wash the neurons twice and they were dehydrated in 50%, 70%, 95% and twice in 100% alcohol. Neurons were embedded at 60°C for 2 d in TAAB Epon and serial 150 nm ultramicrotome sections were collected alternating on glass slides and EM grids. The glass slides were immunostained as described in (Shahmoradian et al., 2019) using the pS129 αS antibody 11a5 (Prothena) at a 1:100,000 dilution detected with HRP-green (42lifesciences). Light microscopy images were taken on a THUNDER 3D tissue imager (Leica). Immediately adjacent tissue sections mounted onto electron microscopy grids were imaged on a Philips CM100 transmission electron microscope at 80kV equipped with a TVIPS F416 camera. For A53T postmortem brain tissue, 60 µm vibratome sections post-fixed in 2.5% PFA and 2% glutaraldehyde were used. Immunohistochemistry was performed using the full-length αS antibody LB509 clone 42 (BD Biosciences) after formic acid pre-treatment in Tris-EDTA, pH 9.0, for 30 min at 95°C and a 1:100 dilution for 4 h at room temperature.

### Membrane Yeast Two-Hybrid

Membrane yeast two-hybrid (MYTH) to test protein interactions with αS was conducted in two rounds, with 2 technical replicates per round. Interactions between 20 pairs of prey-bait proteins were included as positive controls (EGFR or ATP13A2 as prey with 10 bait proteins each), and 188 prey-bait pairs were included as random controls in the second round. A total of 776 proteins were tested for interaction with αS.

The MYTH method was as described in (Chung et al., 2017). Briefly, host yeast strain NMY51 (Dualsystems Biotech AG; genotype *MAT*a *his3delta200 trp1-901 leu2-3,112 ade2 LYS2::(lexAop)4-HIS3 ura3::(lexAop)8-lacZ (lexAop)8-ADE2 GAL4*) was transformed with bait vector pGBT3-STE harboring TF-Cub-Bait and prey vector pGPR3-N harboring Nub-Prey. Control construct (either Nubi-Ost1 or NubG-Ost1) was initially co-transformed with the bait vector to serve as one positive or negative control, respectively, to calibrate and optimize the assay conditions. Yeast harboring both bait and prey plasmids exhibited growth on synthetic complete (SC) medium depleted of Leu and Trp (SC-Leu-Trp). Protein interactions between bait and prey were detected by growth on SC-Leu-Trp-His supplemented with 10 mM of 3-amino-12,4-triazole (3AT) to test for *HIS3* reporter expression. Yeast growth phenotypes were scored 1-4 based on the size of the growth spot. Protein interactions were considered positive if both repeats got a score ≥1 (lenient interaction call), or if both scores were >1, or at least one score was 4 and the other ≥1 (stringent interaction call).

### Proximity ligation assay (PLA)

For the PLA assay, the Duolink™ In Situ Orange Starter Kit Mouse/Rabbit (Sigma, DUO92102) was used. For permeabilization, 1X Saponin in PBS was incubated for 1 h at room temperature prior to blocking in Duolink™ Blocking Solution (80 μL/well; vortex well before use) for 1 h at 37°C. Primary antibodies were diluted in the Duolink™ Antibody Diluent corresponding to IF concentrations. For PLA negative controls, only one or no primary antibody was added. The plate was sealed tight with aluminum tape to prevent dehydration and incubated at 4°C overnight. Wash Buffers A and B were prepared to 1X concentration according to Duolink™ manufacturer protocol. In short, each wash buffer pouch was dissolved in 1 L autoclaved MilliQ water. The wash buffers were stored at 4°C shielded from light and were brought to room temperature before use. Following overnight primary antibody incubation, neurons were washed twice with 1X Wash Buffer A (200 μL/well) at room temperature for 5 min each. PLUS and MINUS PLA Probes were diluted 1:5 in pre-vortexed Duolink™ Antibody Diluent accordingly and incubated in the diluted PLA probe solution for 1 h at 37°C, followed by two subsequent washes with 1X Wash Buffer A at room temperature. 5X Ligation Buffer was thawed and diluted to 1X with Ultrapure distilled water (Invitrogen: 10977-015) immediately before use and Ligase was added to the freshly made 1X Ligation Buffer at 1:40 dilution. The neurons were incubated in the ligase solution for 30 min at 37°C. Following the ligation step, the neurons were washed twice with 1X Wash Buffer A at room temperature. 5X Amplification Buffer was thawed and diluted to 1X with UltraPure distilled water immediately before use, and Polymerase was added to the freshly made 1X Ligation Buffer at 1:80 dilution. The neurons were incubated in the ligase solution for 100 min at 37°C. After two washes with 1X Wash Buffer B (200 μL/well) at room temperature for 10 min each and an additional wash with 0.01X Wash Buffer B (diluted in UltraPure distilled water) at room temperature for 2 min, neurons were incubated with PBS-Hoechst solution (2000x fold dilution to PBS from commercial stock, Hoechst 33342, Invitrogen) at room temperature for 10 min, and further washed with 1X DPBS twice for 5 min each. The plate was sealed with aluminum tape and imaged immediately.

### Genome-wide CRISPR/Cas9 screen in U2OS cells

A library of sgRNAs targeting 18,166 genes with 5 gRNAs per gene for a total of approximately 91,000 gRNAs was synthesized by oligo array and cloned into the lentiCRISPR V2 puro vector (Addgene plasmid # 52961). Sequences of sgRNAs were chosen based on performance in previous screens which identified essential genes (Wang et al., 2015). Cloning of the library was performed as follows. Oligo array synthesized oligos were PCR amplified and digested with BbsI. Digests were run on a 10% TBE gel and the 28 bp sgRNA band excised and gel purified. Digested sgRNAs were ligated into BsmBI digested lentiCRISPR V2 puro and ligated overnight at 16°C (250 ng digested lentiCRISPR V2, 1 µL Invitrogen T4 ligase, 4 µL 5X ligase buffer, 0.5 or 1 or 2 µL insert, to 20 µL with H_2_O). Ligated library was precipitated for 1 h at −80°C (20 µL ligation reaction + 2 µL 3 M sodium acetate pH 5.2, 0.6 µL 2 mg/mL glycogen, 50 µL 100% ethanol). Precipitated ligation product was resuspended in 5 µL of water and used to electroporate DH10beta bacteria (1 µL per 25 µL DH10beta electroporation reaction). Electroporated bacteria were grown in 1 mL SOC at 37°C for 1 h and then plated onto LB + ampicillin plates (500 µL of transformation per plate). Transformation efficiency was determined by counting colonies on dilution plates. Transformation should result in sufficient colonies for 20-30X coverage of the library (2-3 million colonies). Colonies were scraped into a 500 mL LB + ampicillin liquid culture and grown overnight at 30°C for pooled library maxiprep (Invitrogen, 2 columns per 500 mL culture). Pooled virus was prepared by transfecting 293T cells with the library plasmid pool with psPax2 and pMD2.G lentiviral packaging vectors. Viral supernatants were harvested at 48 and 72 h post transfection and concentrated with lenti-X concentrator solution (Clontech).

To determine the amount of virus to use in the infection, a titering experiment was performed in which 10-fold serial dilutions of the virus (10 µL, 1 µL, 0.1 µL, 0.01 µL, 0.001 µL) were used to infect cells seeded in a 6-well plate at the same seeding density as a 15-cm plate (i.e., 17-fold fewer cells based on the surface area difference between a 6-well plate and a 15-cm plate). Growth media supplemented with 2 µg/mL puromycin was added 1 d after infection and selection proceeded until the uninfected well was completely dead. The amount of virus resulting in 60-80% killing was recorded. This virus amount translates to a multiplicity of infection (MOI) of around 0.2-0.3 for the screen. The number of cells needed for the start of the screen depends on the size of the library to be screened. For a library of 40,000 gRNAs and a representation of 500 cells/gRNA, 20 million cells are required per replicate. For screening in triplicate, this means that 60 million cells are required. A low MOI is used to ensure that there is only 1 gRNA per cell, thus 3-5 times as many cells as virus are required. Taken together, a library of 40,000 gRNAs at a representation of 500 in triplicate requires 180-300 million cells at the start of the screen.

U2OS cell lines were expanded to 27×15-cm plates/line at 10 million cells/plate for a total of 270 million cells for the start of the screen. U2OS cells were passaged by washing adherent cells with 1X DPBS, incubating with Trypsin for 5 min at 37°C, centrifuging for 5 min at 300 g, aspirating the supernatant, resuspending the cell pellet in growth media (McCoy’s 5A, 10% fetal bovine serum (FBS), penicillin-streptomycin), and plating at the desired cell density. Cells were infected with a gRNA/Cas9 lentivirus library at low MOI (0.2) with a representation of 500 cells/gRNA in triplicate, followed by puromycin (2 µg/mL) selection for 1 week, or until an uninfected control plate completely died. An initial cell pellet (50 million cells) was harvested as day 0 after expansion of the puromycin-selected cells to the appropriate scale to begin the screen (100 million cells/line). The remaining 50 million cells were re-plated and treated with doxycycline (100 ng/mL) to induce aS. Cell pellets were harvested 7 d (7 population doublings) and 14 d (14 population doublings) after doxycycline induction.

Genomic DNA was isolated from the day 0, 7, 14 cell pellets by phenol:chloroform extraction. Briefly, the cell pellet is resuspended in TE (10 mM Tris pH 8.0, 10 mM EDTA) to a final concentration of 2-10 million cells/mL of TE and combined with 0.5% SDS and 0.5 mg/mL Proteinase K. The suspension was incubated at 55°C overnight, with shaking/inverting the cell suspension over the course of 1 h to ensure complete digestion. Next, 0.2 M NaCl was added, followed by phenol chloroform extraction in phase lock gel tubes. Equal parts of phenol:chloroform and sample were mixed in phase lock gel tubes, shaken for 1 min to extract, then centrifuged for 5 min. The DNA aqueous phase (top layer) was subsequently chloroform extracted by mixing equal parts with chloroform, shaken for 1 min, and centrifuged for 5 min. The tubes were incubated with the caps open for 1 h at 50°C to evaporate the chloroform. Samples were treated with 25 µg/mL RNase A overnight at 37°C, then extracted with phenol:chloroform and chloroform as described above. DNA was precipitated with ethanol overnight at −20°C, or for 3 h at −80°C. Next, 1/10 v/v 3M sodium acetate pH 5.2 and 2 volumes 100% ethanol was added and the mixture centrifuged for 30-45 min at 4500 rpm at 4°C. The DNA pellet was washed once with 70% ethanol and transferred to an Eppendorf tube, followed by two more washes with 70% ethanol. The DNA pellet was dried at 37°C for 10-20 min, then resuspended in 1 mL EB/TE by incubating at 55°C. gRNAs were PCR amplified with barcoded primers for sequencing on an Illumina NextSeq 500. Sequencing reads were aligned to the initial library and counts were obtained for each gRNA. MAGeCK and edgeR were used to calculate p-values, FDRs, and log_2_ fold changes for comparison between the day 14 and day 0 samples for each cell line.

A list of 147 top hits from the CRISPR screen was compiled by applying the following thresholds in MAGeCK and edgeR analyses: log2 fold-change <= −1 or log2 fold-change >= 1 and FDR <= 0.1 for pairwise comparisons between SNCA-3K-sfGFP, SNCA-WT-sfGFP, and sfGFP fold-change. For the heatmap in **Figure S9B**, arbitrary cutoffs were used to categorize the data. The following cutoffs were used for enhancers of SNCA-3K-sfGFP: genes with log2 fold-change differential between SNCA-3K-sfGFP and SNCA-WT-sfGFP < −0.7 (dropout in 3K condition), SNCA-3K-sfGFP FDR < 0.25 for significance cutoff, and log2 fold-change for WT < 1 (to avoid genes that make WT cells grow better). The following cutoffs were used for suppressors of SNCA-3K-sfGFP: log2 fold-change differential between SNCA-3K-sfGFP and SNCA-WT-sfGFP > 1 to indicate more enrichment in SNCA-3K-sfGFP condition, and SNCA-3K-sfGFP FDR < 0.2 for significance cutoff. The list of 147 top hits was used in the Panther Gene Ontology (GO) enrichment analysis.

### ROS/MAP differential expression analysis

#### ROS/MAP study participants

Postmortem data analyzed in this study were gathered as part of the Religious Orders Study and Memory and Aging Project (ROS/MAP)(Bennett et al., 2012, 2011; Jager et al., 2018), two longitudinal cohort studies of the elderly, one from across the United States and the other from the greater Chicago area. All subjects were recruited free of dementia (mean age at entry=78±8.7 (SD) years) and signed an Anatomical Gift Act allowing for brain autopsy at time of death. Written informed consent was obtained from all ROS/MAP participants and study protocols were approved by the Rush University Institutional Review Board.

#### Study participants neuropathological evaluation

All subjects’ brains were examined postmortem by a board-certified neuropathologist blinded to clinical data. Brains were removed in a standard fashion as previously described(Bennett et al., 2006). Each brain was cut into 1 cm coronal slabs. Slabs from one hemisphere, and slabs from the other hemisphere not designated for rapid freezing, were fixed for at least three days in 4% paraformaldehyde. Tissue blocks from eight brain regions were processed, embedded in paraffin, cut into either 6 micron or 20 μm sections, and mounted on glass slides. Lewy body pathology was measured by immunostaining, as described(Schneider et al., 2012). Briefly, Lewy bodies in substantia nigra and cortex were separately identified, with only intracytoplasmic Lewy bodies indicating positive staining. To simplify the staging of pathology, the McKeith criteria(McKeith et al., 1996) were modified such that nigral predominant Lewy body pathology included cases with Lewy bodies in the substantia nigra without evidence of Lewy bodies in the limbic or neocortical regions. Limbic-type Lewy body disease included cases with either anterior cingulate or entorhinal positivity without neocortical Lewy body pathology. Neocortical-type Lewy body pathology required Lewy bodies in either midfrontal, temporal, or inferior parietal cortex with either nigral or limbic positivity, but often with both. Based on this, each subject was assigned to one of four mutually exclusive categories: 0 = no, 1 = nigral-predominant, 2 = limbic-type or 3 = neocortical-type Lewy body pathology. Further descriptions of clinical and pathological outcomes are available at the Rush Alzheimer’s Disease Centre Research Resource Sharing Hub (https://www.radc.rush.edu).

#### RNA sequencing data processing and quality control

The pipeline for sequencing has been described previously(Felsky et al., 2022). RNA sequencing on DLPFC tissue was carried out in 13 batches within three distinct library preparation and sequencing pipelines. All samples were extracted using Qiagen’s miRNeasy mini kit (cat. no. 217004) and the RNase free DNase Set (cat. no. 79254), and quantified by Nanodrop and quality was evaluated by Agilent Bioanalyzer. Full details on these methods are available on the AMP-AD knowledge portal (syn3219045). Briefly, for pipeline #1, The Broad Institutes’s Genomics Platform performed RNA-Seq library preparation using the strand specific dUTP method(Levin et al., 2010) with poly-A selection(Adiconis et al., 2013). Sequencing was performed on the Illumina HiSeq with 101bp paired-end reads and achieved coverage of 150M reads of the first 12 samples. The remaining samples were sequenced with coverage of 50M reads. For pipeline #2, RNA sequencing libraries were prepared using the KAPA Stranded RNA-Seq Kit with RiboErase (kapabiosystems) in accordance with the manufacturer’s instructions. Sequencing was performed on the Illumina NovaSeq6000 using 2 x 100bp cycles targeting 30 million reads per sample. For pipeline #3, RNA was extracted using Chemagic RNA tissue kits (Perkin Elmer, CMG-1212) on a Chemagic 360 instrument. 500 ng total RNA was used as input for sequencing library generation and rRNA was depleted with RiboGold (Illumina, 20020599). A Zephyr G3 NGS workstation (Perkin Elmer) was utilized to generate TruSeq stranded sequencing libraries (Illumina, 20020599). Sequencing was performed on a NovaSeq 6000 (Illumina) at 40-50M reads (2×150 bp paired end).

#### Differential expression analysis

All analyses were performed in R(R-Core-Team, 2014) (v3.6.3) using the same process as described(Felsky et al., 2022). Prior to calculating differential expression, a brain cell type-corrected expression matrix was generated for DLPFC on voom-transformed expression values. This matrix was used as input for differential expression analyses. Cell type proportions were estimated using the Brain Cell Type Specific Gene Expression Analysis (BRETIGEA)(McKenzie et al., 2018) package in R. Human marker genes (n=50 per set) from (Darmanis et al., 2015) were used in the application of a validated singular value decomposition method (Chikina et al., 2015) (the “adjustBrainCells” function). The limma package ‘lmFit’ function was then used to model cell type-corrected log2(expected counts) as a linear function of Lewy body pathological stage, including batch, study, biological sex, age at death, PMI, median coefficient of variation for coverage values of the 1000 most highly expressed genes, % of aligned bases mapping to ribosomal RNA, % coding bases, % UTR bases, log(estimated library size), log(passed filter aligned reads), median 5’ to 3’ bias, % of passed filter reads aligned, % read duplicates, median 3’ bias, and % of intergenic bases as covariates. Robust linear modeling was used (using Huber M estimators (Huber, 1981)), allowing for a high number (10,000) of iterations to reach convergence. Significance for effects of pathological variables on gene expression were determined using empirical Bayes moderation.

#### Rank-based enrichment of U2OS CRISPR/Cas9 screen and MYTH prioritized gene sets

The full set of gene-wise differential expression summary statistics were ranked based on moderated t-statistics. AUC-based enrichment was then performed using the R ‘tmod’ package (https://cran.r-project.org/package=tmod) to test for enrichment of three custom gene sets derived from functional screening experiments described above, selecting those which were present in the DLPFC transcriptomic data following quality control (MYTH n_genes_=269; CRISPR n_genes_=138; combined n_genes_=401). An AUC greater than 0.5 indicates a GO group enriched for higher expression with increasing Lewy body pathology, and an AUC less than 0.5 indicates a group enriched for lower expression with increasing Lewy body pathology. P-values in these comparisons are two-sided and uncorrected, corresponding to the AUC-based test for each gene set independently.

### Quantification and statistical analysis

Unless otherwise stated, plots were generated by GraphPad Prism or R. Unless otherwise noted, statistical tests were done in GraphPad Prism or R. Unpaired t-test was used to compare two groups. For multiple t-test comparisons, multiple hypothesis corrections were performed as indicated in the respective figure legends. Log-rank test was used to compare the survival distributions of two groups. For image analysis, FIJI distribution of ImageJ, CL-Quant, and ImageStudio software was used. For quantification of aS pS129 area, a custom macro was written.

## Figure Legends

**Supplementary Figure 1:**
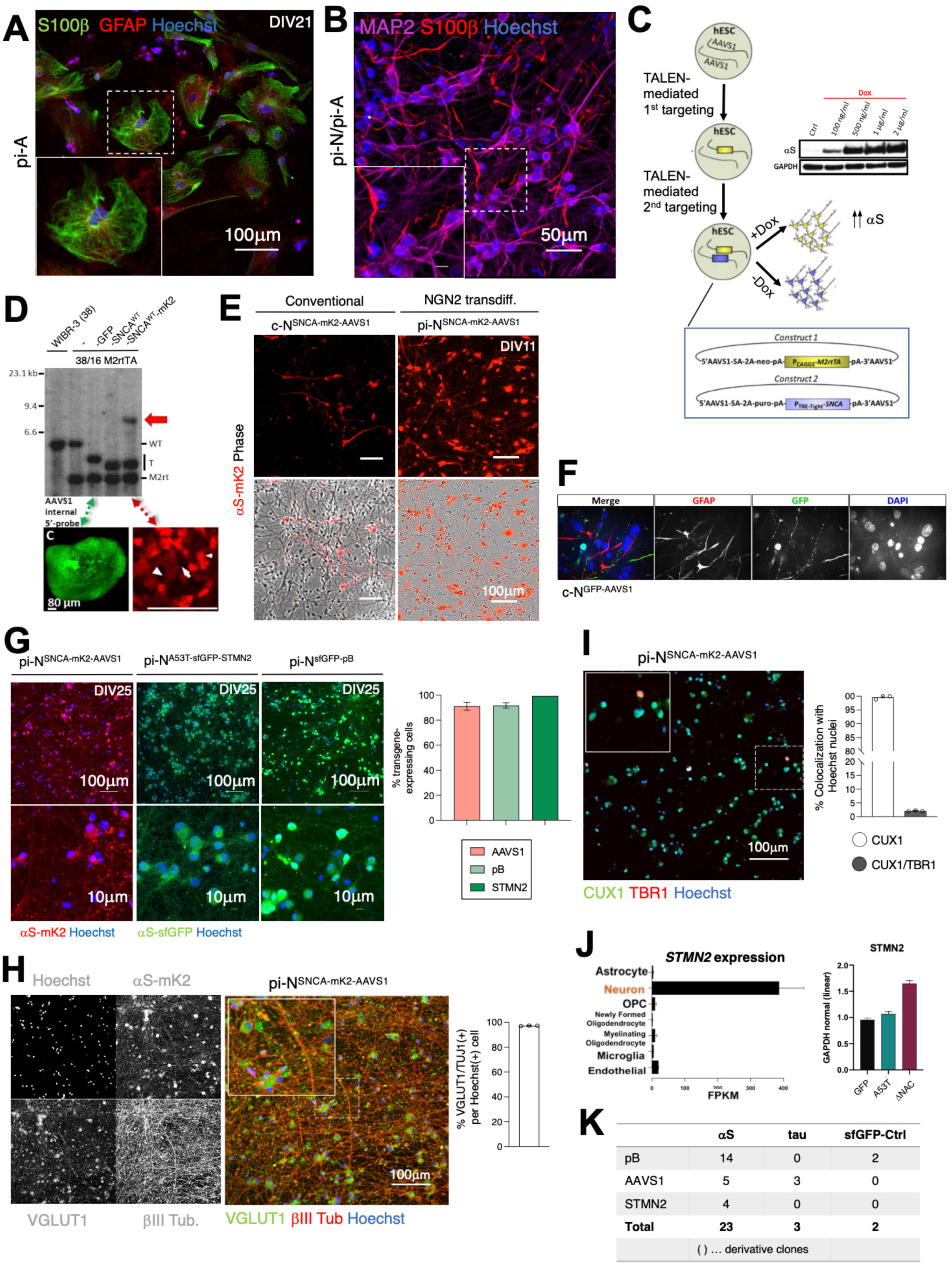
Generation of targeted inducible *SNCA* transgene at Safe Harbor (*AAVS1*) locus. (A) IF of H1/pi-As at DIV21 immunopositive for S100β(+) (green), GFAP(+) (red), and Hoechst(+) (blue). (B) Neuron-astrocyte coculture of CORR/pi-Ns (DIV25) and H1/pi-As labelled with MAP2 (purple), S100β(+) (red) and Hoechst(+) (blue). (C) Schematic of TALEN-mediated gene editing in WIBR-3, clone 38 hESC line with two constructs to allow doxycycline (Dox)-dependent regulation, one containing the *M2rtTA* reverse tetracycline transactivator, and the second construct containing the P_TRE-Tight_ *-SNCA* transgene (TRE, tetracycline response element). Treatment with doxycycline results in αS overexpression. Drawbacks of this system include transgene silencing during conventional differentiation and poor glial expression. Inset: Western blot showing doxycycline dose-response of transgene expression in hESCs harboring integration of wild-type (WT) *SNCA* transgene at *AAVS1* locus. Ctrl, control sample (no Dox). (D) Example of Southern blot analysis to confirm integration of P_TRE-TIGHT_-*GFP* (4.2 kb), −*SNCA* (4.09 kb), and −*SNCA-mK2* (4.11 kb) constructs at *AAVS1* locus in single targeted P_CAGGS_-*M2rtTA* (3.6 kb) hESC lines, compared to the untargeted alleles (5.4 kb) of the wild-type parental WIBR-3, clone 38. Red solid arrow: extra integration site. Micrographs show examples of GFP(+) and mK2(+) hESCs. White small arrowhead: very bright mK2(+) hESC; large arrowhead and arrow: faintly mK2-positive and mK2-negative hESC, respectively. Coexistence of bright and faint mK2(+) cells may be due to extra integration in some cells. T, correctly targeted allele. (E) Comparison of *SNCA-mK2-AAVS1* transgene expression efficiency via conventional differentiation or NGN2 transdifferentiation, with 1-week exposure to doxycycline, shows higher percentage of transgene silencing in conventionally differentiated neurons. (F) c-N^GFP-AAVS1^ differentiated via the spin EB method for neural induction and exposed to doxycycline from DIV8 for 3 weeks demonstrate lack of co-localization of GFP with astroglial marker GFAP. (G) Comparison of transgene expression efficiency by targeting to neuron-specific locus *STMN2*, safe harbor locus *AAVS1*, and pB random integration in pi-Ns. Right: Quantification of percent transgene-expressing cells from IF of NGN2 transdifferentiation. (H) IF staining and quantification of transdifferentiated neurons generated from hESC with targeted *SNCA-mK2* transgene at *AAVS1* locus. VGLUT1 (green) and TUJ1 (red) signal confirm glutamatergic neuron identity. (I) IF staining and quantification of CUX1 (green) and TBR1 (red) confirm superficial cortical neuron identity (layer II/III). (J) Left: *STMN2* mRNA expression is neuron-specific (source: Brain RNA-Seq browser). Right: SNCA-GFP transgene integration (with A53T or DNAC mutations) at *STMN2* locus does not alter *STMN2* expression as measured by qPCR. (K) Summary table of number of iPSC line with pB integration, *AAVS1,* or *STMN2* targeted integration, and the proteotoxic transgene that was targeted.

**Supplementary Figure 2:**
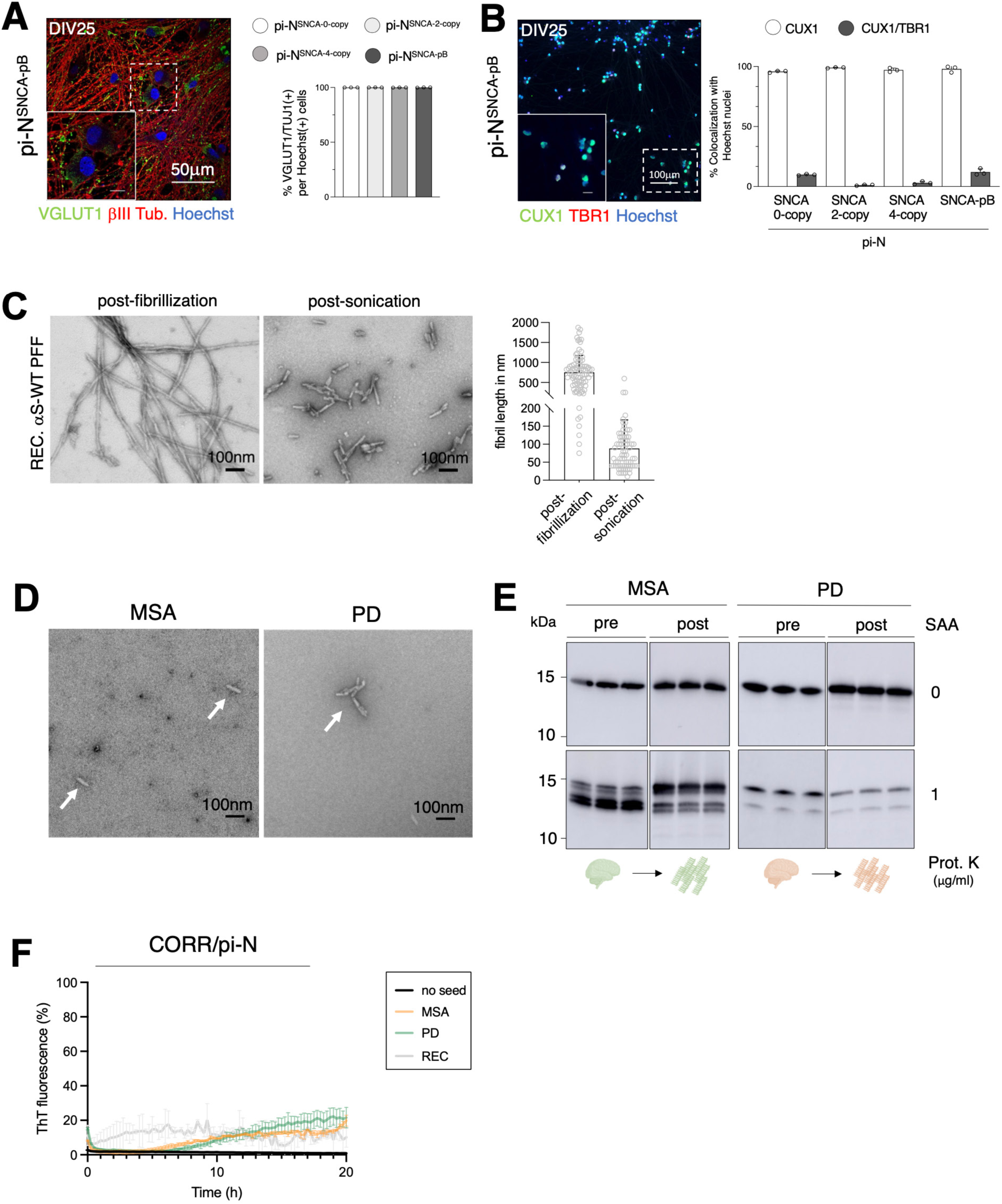
Analysis of αS PFFs used in seeding experiments. (A) IF and quantification of VGLUT1 and TUJ1 in pi-N-SNCA^4/2/0-copy^ and pi-N^SNCA-pB^ models confirm cortical glutamatergic neuron identity. (B) IF and quantification of pi-N-SNCA^4/2/0-copy^ and pi-N^SNCA-pB^ for CUX and TBR1 confirms cortical neuronal identity (layer II/III). (C) Negative staining of recombinant wild-type αS PFFs post-fibrillization (but pre-sonication), and post-sonication. Right: Quantification of fibril length. (D) Electron microscopy of MSA and PD brain-derived PFFs post- SAA. (E) Proteinase K digestion (1µg/ml) of MSA and PD brain-derived material pre- and post-SAA. (F) SAA re-amplification of CORR/pi-N neurons exposed to MSA, PD, and recombinant PFFs for 14 days.

**Supplementary Figure 3:**
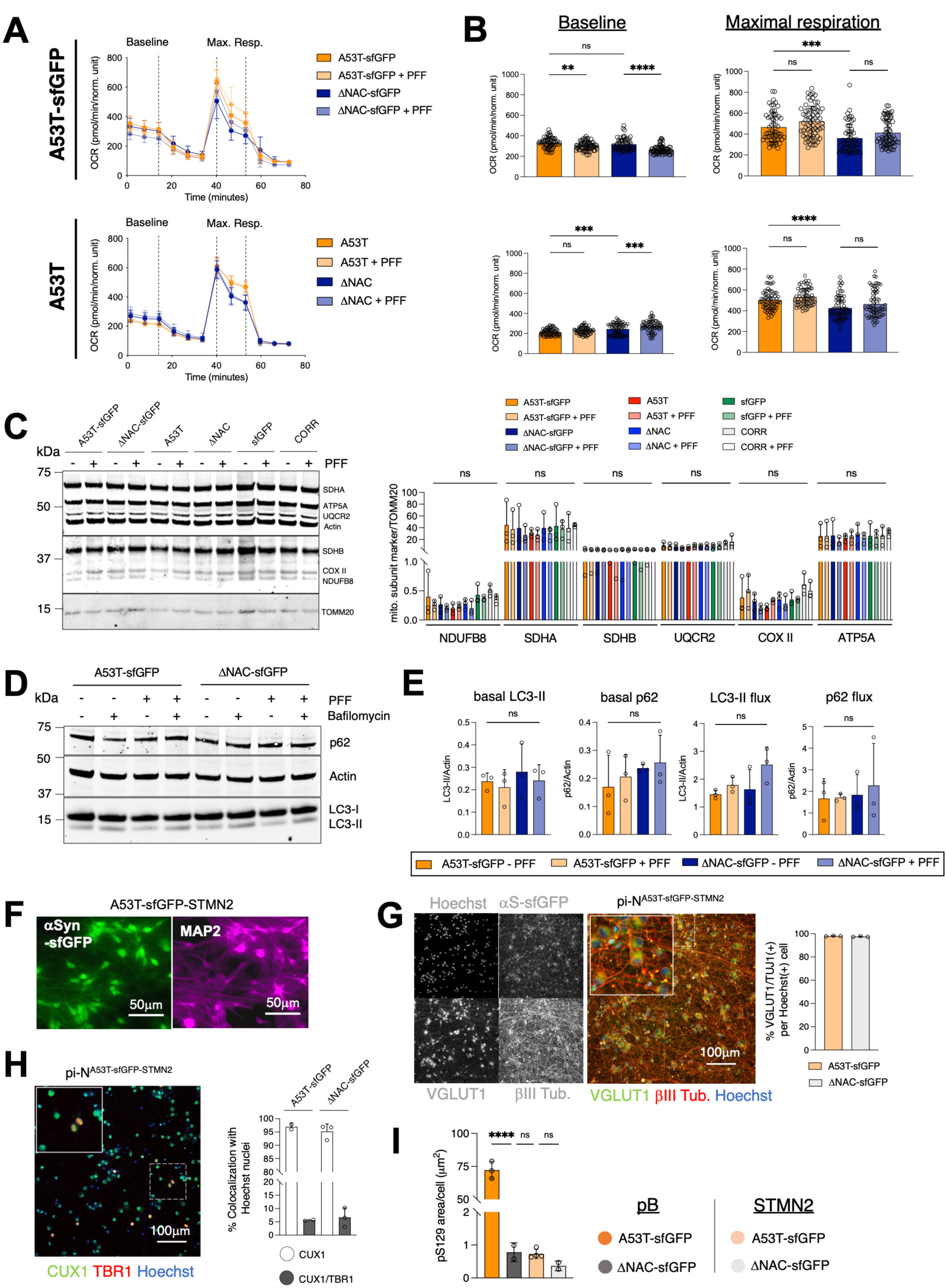
Phenotypic characterization of mitochondrial and lysosomal function in seeded inclusionopathy model. (A) Seahorse assay in PFF-seeded and unseeded, sfGFP-tagged (top) and untagged (bottom) pi N^A53T-pB^ and pi-N^ΔNAC-pB^ induced neurons (DIV20). (B) Bar graphs comparing oxygen consumption rate (OCR) across the different conditions taken at baseline (0-18 min of the Seahorse assay) or at maximal respiration (taken at 36-54 min of the Seahorse assay). One-way ANOVA + Tukey’s multiple comparison test: ** p<0.01, *** p<0.001, **** p<0.0001. (C) Mitochondrial subunit assay in seeded inclusion model and control lines pi-N^sfGFP-pB^ and CORR/pi-N (DIV56). Right: Quantification of WB, normalized to TOMM20. (D) Lysosomal flux assay in PFF-seeded and unseeded, sfGFP-tagged and untagged pi-N^A53T-pB^ and pi-N^ΔNAC-pB^ induced neurons (DIV56). Neurons were treated with 100 nM bafilomycin for 12 h before harvesting and protein extracted. (E) Analysis of autophagic flux based on quantification of western blots probed for p62, actin, LC3-I and LC3-II in panel D. (F) SNCA-A53T-sfGFP transgene knock-in at *STMN2* locus shows expression in MAP2(+) neuronal cells. (G) IF and quantification of VGLUT1 and TUJ1 in pi-N^STMN2^ model confirms cortical glutamatergic neuron identity. (H) IF and quantification of pi-N^STMN2^ model for CUX and TBR1 confirms cortical neuronal identity (layer II/III). (I) Quantification of pS129 area from IF in the seeded inclusion models pi-N^A53T-sfGFP-pB^ and pi-N^A53T-sfGFP-STMN2^ compared to control lines pi-N^ΔNAC-sfGFP-pB^ and pi-N^ΔNAC-sfGFP-STMN2^.

**Supplementary Figure 4:**
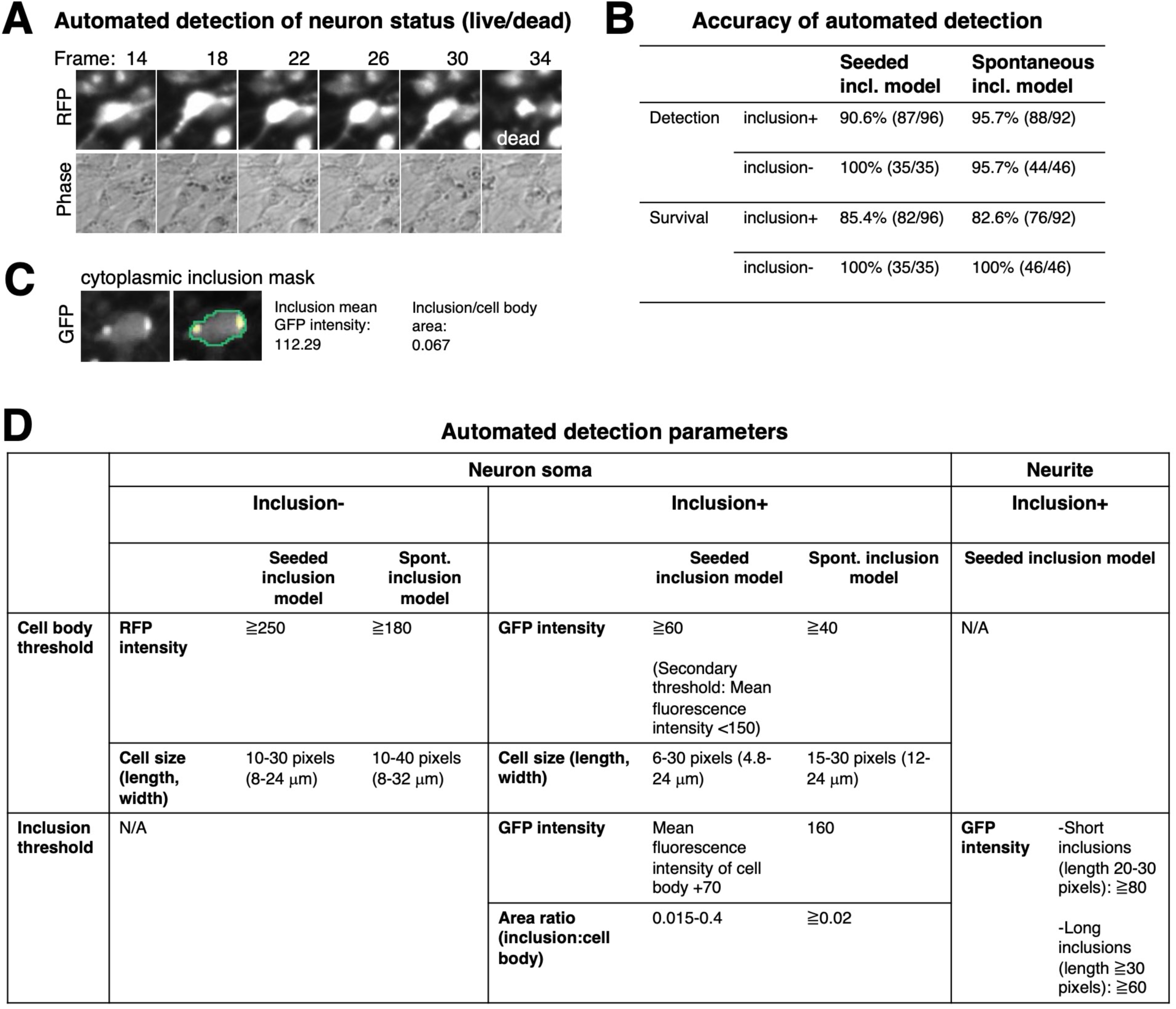
Longitudinal quantitative single-cell tracking of inclusionopathy models. (A) Example of automated detection of an RFP(+) neuron and automated detection of its death at frame 34. (B) Accuracy of automated detection and survival (live/dead status identification) of inclusion(+) and inclusion(−) neurons in seeded and spontaneous inclusion models. Denominator in parenthesis indicates number of neurons examined when evaluating accuracy of the algorithm. Detection indicates that neuron was identified correctly as a live, trackable cell. Survival indicates that the neuron was identified as dead at the correct frame. (C) Example of cytoplasmic inclusion mask detecting inclusions in PFF-seeded pi-N^A53T-sfGFP-pB^ neurons and algorithm output. (D) Thresholds for cell and inclusion automated detection. Neurons that meet the cell body thresholds go through a sorting step with secondary thresholds to eliminate dying, bright GFP(+) cells that were incorrectly identified as inclusion(+) cells.

**Supplementary Figure 5:**
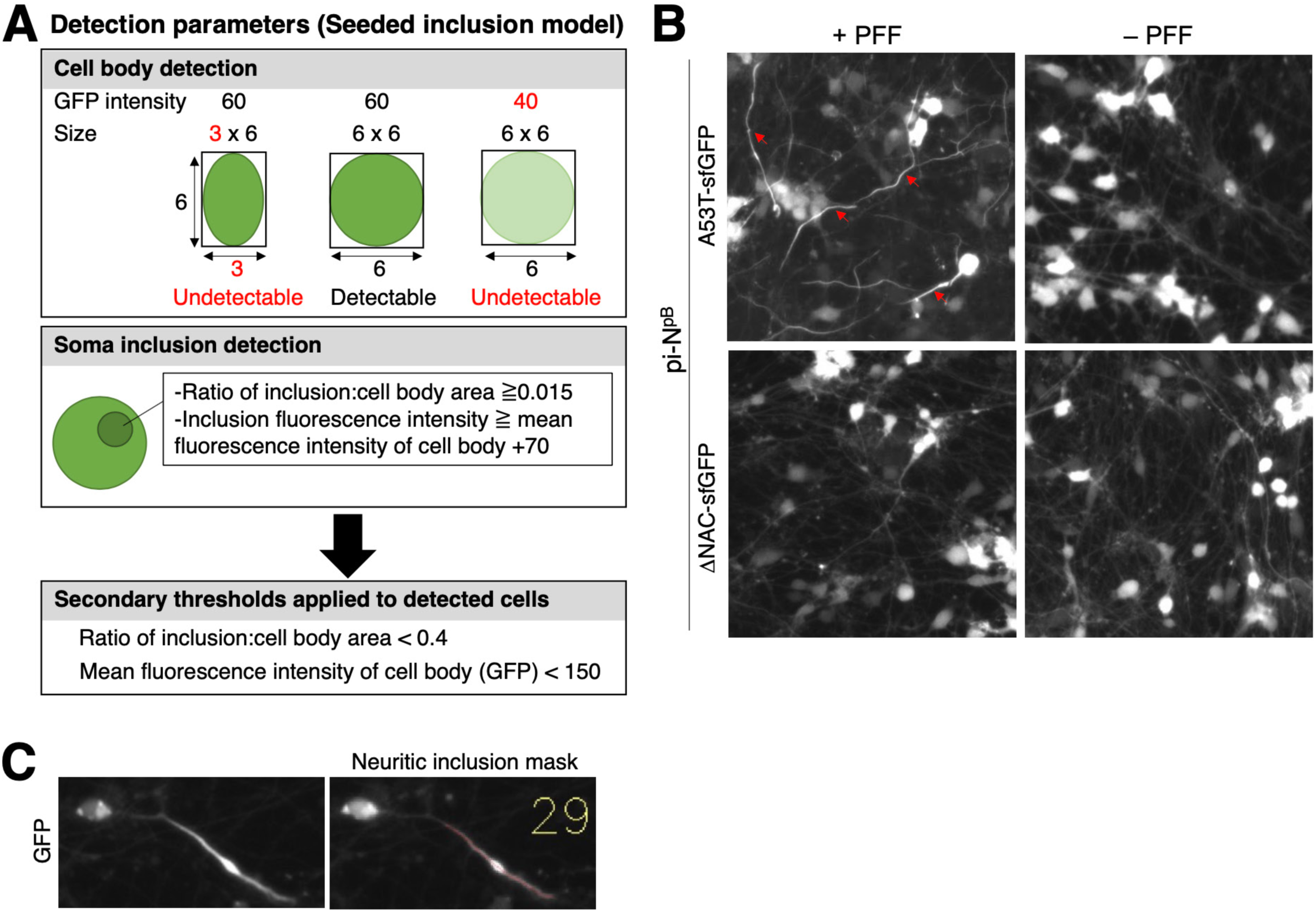
Longitudinal quantitative single-cell tracking of inclusionopathy models. (A) Diagram of detection parameters in inclusion survival tracking algorithm for seeded inclusionopathy model. (B) Biostation images of seeded and unseeded pi-N^A53T-sfGFP-pB^ or pi-N^ΔNAC-sfGFP-pB^ showing the abundance of neuritic inclusions in PFF-seeded pi-N^A53T-sfGFP-pB^ (red arrows). (C) Example of neuritic inclusion mask generated by the inclusion survival tracking algorithm.

**Supplementary Figure 6:**
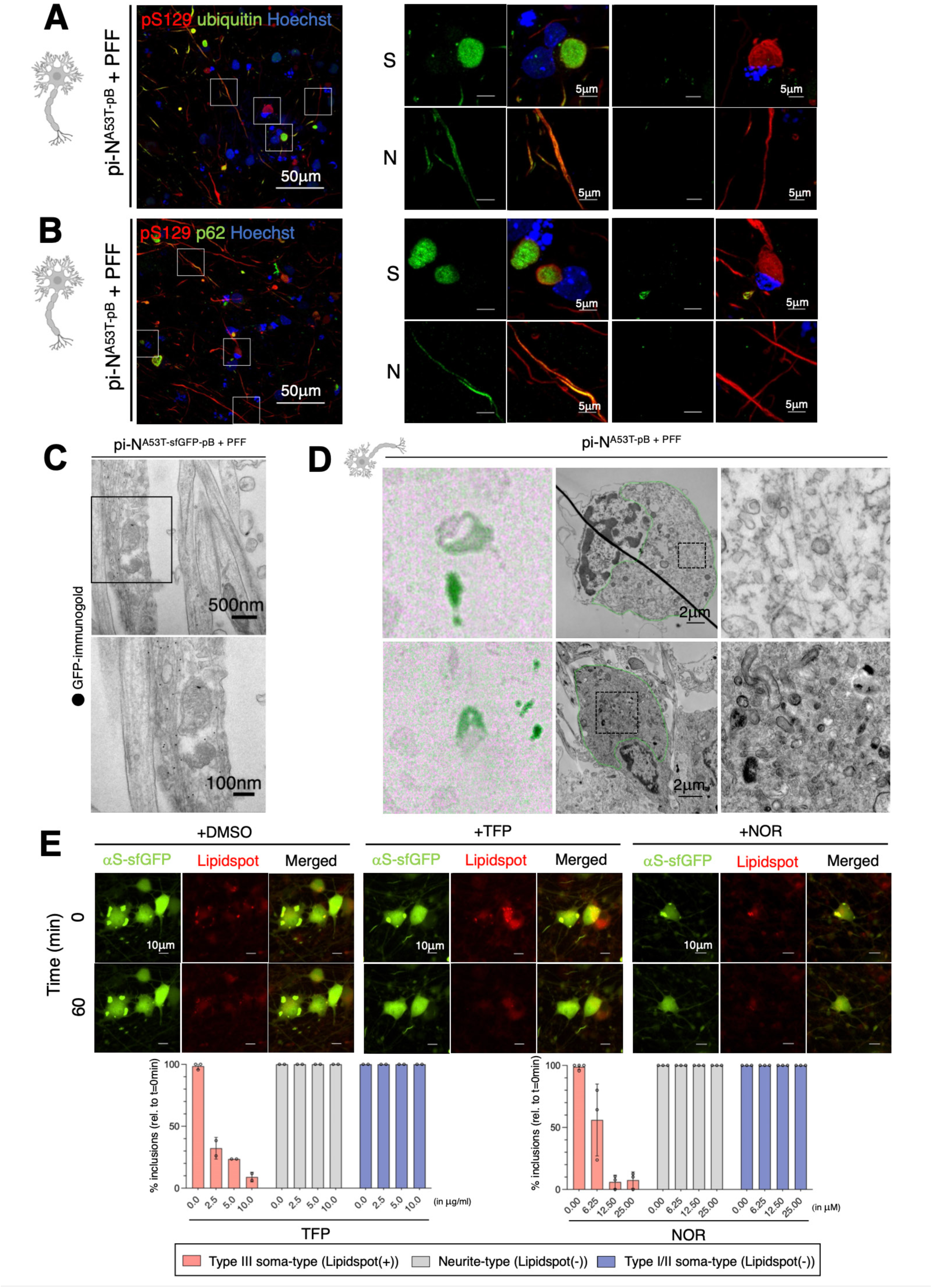
Examples of inclusion subtypes detected in seeded inclusionopathy models. (A) Left: Low magnification view showing the prevalence of ubiquitin(+) pS129(+) inclusions in seeded untagged pi-N^A53T-pB^ model. Right panels show representative examples of soma-type or neurite-type inclusions and their co-staining with ubiquitin. (B) Same as (A), except immunostainings are with p62 and pS129. (C) Neuritic inclusion detected by GFP-immunogold EM of tagged PFF-seeded pi-N^A53T-sfGFP-pB^ model. (D) CLEM of untagged seeded pi-N^A53T-pB^ model demonstrates that fibril-rich and lipid-rich αS(+) inclusions are also detected in the absence of sfGFP tag. (E) Previously reported PD drug candidates trifluoperazine and nortriptyline selectively dissolve Lipidspot(+) inclusions within min, but not Lipidspot(−) soma-type or neuritic inclusions. Top: High magnification images pre- (T=0min) and post-treatment (T=60min) of Lipidspot(+) inclusions with DMSO control, trifluoperazine or nortriptyline. Bottom: Quantification of percentage of Lipidspot(+), Lipidspot(−) neuritic or soma-type inclusions at T=60min post-treatment.

**Supplementary Figure 7:**
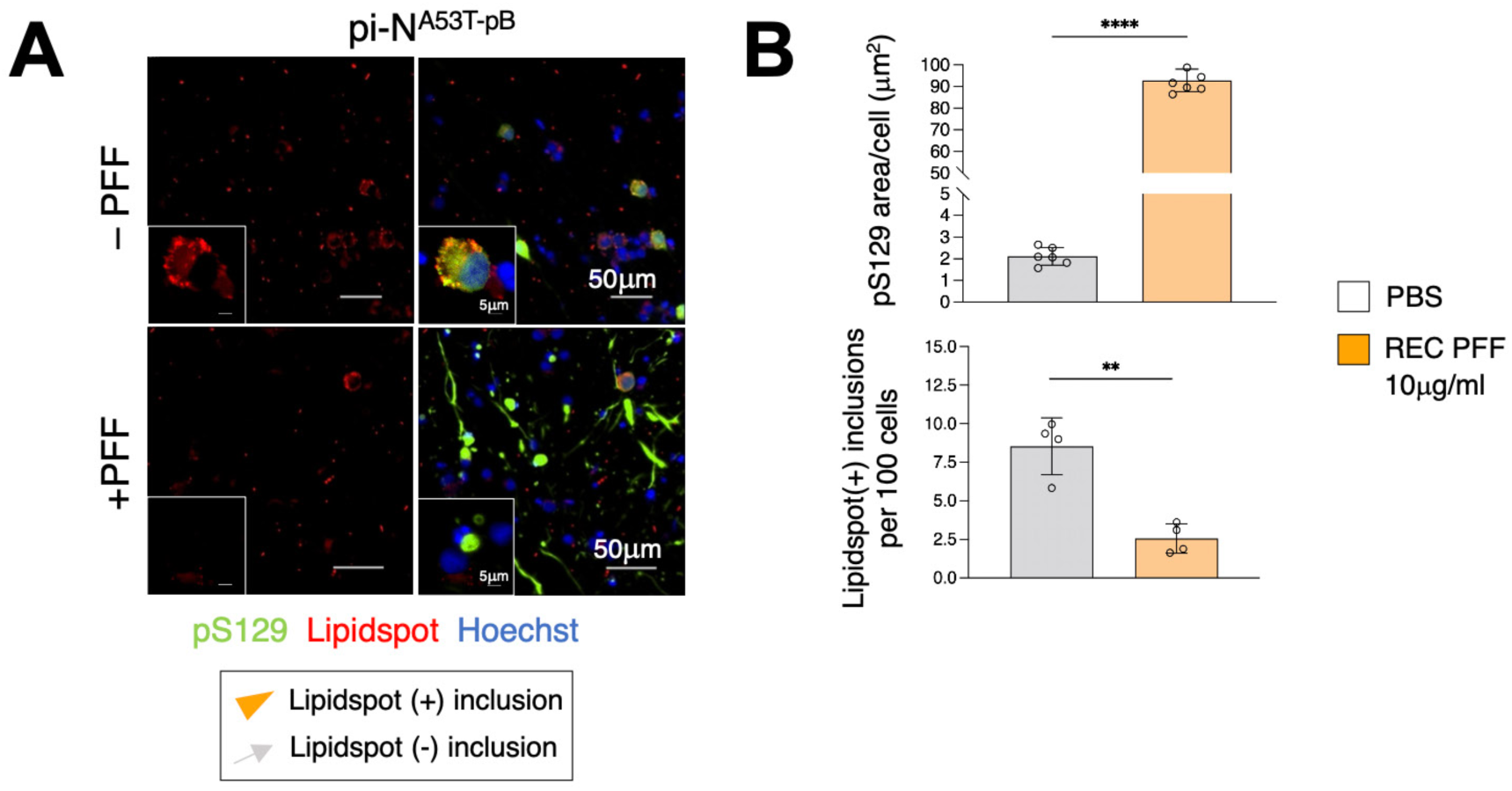
Frequency of Lipidspot(+) inclusions in seeded inclusionopathy model decreases upon seeding with PFFs. (A) Lipid-rich Lipidspot(+) inclusions are present in untagged seeded inclusion model (pi-N^A53T-pB^) and form in the absence of PFFs. Orange arrowheads, Lipidspot(+) inclusions; gray arrows, Lipidspot(−) inclusions. (B) Top: Frequency of inclusions increases upon seeding pi-N^A53T-pB^ neurons with recombinant PFFs, as detected by quantification of pS129 area/cell. Bottom: Concomitantly, the frequency of Lipidspot(+) inclusions decreases upon seeding with recombinant PFFs. One-way ANOVA + Tukey’s multiple comparison test: **, p<0.01; **** p<0.0001.

**Supplementary Figure 8:**
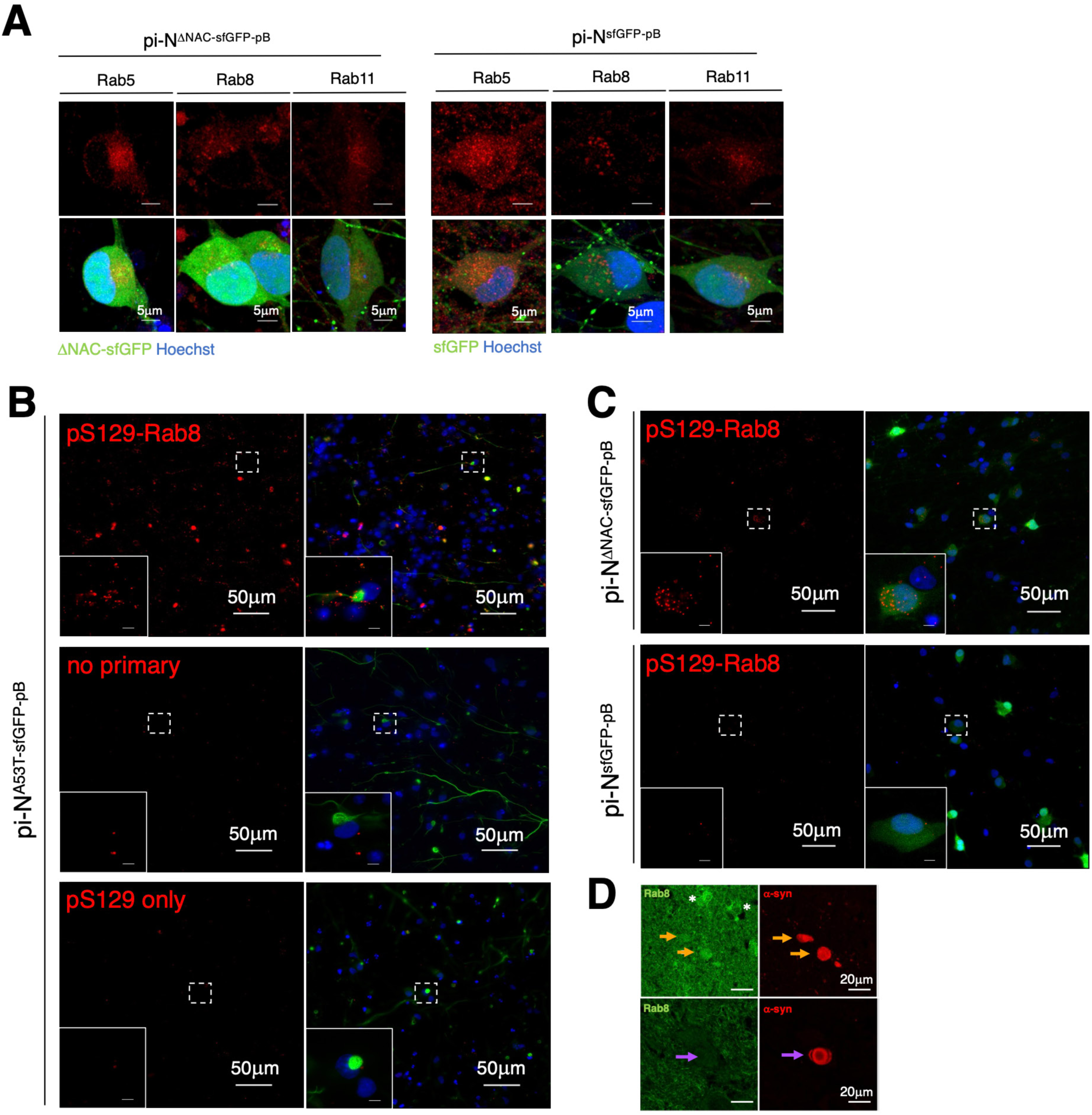
Co-localization of Rab markers with inclusion subtypes. (A) Rab5, Rab8 and Rab11 show diffuse, punctate staining pattern in pi-N^ΔNAC-sfGFP-pB^ and pi-N^sfGFP-pB^ control neurons. (B) Low-magnification views of Rab8-pS129 PLA micrographs. Signal is detected in seeded pi-N^A53T-sfGFP-pB^ when both pS129 and Rab8 primary antibodies are included in the assay, but not in the negative controls when no primary antibodies are included, or when only pS129 antibody is included. Right panels are merged images with sfGFP signal (indicating inclusions) and Hoechst nuclear stain. (C) Little or no Rab8-pS129 signal detected when PLA is performed in pi-N^ΔNAC-sfGFP-pB^ and pi-N^sfGFP-pB^ control neurons. (D) Rab8 signal is detected in an αS(+) inclusion that appears homogeneous (top row, orange arrow), but not in an inclusion with a dense core (bottom row, purple arrow) in familial E46K patient postmortem brain (substantia nigra pars compacta).

**Supplementary Figure 9:**
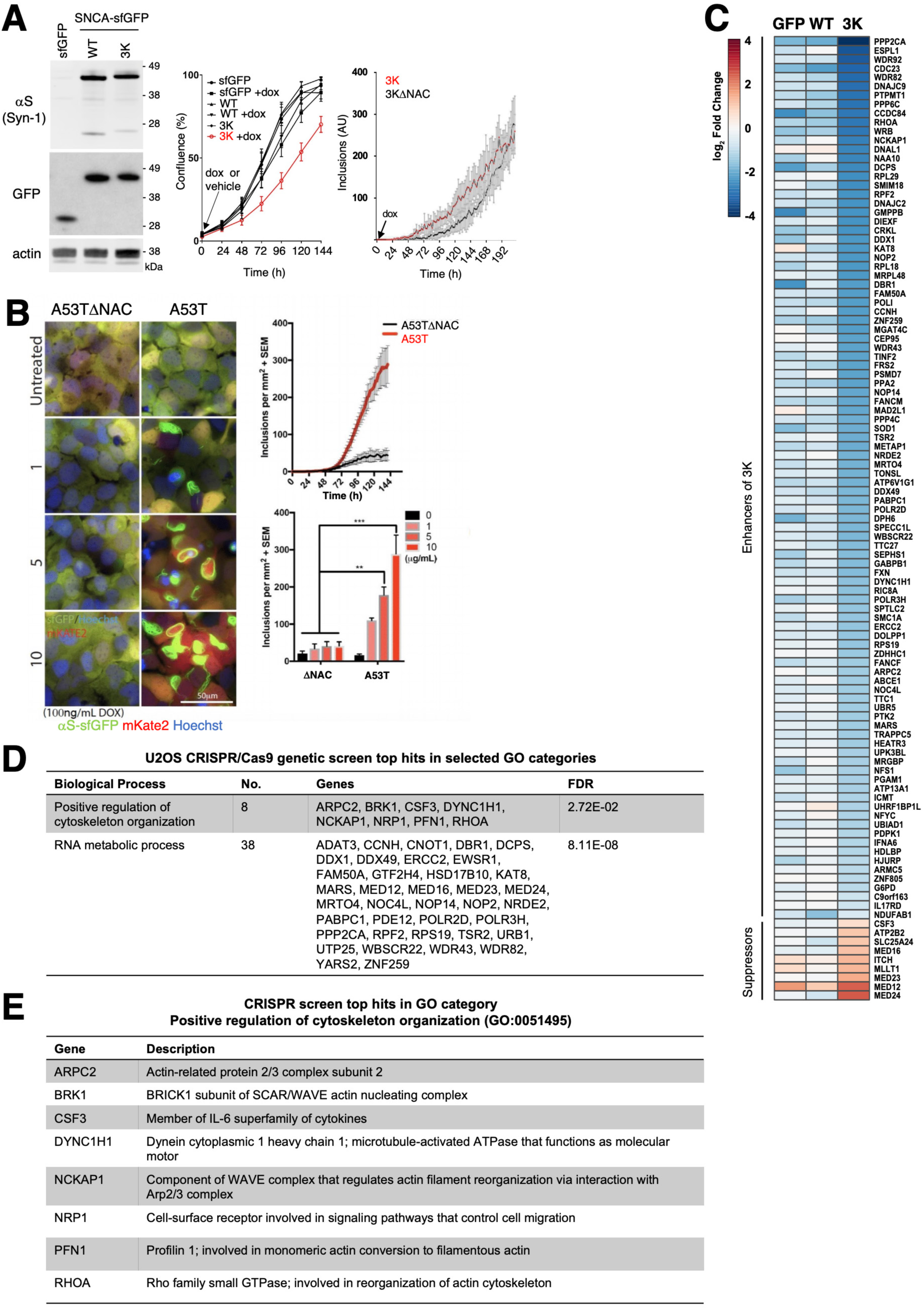
Convergence of CRISPR screen and MYTH on RhoA. (A) Left: Western blot showing pB-SNCA (3K-sfGFP, WT-sfGFP) or -sfGFP transgene expression upon doxycycline treatment in U2OS cells. Center: Incucyte cell culture confluence analysis of U2OS pB lines with and without doxycycline treatment. Expression of SNCA-3K-sfGFP results in cellular toxicity. Right: Inclusions in SNCA-3K-sfGFP are largely NAC-independent. (B) Left: Immunostaining of U2OS cells expressing pB-SNCA (A53T or A53TDNAC) induced with 100ng/mL doxycycline and treated with increasing amounts of recombinant A53T PFFs (µg/mL). Right, top: Inclusions in SNCA-A53T-sfGFP are NAC-dependent. Right, bottom: Increasing levels of inclusions are detected in SNCA-A53T-sfGFP with increasing amounts of recombinant PFFs seeded. (C) Heat map of U2OS CRISPR screen gene hits with highest log_2_ fold-change differential between 3K-sfGFP, WT-sfGFP and sfGFP indicating genetic modifiers of toxicity specific to 3K expression. (D) U2OS CRISPR/Cas9 genetic screen top hits in selected GO categories with statistically significant enrichment of gene hits. (E) CRISPR screen top hits in GO category “Positive regulation of cytoskeleton organization” (GO: 0051495).

**Supplementary Figure 10:**
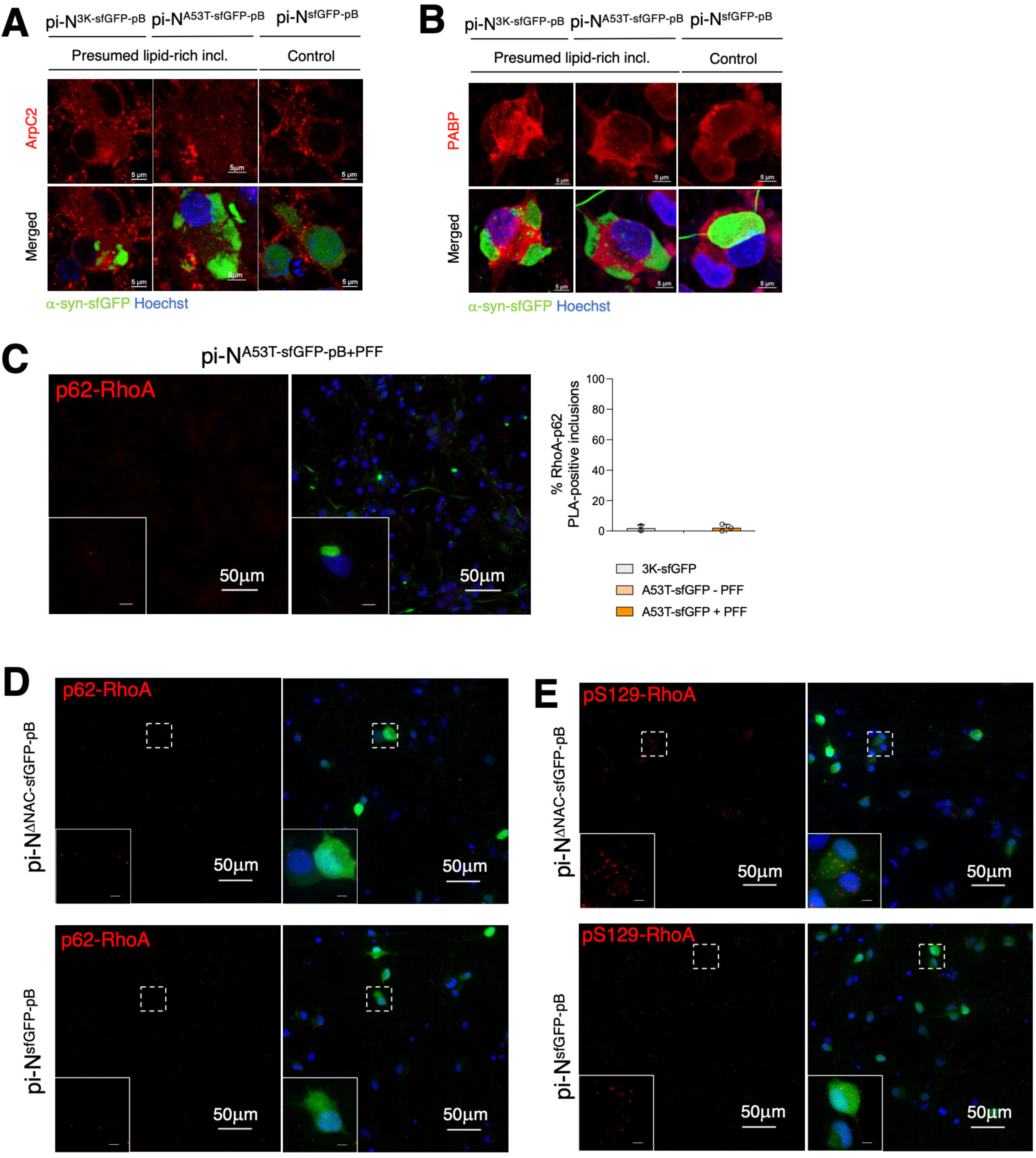
Co-localization of CRISPR screen hits with αS within inclusions. (A) ArpC2 immunostaining showing altered spatial pattern in pi-Ns with lipid-rich inclusions in both spontaneous and seeded inclusionopathy models, compared to sfGFP control neurons. Inclusion(+) neurons show diffuse ArpC2 staining, whereas neurons without inclusions exhibit larger ArpC2 puncta. (B) PABP immunostaining showing reduced signal within inclusions. (C) RhoA does not co-localize with p62(+) inclusions by PLA in seeded inclusionopathy model. Left: Panels with p62-RhoA PLA signal and merged A53T-sfGFP/Hoechst signal. Inset shows magnification of an inclusion(+) neuron. Center: Example of a neuron with detectable p62-RhoA PLA signal. Right: Quantification of αS-sfGFP(+) cytoplasmic inclusions that are positive for RhoA and p62 in spontaneous and seeded inclusion models (+/− PFFs). (D) p62-RhoA PLA signal in pi-N^ΔNAC-sfGFP-pB^ and pi-N^sfGFP-pB^ control neurons. (E) pS129-RhoA PLA signal in pi-N^ΔNAC-sfGFP-pB^ and pi-N^sfGFP-pB^ control neurons.

