## Supplementary material for "Rapid iPSC inclusionopathy models shed light on formation, consequence and molecular subtype of α-synuclein inclusions": Table S2

**Table S2: GO Panther Overrepresesentation Test for U2OS CRISPR genetic screen top 147 hits**

| GO biological process complete | Homo sapiens (REF) count (20589) | Count (147) | Expected | (over/under) | Fold enrichment | raw P-value | FDR |
|---|---|---|---|---|---|---|---|
| cellular nitrogen compound metabolic process (GO:0034641) | 3573 | 68 | 25.51 | + | 2.67 | 7.66E-16 | 1.20E-11 |
| nucleobase-containing compound metabolic process (GO:0006139) | 2825 | 57 | 20.17 | + | 2.83 | 5.82E-14 | 3.04E-10 |
| heterocycle metabolic process (GO:0046483) | 2999 | 59 | 21.41 | + | 2.76 | 4.94E-14 | 3.87E-10 |
| nucleic acid metabolic process (GO:0090304) | 2276 | 50 | 16.25 | + | 3.08 | 1.80E-13 | 5.63E-10 |
| cellular metabolic process (GO:0044237) | 6606 | 91 | 47.17 | + | 1.93 | 2.17E-13 | 5.68E-10 |
| nitrogen compound metabolic process (GO:0006807) | 6710 | 92 | 47.91 | + | 1.92 | 1.46E-13 | 5.71E-10 |
| cellular aromatic compound metabolic process (GO:0006725) | 3050 | 58 | 21.78 | + | 2.66 | 3.92E-13 | 6.82E-10 |
| macromolecule metabolic process (GO:0043170) | 5941 | 85 | 42.42 | + | 2 | 3.63E-13 | 7.12E-10 |
| gene expression (GO:0010467) | 2314 | 50 | 16.52 | + | 3.03 | 3.35E-13 | 7.50E-10 |
| organic cyclic compound metabolic process (GO:1901360) | 3292 | 60 | 23.5 | + | 2.55 | 7.64E-13 | 1.20E-09 |
| primary metabolic process (GO:0044238) | 7228 | 94 | 51.61 | + | 1.82 | 1.78E-12 | 2.54E-09 |
| organic substance metabolic process (GO:0071704) | 7697 | 97 | 54.95 | + | 1.77 | 3.66E-12 | 4.78E-09 |
| metabolic process (GO:0008152) | 8131 | 98 | 58.05 | + | 1.69 | 4.37E-11 | 5.27E-08 |
| RNA metabolic process (GO:0016070) | 1635 | 38 | 11.67 | + | 3.26 | 7.24E-11 | 8.11E-08 |
| ribosome biogenesis (GO:0042254) | 303 | 17 | 2.16 | + | 7.86 | 1.28E-10 | 1.34E-07 |
| cellular macromolecule metabolic process (GO:0044260) | 2518 | 47 | 17.98 | + | 2.61 | 3.39E-10 | 3.32E-07 |
| cellular component organization or biogenesis (GO:0071840) | 5727 | 77 | 40.89 | + | 1.88 | 4.43E-10 | 4.09E-07 |
| ribonucleoprotein complex biogenesis (GO:0022613) | 449 | 19 | 3.21 | + | 5.93 | 9.34E-10 | 8.13E-07 |
| cellular component biogenesis (GO:0044085) | 2633 | 47 | 18.8 | + | 2.5 | 1.47E-09 | 1.21E-06 |
| RNA processing (GO:0006396) | 868 | 25 | 6.2 | + | 4.03 | 3.88E-09 | 3.04E-06 |
| cellular process (GO:0009987) | 15044 | 136 | 107.41 | + | 1.27 | 5.29E-09 | 3.95E-06 |
| cellular biosynthetic process (GO:0044249) | 2464 | 44 | 17.59 | + | 2.5 | 6.02E-09 | 4.29E-06 |
| rRNA metabolic process (GO:0016072) | 254 | 14 | 1.81 | + | 7.72 | 7.45E-09 | 5.08E-06 |
| biosynthetic process (GO:0009058) | 2603 | 45 | 18.58 | + | 2.42 | 1.19E-08 | 7.78E-06 |
| rRNA processing (GO:0006364) | 223 | 13 | 1.59 | + | 8.17 | 1.40E-08 | 8.77E-06 |
| ncRNA metabolic process (GO:0034660) | 536 | 19 | 3.83 | + | 4.96 | 1.50E-08 | 9.06E-06 |
| ncRNA processing (GO:0034470) | 413 | 16 | 2.95 | + | 5.43 | 6.99E-08 | 4.06E-05 |
| organic substance biosynthetic process (GO:1901576) | 2534 | 42 | 18.09 | + | 2.32 | 1.93E-07 | 1.08E-04 |
| macromolecule biosynthetic process (GO:0009059) | 1487 | 30 | 10.62 | + | 2.83 | 2.38E-07 | 1.29E-04 |
| organelle organization (GO:0006996) | 3026 | 46 | 21.6 | + | 2.13 | 4.77E-07 | 2.49E-04 |
| cellular component organization (GO:0016043) | 5523 | 68 | 39.43 | + | 1.72 | 5.21E-07 | 2.64E-04 |
| maturation of SSU-rRNA from tricistronic rRNA transcript (SSU-rRNA, [illegible] | 39 | 6 | 0.28 | + | 21.55 | 7.43E-07 | 3.64E-04 |
| cellular nitrogen compound biosynthetic process (GO:0044271) | 1588 | 30 | 11.34 | + | 2.65 | 9.31E-07 | 4.42E-04 |
| ribosomal large subunit biogenesis (GO:0042273) | 76 | 7 | 0.54 | + | 12.9 | 2.06E-06 | 9.52E-04 |
| maturation of SSU-rRNA (GO:0030490) | 55 | 6 | 0.39 | + | 15.28 | 4.61E-06 | 2.07E-03 |
| protein-containing complex assembly (GO:0065003) | 1270 | 24 | 9.07 | + | 2.65 | 1.36E-05 | 5.91E-03 |
| cellular component assembly (GO:0022607) | 2394 | 36 | 17.09 | + | 2.11 | 1.40E-05 | 5.92E-03 |
| post-transcriptional regulation of gene expression (GO:0010608) | 498 | 14 | 3.56 | + | 3.94 | 1.74E-05 | 7.17E-03 |

| | | | | | | | |
|---|---|---|---|---|---|---|---|
| protein metabolic process (GO:0019538) | 3920 | 50 | 27.99 | + | 1.79 | 1.94E-05 | 7.81E-03 |
| organic cyclic compound biosynthetic process (GO:1901362) | 1216 | 23 | 8.68 | + | 2.65 | 2.07E-05 | 8.13E-03 |
| RNA catabolic process (GO:0006401) | 162 | 8 | 1.16 | + | 6.92 | 2.93E-05 | 1.09E-02 |
| protein-containing complex organization (GO:0043933) | 1423 | 25 | 10.16 | + | 2.46 | 2.93E-05 | 1.12E-02 |
| ribosomal small subunit biogenesis (GO:0042274) | 79 | 6 | 0.56 | + | 10.64 | 3.16E-05 | 1.15E-02 |
| heterocycle biosynthetic process (GO:0018130) | 1079 | 21 | 7.7 | + | 2.73 | 3.29E-05 | 1.17E-02 |
| negative regulation of chromosome organization (GO:2001251) | 82 | 6 | 0.59 | + | 10.25 | 3.85E-05 | 1.34E-02 |
| regulation of cellular amide metabolic process (GO:0034248) | 469 | 13 | 3.35 | + | 3.88 | 4.05E-05 | 1.38E-02 |
| DNA-templated transcription (GO:0006351) | 613 | 15 | 4.38 | + | 3.43 | 4.14E-05 | 1.38E-02 |
| nucleic acid-templated transcription (GO:0097659) | 614 | 15 | 4.38 | + | 3.42 | 4.21E-05 | 1.38E-02 |
| regulation of translation (GO:0006417) | 410 | 12 | 2.93 | + | 4.1 | 4.88E-05 | 1.56E-02 |
| RNA biosynthetic process (GO:0032774) | 624 | 15 | 4.46 | + | 3.37 | 5.03E-05 | 1.58E-02 |
| ribosomal large subunit assembly (GO:0000027) | 26 | 4 | 0.19 | + | 21.55 | 5.76E-05 | 1.74E-02 |
| positive regulation of actin filament polymerization (GO:0030838) | 53 | 5 | 0.38 | + | 13.21 | 5.66E-05 | 1.74E-02 |
| regulation of cellular macromolecule biosynthetic process (GO:200011 | 493 | 13 | 3.52 | + | 3.69 | 6.64E-05 | 1.89E-02 |
| macromolecule modification (GO:0043412) | 2883 | 39 | 20.58 | + | 1.89 | 6.63E-05 | 1.92E-02 |
| organonitrogen compound metabolic process (GO:1901564) | 5013 | 58 | 35.79 | + | 1.62 | 6.49E-05 | 1.92E-02 |
| positive regulation of organelle organization (GO:0010638) | 508 | 13 | 3.63 | + | 3.58 | 8.90E-05 | 2.49E-02 |
| positive regulation of transcription initiation by RNA polymerase II (GO | 59 | 5 | 0.42 | + | 11.87 | 9.11E-05 | 2.51E-02 |
| positive regulation of cytoskeleton organization (GO:0051495) | 195 | 8 | 1.39 | + | 5.75 | 1.02E-04 | 2.72E-02 |
| chromosome organization (GO:0051276) | 444 | 12 | 3.17 | + | 3.79 | 1.02E-04 | 2.76E-02 |
| ribosome assembly (GO:0042255) | 62 | 5 | 0.44 | + | 11.3 | 1.13E-04 | 2.92E-02 |
| aromatic compound biosynthetic process (GO:0019438) | 1089 | 20 | 7.78 | + | 2.57 | 1.13E-04 | 2.94E-02 |
| nuclear-transcribed mRNA catabolic process (GO:0000956) | 102 | 6 | 0.73 | + | 8.24 | 1.21E-04 | 3.01E-02 |
| nucleobase-containing compound biosynthetic process (GO:0034654) | 1007 | 19 | 7.19 | + | 2.64 | 1.20E-04 | 3.04E-02 |
| nucleic acid phosphodiester bond hydrolysis (GO:0090305) | 258 | 9 | 1.84 | + | 4.89 | 1.25E-04 | 3.06E-02 |
| regulation of organelle organization (GO:0033043) | 1189 | 21 | 8.49 | + | 2.47 | 1.28E-04 | 3.09E-02 |
| positive regulation of DNA-templated transcription initiation (GO:2000 | 66 | 5 | 0.47 | + | 10.61 | 1.49E-04 | 3.55E-02 |
| peptide metabolic process (GO:0006518) | 537 | 13 | 3.83 | + | 3.39 | 1.52E-04 | 3.56E-02 |
| regulation of transcription initiation by RNA polymerase II (GO:006026 | 67 | 5 | 0.48 | + | 10.45 | 1.60E-04 | 3.68E-02 |
| protein modification process (GO:0036211) | 2658 | 36 | 18.98 | + | 1.9 | 1.70E-04 | 3.86E-02 |
| DNA metabolic process (GO:0006259) | 794 | 16 | 5.67 | + | 2.82 | 2.09E-04 | 4.68E-02 |
| DNA-templated transcription initiation (GO:0006352) | 114 | 6 | 0.81 | + | 7.37 | 2.16E-04 | 4.76E-02 |
| negative regulation of organelle organization (GO:0010639) | 345 | 10 | 2.46 | + | 4.06 | 2.27E-04 | 4.95E-02 |

| | |
|---|---|
| Analysis Type: | PANTHER Overrepresentation Test (Released 20220712) |
| Annotation Version and Release Date: | GO Ontology database DOI: 10.5281/zenodo.6799722 Released 2022-07-01 |
| Analyzed List: | U2OS 147 hits alias.txt (Homo sapiens) |
| Reference List: | Homo sapiens (all genes in database) |
| Test Type: | FISHER |
| Correction: | FDR |
| Analysis run on 9/22/22 | |
