## Supplementary material for "Rapid iPSC inclusionopathy models shed light on formation, consequence and molecular subtype of α-synuclein inclusions": Table S3

**Table S3: U2OS CRISPR/Cas9 genetic screen top hits in GO category RNA metabolic process (GO:0016070)**

| Gene | Description |
|---|---|
| NOP14 | Nucleolar protein 14 |
| TSR2 | Pre-rRNA-processing protein TSR2 homolog |
| KAT8 | Histone acetyltransferase |
| POLR2D | DNA-directed RNA polymerase II subunit RPB4 |
| ADAT3 | Probable inactive tRNA-specific adenosine deaminase-like protein 3 |
| FAM50A | Protein FAM50A; may be involved in mRNA processing and splicing |
| GTF2H4 | General transcription factor IIH subunit 4 |
| EWSR1 | RNA-binding protein |
| WDR43 | WD repeat-containing protein 43 |
| MRTO4 | mRNA turnover protein 4 homolog |
| DDX1 | ATP-dependent RNA helicase |
| DDX49 | Probable ATP-dependent RNA helicase |
| ERCC2 | General transcription and DNA repair factor IIH helicase subunit XPD |
| MED24 | Mediator of RNA polymerase II transcription subunit 24 |
| DCPS | m7GpppX diphosphatase |
| YARS2 | Tyrosine--tRNA ligase, mitochondrial |
| CNOT1 | CCR4-NOT transcription complex subunit 1 |
| MED12 | Mediator of RNA polymerase II transcription subunit 12 |
| URB1 | Nucleolar pre-ribosomal-associated protein 1 |
| ZNF259 | Zinc finger protein ZPR1 |
| POLR3H | DNA-directed RNA polymerase III subunit RPC8 |
| CCNH | Cyclin-H kinase activator |
| MED23 | Mediator of RNA polymerase II transcription subunit 23 |
| HSD17B10 | 3-hydroxyacyl-CoA dehydrogenase type-2 |
| WBSCR22 | Probable 18S rRNA (guanine-N(7))-methyltransferase |
| NRDE2 | Nuclear exosome regulator |
| MED16 | Mediator of RNA polymerase II transcription subunit 16 |
| UTP25 | U3 small nucleolar RNA-associated protein 25 homolog |
| RPS19 | 40S ribosomal protein S19 |
| NOP2 | Probable 28S rRNA (cytosine(4447)-C(5))-methyltransferase |
| PDE12 | 2',5'-phosphodiesterase 12; mRNA polyadenylation factor |

| | |
|---|---|
| NOC4L | Nucleolar complex protein 4 homolog |
| P56192 | Methionine--tRNA ligase, cytoplasmic |
| PPP2CA | Serine/threonine-protein phosphatase 2A catalytic subunit alpha isoform |
| PABPC1 | Polyadenylate-binding protein 1 |
| RPF2 | Ribosome production factor 2 homolog |
| WDR82 | WD repeat-containing protein 82; component of SET1 histone H3-Lys4 methyltransferase complex |
| DBR1 | Lariat debranching enzyme |
