## Supplementary material for "Rapid iPSC inclusionopathy models shed light on formation, consequence and molecular subtype of α-synuclein inclusions": Table S1

**Table S1A: Non-transgenic iPSC lines**

| Source | iPSC line | Gender | Derivative line & clone | pB-NGN2 | pB-NFIB | Plasmid ID |
|---|---|---|---|---|---|---|
| **Synucleinopathy** | | | | | | |
| Familial PD | | | | | | |
| German Center for Neurodegenerative Diseas | *GBA* (L444P) | male | L444P (PD2-2 GBA 2/6) | pB-rtTA (#4)-NGN2-puro-SNAP | n/a | VK_1018 |
| German Center for Neurodegenerative Diseas | *GBA* (L444P) | male | CORR (PD2-2 GC2 GBA 2/7) | pB-rtTA (#4)-NGN2-puro-SNAP | n/a | VK_1018 |
| German Center for Neurodegenerative Diseas | *LRRK2* (2019S) | female | G2019S (LRRK2-GS-1, T4.6mut) | pB-rtTA (#4)-NGN2-puro-SNAP | n/a | VK_1018 |
| German Center for Neurodegenerative Diseas | *LRRK2* (2019S) | female | G2019S (LRRK2-GS-2, L2.1mut) | pB-rtTA (#4)-NGN2-puro-SNAP | n/a | VK_1018 |
| German Center for Neurodegenerative Diseas | *LRRK2* (2019S) | female | CORR/GS-1 | pB-rtTA (#4)-NGN2-puro-SNAP | n/a | VK_1018 |
| German Center for Neurodegenerative Diseas | *LRRK2* (2019S) | female | CORR/GS-2 | pB-rtTA (#4)-NGN2-puro-SNAP | n/a | VK_1018 |
| Yumanity Therapeutics/BWH | *SNCA* (A53>T) | male | A53T-1A | pB-rtTA (#4)-NGN2-puro-SNAP | n/a | VK_1018 |
| Yumanity Therapeutics/BWH | *SNCA* (A53>T) | male | A53T-1C | pB-rtTA (#4)-NGN2-puro-SNAP | n/a | VK_1018 |
| Yumanity Therapeutics/BWH | *SNCA* (A53>T) | male | A53T-1D | pB-rtTA (#4)-NGN2-puro-SNAP | n/a | VK_1018 |
| Yumanity Therapeutics/BWH | *SNCA* (A53>T) | male | CORR28 (A53T-1A) | pB-rtTA (#4)-NGN2-puro-SNAP | n/a | VK_1018 |
| Yumanity Therapeutics/BWH | *SNCA* (A53>T) | male | CORR63 (A53T-1C) | pB-rtTA (#4)-NGN2-puro-SNAP | n/a | VK_1018 |
| Coriell Institute | *SNCA* (triplication)-1 | female | clone C34139 (SNCA 4-copy) | pB-rtTA (#4)-NGN2-puro-SNAP | n/a | VK_1018 |
| Coriell Institute | *SNCA* (triplication)-1 | female | clone C6 (SNCA 2-copy) | pB-rtTA (#4)-NGN2-puro-SNAP | n/a | VK_1018 |
| Coriell Institute | *SNCA* (triplication)-1 | female | E2 (SNCA 0-copy) | pB-rtTA (#4)-NGN2-puro-SNAP | n/a | VK_1018 |
| Whitehead Institute | *SNCA* (triplication)-2 | male | clone G8 (SNCA 4-copy) | pB-rtTA (#4)-NGN2-puro-SNAP | n/a | VK_1018 |
| Whitehead Institute | *SNCA* (triplication)-1 | male | clone D9 (SNCA 4-copy) | pB-rtTA (#4)-NGN2-puro-SNAP | n/a | VK_1018 |
| Whitehead Institute | *SNCA* (triplication)-1 | male | clone F2 (SNCA 2-copy) | pB-rtTA (#4)-NGN2-puro-SNAP | n/a | VK_1018 |
| Whitehead Institute | *SNCA* (triplication)-1 | male | clone H6 (SNCA 2-copy) | pB-rtTA (#4)-NGN2-puro-SNAP | n/a | VK_1018 |
| Whitehead Institute | *SNCA* (triplication)-1 | male | clone E9 (SNCA 0-copy) | pB-rtTA (#4)-NGN2-puro-SNAP | n/a | VK_1018 |
| MSA | | | | | | |
| Mass General Brigham SCiN | MSA-1 | female | n/a | pB-rtTA (#4)-NGN2-puro-SNAP | n/a | VK_1018 |
| Mass General Brigham SCiN | MSA-5 | female | n/a | pB-rtTA (#4)-NGN2-puro-SNAP | n/a | VK_1018 |
| Mass General Brigham SCiN | MSA6 | male | n/a | n/a | n/a | n/a |
| Mass General Brigham SCiN | MSA-10 | male | n/a | n/a | n/a | n/a |
| Mass General Brigham SCiN | MSA-12 | male | n/a | n/a | n/a | n/a |
| Mass General Brigham SCiN | MSA-16 | female | n/a | n/a | n/a | n/a |
| Mass General Brigham SCiN | MSA-17 | female | n/a | n/a | n/a | n/a |
| Mass General Brigham SCiN | MSA-20 | male | n/a | pB-rtTA (#4)-NGN2-puro-SNAP | n/a | VK_1018 |
| Mass General Brigham SCiN | MSA-21 | male | n/a | n/a | pB-rtTA (#4)-NFIB-mKate2 | VK_1236 |
| Mass General Brigham SCiN | MSA-22 | male | n/a | n/a | n/a | n/a |
| Mass General Brigham SCiN | MSA-23 | female | n/a | n/a | pB-rtTA (#4)-NFIB-mKate2 | VK_1236 |
| Sporadic LBD | | | | | | |
| Mass General Brigham SCiN | PDD-1 | male | n/a | n/a | n/a | n/a |
| Mass General Brigham SCiN | DLB-2 | male | n/a | n/a | n/a | n/a |
| Mass General Brigham SCiN | PDD-2 | male | n/a | n/a | n/a | n/a |
| **Non-synucleinopathy** | | | | | | |
| Tauopathy | | | | | | |
| Mass General Brigham SCiN | Tau-1 | female | n/a | n/a | n/a | n/a |
| Mass General Brigham SCiN | PSP-1 | male | n/a | n/a | n/a | n/a |
| CAG expansion disease | | | | | | |
| Mass General Brigham SCiN | SCA2-2 | male | n/a | n/a | n/a | n/a |
| Mass General Brigham SCiN | SCA3-1 | female | n/a | pB-rtTA (#4)-NGN2-puro-SNAP | n/a | VK_1018 |
| Mass General Brigham SCiN | SCA3-3 | male | n/a | n/a | n/a | n/a |
| Mass General Brigham SCiN | SCA7-1 | female | n/a | n/a | n/a | n/a |
| Mass General Brigham SCiN | SCA7-2 | male | n/a | pB-rtTA (#4)-NGN2-puro-SNAP | n/a | VK_1018 |
| Healthy controls | | | | | | |
| Mass General Brigham SCiN | Ctrl-1 (C2) | female | n/a | pB-rtTA (#4)-NGN2-puro-SNAP | n/a | VK_1018 |
| Mass General Brigham SCiN | Ctrl-Tau-1 | male | n/a | n/a | n/a | n/a |
| Mass General Brigham SCiN | Ctrl-Tau-2 | male | n/a | n/a | n/a | n/a |
| Mass General Brigham SCiN | SCA7-Ctrl | male | n/a | pB-rtTA (#4)-NGN2-puro-SNAP | n/a | VK_1018 |
| Mass General Brigham SCiN | Ctrl-20 | male | n/a | pB-rtTA (#4)-NGN2-puro-SNAP | n/a | VK_1018 |
| The Translational Genomics Research Institut | TGEN/F01 036 | female | n/a | pB-rtTA (#4)-NGN2-puro | n/a | VK_1018 |
| The Translational Genomics Research Institut | TGEN/F02 037 | male | n/a | n/a | n/a | n/a |
| The Translational Genomics Research Institut | TGEN/F07 038 | female | n/a | n/a | n/a | n/a |
| The Translational Genomics Research Institut | TGEN/14-20 035 | female | n/a | n/a | n/a | n/a |
| The Translational Genomics Research Institut | TGEN/14-42 039 | male | n/a | n/a | n/a | n/a |
| The Translational Genomics Research Institut | TGEN/10-57 040 | male | n/a | n/a | n/a | n/a |
| hESC lines | | | | | | |
| WiCell Research Institute | H1 | male | n/a | n/a | pB-rtTA (#4)-NFIB-mKate2 | VK_1236 |
| WiCell Research Institute | H9 | female | n/a | pB-rtTA (#4)-NGN2-puro-SNAP | n/a | VK_1018 |

SCiN ... stem cells in neurodegeneration

**Table S1B: Transgenic lines**

| Source | iPSC/hESC line | Gender | Derivate clone | Protein of interest overexpressed | Fluorescent tag | Transgenic protein | Integration site | Transgenic construct | Piggybac construct | Plasmid ID |
|---|---|---|---|---|---|---|---|---|---|---|
| Yumanity Therapeutics/BWH | *SNCA* (A53>T) | male | CORR28 (A53T-1A) | aSyn (WT) | / | aSyn (WT) | pB random integration | pB-rtTA (#4)-aSyn(WT)-IRES-NGN2- | pB-rtTA (#4)-aSyn(WT)-IRES-NGN2- | VK_1203 |
| Whitehead Institute | WIBR-3 | female | WIBR3 38/16-1 hESC | aSyn (WT) | / | aSyn (WT) | AAVS1 locus | AAVS1:M2rrTA/ aSyn | pB-rtTA (#4)-NGN2-puro-SNAP | VK_1018 |
| Whitehead Institute | WIBR-3 | female | WIBR3 38/16-1 hESC | aSyn-G209A | / | aSyn-G209A | AAVS1 locus | AAVS1:M2rrTA/ aSyn-G209A | pB-rtTA (#4)-NGN2-puro-SNAP | VK_1018 |
| Whitehead Institute | WIBR-3 | female | WIBR3 38/16-1 hESC | aSyn-G209A | / | aSyn-G209A | AAVS1 locus | AAVS1:M2rrTA/ mKate2-2A-aSyn-G2 | pB-rtTA (#4)-NGN2-puro-SNAP | VK_1018 |
| Yumanity Therapeutics/BWH | *SNCA* (A53>T) | male | CORR28 (A53T-1A) | aSyn (A53T) | / | aSyn (A53T) | pB random integration | pB-rtTA (#4)-aSyn(A53T)-IRES-NGN | pB-rtTA (#4)-aSyn(A53T)-IRES-NGN | VK_1027 |
| Yumanity Therapeutics/BWH | *SNCA* (A53>T) | male | CORR28 (A53T-1A) | aSyn (A53T-DNAC) | / | aSyn (A53T-DNAC) | pB random integration | pB-rtTA (#4)-aSyn(A53T-DNAC)-IRE | pB-rtTA (#4)-aSyn(A53T-DNAC)-IRE | VK_1028 |
| Yumanity Therapeutics/BWH | *SNCA* (A53>T) | male | CORR28 (A53T-1A) | aSyn (3xK) | / | aSyn (3xK) | pB random integration | pB-rtTA (#4)-aSyn(3K)-IRES-NGN2-p | pB-rtTA (#4)-aSyn(3K)-IRES-NGN2-p | VK_1054 |
| Whitehead Institute | WIBR-3 | female | WIBR-3 (hESC) | aSyn (WT) | mKate2 | aSyn-mKate2 | AAVS1 locus | AAVS1:M2rrTA/ aSyn-mKate2 | pB-rtTA (#4)-NGN2-puro-SNAP | VK_1018 |
| Whitehead Institute | WIBR-3 | female | WIBR-3 (hESC) | aSyn (WT) | sfGFP | aSyn (WT)-sfGFP | AAVS1 locus | AAVS1:M2rrTA/ aSyn-sfGFP | pB-rtTA (#4)-NGN2-puro-SNAP | VK_1018 |
| Yumanity Therapeutics/BWH | *SNCA* (A53>T) | male | CORR28 (A53T-1A) | aSyn (WT) | sfGFP | aSyn (WT)-sfGFP | pB random integration | pB-rtTA (#4)-aSyn(WT)-sfGFP-IRES- | pB-rtTA (#4)-aSyn(WT)-sfGFP-IRES- | VK_1056 |
| Whitehead Institute | *SNCA* (triplication)-2 | male | clone D9 (SNCA 4-copy) | aSyn (WT) | sfGFP | aSyn (WT)-sfGFP | pB random integration | pB-rtTA (#4)-aSyn(WT)-sfGFP-IRES- | pB-rtTA (#4)-aSyn(WT)-sfGFP-IRES- | VK_1056 |
| Whitehead Institute | *SNCA* (triplication)-2 | male | clone E9 (SNCA 0-copy) | aSyn (WT) | sfGFP | aSyn (WT)-sfGFP | pB random integration | pB-rtTA (#4)-aSyn(WT)-sfGFP-IRES- | pB-rtTA (#4)-aSyn(WT)-sfGFP-IRES- | VK_1056 |
| Yumanity Therapeutics/BWH | *SNCA* (A53>T) | male | CORR28 (A53T-1A) | aSyn (A53T) | sfGFP | aSyn (A53T)-sfGFP | pB random integration | pB-rtTA (#4)-aSyn(A53T)-sfGFP-IRE | pB-rtTA (#4)-aSyn(A53T)-sfGFP-IRE | VK_1123 |
| Yumanity Therapeutics/BWH | *SNCA* (A53>T) | male | CORR28 (A53T-1A) | aSyn (A53T-DNAC) | sfGFP | aSyn (A53T-DNAC)-sfGFP | pB random integration | pB-rtTA (#4)-aSyn(A53T-DNAC)-sfG | pB-rtTA (#4)-aSyn(A53T-DNAC)-sfG | VK_1124 |
| WiCell Research Institute | H9 | female | 9 hESC STMN2-A53T-sfGFP clone 4- | aSyn (A53T) | sfGFP | aSyn (A53T)-sfGFP | STMN2 locus | STMN2-aSyn(A53T)-sfGFP | pB-rtTA (#4)-NGN2-puro-SNAP | VK_1018 |
| WiCell Research Institute | H9 | female | 9 hESC STMN2-A53T-sfGFP clone 6- | aSyn (A53T) | sfGFP | aSyn (A53T)-sfGFP | STMN2 locus | STMN2-aSyn(A53T)-sfGFP | pB-rtTA (#4)-NGN2-puro-SNAP | VK_1018 |
| WiCell Research Institute | H9 | female | ESC STMN2-A53T-DNAC-sfGFP clon | aSyn (A53T-DNAC) | sfGFP | aSyn (A53T-DNAC)-sfGFP | STMN2 locus | STMN2-aSyn(A53T-DNAC)-sfGFP | pB-rtTA (#4)-NGN2-puro-SNAP | VK_1018 |
| WiCell Research Institute | H9 | female | ESC STMN2-A53T-DNAC-sfGFP clon | aSyn (A53T-DNAC) | sfGFP | aSyn (A53T-DNAC)-sfGFP | STMN2 locus | STMN2-aSyn(A53T-DNAC)-sfGFP | pB-rtTA (#4)-NGN2-puro-SNAP | VK_1018 |
| Yumanity Therapeutics/BWH | *SNCA* (A53>T) | male | CORR28 (A53T-1A) | aSyn (3xK) | sfGFP | aSyn (3xK)-sfGFP | pB random integration | pB-rtTA (#4)-aSyn(3K)-sfGFP-IRES-N | pB-rtTA (#4)-aSyn(3K)-sfGFP-IRES-N | VK_1120 |
| Yumanity Therapeutics/BWH | MSA 1 | female | n/a | aSyn (WT) | eYFP | aSyn (WT)-eYFP | pB random integration | pB-rtTA (#4)-aSyn (WT)-eYFP-IRES- | pB-rtTA (#4)-aSyn (WT)-eYFP-IRES- | VK_1023 |
| Yumanity Therapeutics/BWH | MSA 5 | female | n/a | aSyn (WT) | eYFP | aSyn (WT)-eYFP | pB random integration | pB-rtTA (#4)-aSyn (WT)-eYFP-IRES- | pB-rtTA (#4)-aSyn (WT)-eYFP-IRES- | VK_1023 |
| Mass General Brigham SCiN | Ctrl-1 (C2) | female | C2B-F | aSyn (WT) | eYFP | aSyn (WT)-eYFP | pB random integration | pB-rtTA (#4)-aSyn (WT)-eYFP-IRES- | pB-rtTA (#4)-aSyn (WT)-eYFP-IRES- | VK_1023 |
| The Translational Genomics Research Institut | TGEN/F01 036 | female | F01 036 iPSC | aSyn (WT) | eYFP | aSyn (WT)-eYFP | pB random integration | pB-rtTA (#4)- aSyn (WT)-eYFP-IRES- | pB-rtTA (#4)- aSyn (WT)-eYFP-IRES- | VK_1023 |
| Whitehead Institute | WIBR-3 | female | WIBR3 38/16-1 hESC | tau | / | tau | AAVS1 locus | AAVS1:M2rrTA/ Tau | pB-rtTA (#4)-NGN2-puro-SNAP | VK_1018 |
| Whitehead Institute | WIBR-3 | female | WIBR3 38/16-1 hESC | tau (E14) | / | tau (E14) | AAVS1 locus | AAVS1:M2rrTA/ TauE14 | pB-rtTA (#4)-NGN2-puro-SNAP | VK_1018 |
| Whitehead Institute | WIBR-3 | female | WIBR3 38/16-1 hESC | tau (P301L) | / | tau (P301L) | AAVS1 locus | AAVS1:M2rrTA/ TauP301L | pB-rtTA (#4)-NGN2-puro-SNAP | VK_1018 |
| Yumanity Therapeutics/BWH | *SNCA* (A53>T) | male | CORR28 (A53T-1A) | (sfGFP) | sfGFP | sfGFP | pB random integration | pB-rtTA (#4)-sfGFP-IRES-NGN2-puro | pB-rtTA (#4)-sfGFP-IRES-NGN2-puro | VK_1122 |
| WiCell Research Institute | H9 | female | n/a | (sfGFP) | sfGFP | sfGFP | pB random integration | pB-rtTA (#4)-sfGFP-IRES-NGN2-puro | pB-rtTA (#4)-sfGFP-IRES-NGN2-puro | VK_1122 |

**Table S1C: piggyBac plasmids**

| Plasmid ID | Piggybac construct | Acronym |
|---|---|---|
| VK_1018 | pB-rtTA (#4)-NGN2-puro-SNAP | pB-NGN2 |
| VK_1023 | pB-rtTA (#4)-SNCA(WT)-eYFP-IRES-Ngn2-puro | pB-SNCA(WT)-eYFP |
| VK_1027 | pB-rtTA (#4)-SNCA(A53T)-sfGFP-IRES-mKate2-puro | pB-A53T-IRES-mKate2 |
| VK_1028 | pB-rtTA (#4)-SNCA(A53T-∆NAC)-sfGFP-IRES-mKate2- | pB-SNCA(A53T-∆NAC)-sfGFP-IRES-mKate2 |
| VK_1054 | pB-rtTA (#4)-SNCA(3K)-sfGFP-IRES-mKate2-puro | pB-SNCA(3K)-sfGFP-IRES-mKate2 |
| VK_1056 | pB-rtTA (#4)-SNCA(WT)-sfGFP-IRES-mKate2-puro | pB-SNCA(WT)-sfGFP-IRES-mKate2 |
| VK_1074 | pB-rtTA (#4)-sfGFP-IRES-mKate2-puro | pB-sfGFP-IRES-mKate2 |
| VK_1116 | pB-rtTA (#4)-SNCA(3K-∆NAC)-sfGFP-IRES-mKate2-pu | pB-SNCA(3K-∆NAC)-IRES-mKate2 |
| VK_1120 | pB-rtTA (#4)-SNCA(3K)-sfGFP-IRES-NGN2-puro | pB-SNCA(3K)-sfGFP |
| VK_1122 | pB-rtTA (#4)-sfGFP-IRES-NGN2-puro | pB-sfGFP |
| VK_1123 | pB-rtTA (#4)-SNCA(A53T)-sfGFP-IRES-NGN2-puro | pB-SNCA(A53T)-sfGFP |
| VK_1124 | pB-rtTA (#4)-SNCA(A53T-∆NAC)-sfGFP-IRES-NGN2-p | pB-SNCA(A53T-∆NAC)-sfGFP |
| VK_1200 | pB-rtTA (#4)-SNCA(WT)-sfGFP-IRES-NGN2-puro | pB-SNCA(WT)-sfGFP |
| VK_1201 | pB-rtTA (#4)-SNCA(3K)-IRES-NGN2-puro | pB-SNCA(3K) |
| VK_1202 | pB-rtTA (#4)-SNCA(A53T)-IRES-NGN2-puro | pB-SNCA(A53T) |
| VK_1203 | pB-rtTA (#4)-SNCA(WT)-IRES-NGN2-puro | pB-SNCA(WT) |
| VK_1204 | pB-rtTA (#4)-SNCA(A53T-∆NAC)-IRES-NGN2-puro | pB-SNCA(A53T-∆NAC) |
| VK_1236 | pB-rtTA (#4)-NFIB-IRES-mKate2 | pB-NFIB |
